# Increased ultra-rare variant load in an isolated Scottish population impacts exonic and regulatory regions

**DOI:** 10.1101/809244

**Authors:** Mihail Halachev, Alison Meynert, Martin S Taylor, Veronique Vitart, Shona M Kerr, Lucija Klaric, SGP Consortium, Timothy J Aitman, Chris S Haley, James G Prendergast, Carys Pugh, David A Hume, Sarah E Harris, David C Liewald, Ian J Deary, Colin A Semple, James F Wilson

## Abstract

Human population isolates provide a snapshot of the impact of historical demographic processes on population genetics. Such data facilitate studies of the functional impact of rare sequence variants on biomedical phenotypes, as strong genetic drift can result in higher frequencies of variants that are otherwise rare. We present the first whole genome sequencing (WGS) study of the VIKING cohort, a representative collection of samples from the isolated Shetland population in northern Scotland, and explore how its genetic characteristics compare to a mainland Scottish population. Our analyses reveal the strong contributions played by the founder effect and genetic drift in shaping genomic variation in the VIKING cohort. About one tenth of all high-quality variants discovered are unique to the VIKING cohort or are seen at frequencies at least ten fold higher than in more cosmopolitan control populations. Multiple lines of evidence also suggest relaxation of purifying selection during the evolutionary history of the Shetland isolate. We demonstrate enrichment of ultra-rare VIKING variants in exonic regions and for the first time we also show that ultra-rare variants are enriched within regulatory regions, particularly promoters, suggesting that gene expression patterns may diverge relatively rapidly in human isolates.

**Author Summary:** Population isolates provide a valuable window on the roles of rare genetic variation in human phenotypes, as a result of their unusual evolutionary histories, that often lead to relatively high frequencies of variants that are exceptionally rare elsewhere. Such populations show increased levels of background relatedness among individuals and are often subject to stronger genetic drift, leading to a higher frequency of deleterious variants. Here, for the first time, we present whole genome sequencing data from the Shetland population in Northern Scotland, encompassing 500 individuals, and compare these genomes to the mainland Scottish population. As expected we find the imprint of Shetland population history in the Shetland genome, with strong evidence for founder effects and genetic drift, but we also discover a relaxation of selective constraint across the genome. These influences have combined to endow the Shetland genome with thousands of ultra-rare genetic variants, not observed previously in other populations. Surprisingly these variants are significantly enriched in functional regions including protein coding regions of genes and regulatory elements. Among regulatory regions, promoters are particularly enriched for ultra-rare variants, suggesting the potential for rapid divergence of gene expression in isolates.

## Introduction

Population isolates are subpopulations that originated from a small number of founders and subsequently remained relatively isolated for long periods of time due to geographical, cultural and social barriers. Such populations have been recognised to be of significant interest for some time [1], due to their unusual genetic characteristics. These include higher degrees of linkage-disequilibrium (LD), reduced haplotype complexity, increased numbers and extent of genomic regions within runs of homozygosity (ROH), high kinship, evidence for genetic drift, relatively high frequencies for otherwise rare variants, restricted allelic and locus heterogeneity [2–4]. Isolates are also subject to lower variation in environmental factors, tend to have better genealogical records, more uniform phenotyping and higher participation rates in studies [2]. Taken together, these genetic and other factors increase the power of gene mapping and association studies for both Mendelian and complex diseases and traits [5].

With the recent advances in high throughput sequencing (HTS) technologies, the traditional approach of investigating the genomic architecture of isolated populations via SNP genotyping arrays [6–13] has shifted towards using whole-exome sequencing (WES) [14–16] and low-coverage whole-genome sequencing (WGS) [17–21] to more recent high-coverage WGS studies [22–24]. The breadth and depth of high-coverage WGS provides unprecedented opportunities for interrogation of the effects of rare and ultra-rare variants genome wide, and may prove instrumental for addressing the “missing heritability” problem [25, 26].

For the first time our study used high-coverage WGS to compare the genomic landscapes of samples from an isolated population from the Shetland Islands to a more cosmopolitan mainland Scottish population. By investigating common and rare single nucleotide polymorphisms (SNPs) and short (up to 75bp) insertions/deletions (INDELs) in coding as well as in regulatory regions, we aimed to answer the following questions: *i*) is there any significant difference between the variant load observed in the two populations, *ii*) if so, what are the characteristics and the driving forces behind it and *iii*) which identified variants should be further examined for potential phenotype/trait associations?

The Shetland Islands lie scattered between ∼160-290 km (∼100-180 miles) north of the Scottish mainland and consist of a group of ∼100 islands, of which 16 are inhabited, with a population of ∼23,000 (Fig 1 in S1 File). First settled in the Neolithic period, ∼5400 years ago, the major demographic event in Shetland’s history was the arrival of the Norse Vikings about 800 CE. Shetland became part of the Jarldom of Orkney, centred on the archipelago to the south, until after over 500 years of Norse rule the islands were annexed by Scotland in 1472 [27]. Lowland Scots settled in Shetland both before and after this date; however, until the late 20^th^ century, the extreme geographic location in the north Atlantic served to isolate the population from further major immigration. In common with neighbouring areas, Shetland was variously affected by smallpox epidemics and famines over the centuries. Analyses of uniparental genetic systems reveal Shetland, like Orkney, to be a Norse-Scots hybrid population [28–30], with considerable genetic differentiation from the rest of the British Isles, reduced genetic diversity and longer stretches of linkage disequilibrium [31]. The presence of Norwegian ancestry in Shetland (23-28%) is further confirmed in a recent study based on high density autosomal SNP data [32].

**Fig 1.**
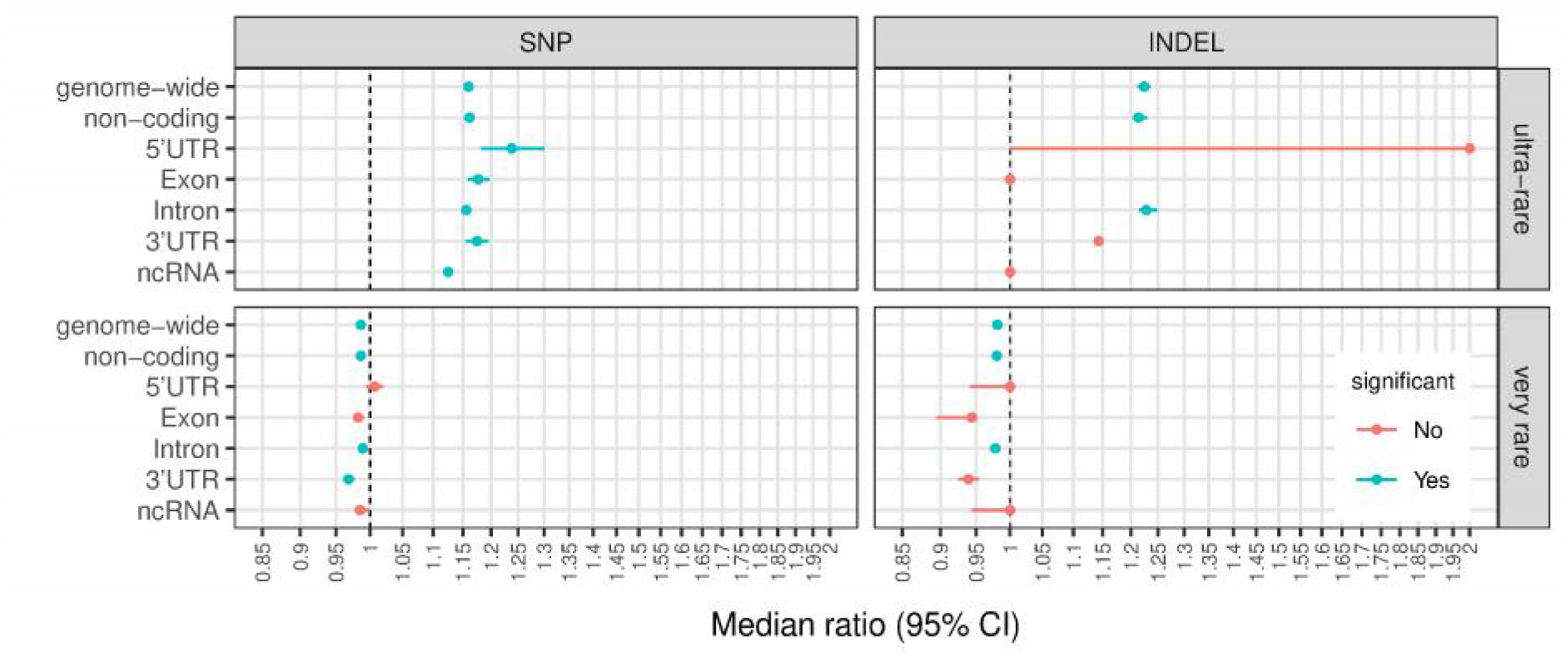
Significant differences in variant load in coding and related regions for ultra-rare (upper panel) and very rare (lower panel) variants. Circle dots represent the ratio of the median number of variants in a VIKING individual to the median number of variants in an LBC individual; whiskers are 95% CI based in 10,000 randomly selected LBC subsets (n = 269, with replacement). Significance: at least 95% of the 10,000 subsets have p-value ≤ 8×10^-4^ (Bonferroni corrected) and no overlap between the 95% CI for the LBC and the VIKING median values (for full results see Fig 4 in S1 File). The higher variance in the 5’UTR and lower variance in ncRNA regions could be explained by their relatively small sizes – 9.3Mb and 7.3Mb, respectively.

## Results

### Establishment of comparable Scottish isolate and mainland WGS datasets

A total of 2,122 participants of the VIKING Health Study – Shetland [33] were genotyped at ∼1 million SNP markers (using the Illumina HumanOmniExpressExome-8 v1.2 BeadChip) and 2,011 passed all quality control thresholds. All participants were selected to be over 18 years old and to have at least two grandparents born in the Shetland Isles (85% had four grandparents from Shetland, 10% had three and 5% had two grandparents born in the Shetland Isles). From the SNP genotyped cohort, 500 individuals were selected for whole-genome sequencing using the ANCHAP method [34] to most effectively represent the haplotypes present across the entire cohort. Unrelated individuals from the largest families were selected first, followed by those from smaller families, and finally some related individuals were selected to best represent the variation in the full cohort. The comparative population was 1369 individuals from the Lothian Birth Cohort (LBC) dataset [35–37] who were selected for WGS at the same facility as the VIKING samples. These are individuals born in 1921 or 1936 who attended Scottish schools and most took part in Scottish Mental Surveys in 1932 and 1947, respectively. Most were living in Edinburgh, Scotland (Fig 1 in S1 File) and the surrounding area (the Lothians) between 1999 and 2007.

The WGS data for the VIKING (median coverage 36.2x, range [27.1-40.2x], mean 36.1x, s.d. 2.0x) and LBC (median 37.3x, range [30.0-65.9x], mean 37.7x, s.d. 4.7x) cohorts were processed in an identical manner to identify and retain only high-quality SNP and INDEL variants (Materials and Methods). Overall concordance analysis between the SNP array data and WGS-derived genotypes for the Shetland cohort was performed to ensure there were no sample mix-ups by using the GenotypeConcordance tool from the GATK 3.6 toolkit [38] and was found to be 99.6%. We selected 269 unrelated (up to and including first cousin once removed and equivalents, pi_hat < 0.0625; for pi_hat definition see Materials & Methods, Sample selection) individuals from the Shetland cohort and 1156 unrelated individuals from the LBC. A total of 10,784,026 SNP sites and 1,082,383 INDEL sites were found in the 269 unrelated Shetland individuals (pi_hat mean = 0.0196, sd = 0.0164, median = 0.0269); the corresponding numbers for the 1156 unrelated LBC individuals (pi_hat mean = 0.0141, sd = 0.0130, median = 0.0188) are 21,152,042 SNPs and 2,065,442 INDELs. The two cohorts exhibited overall similar average numbers of high-quality variant alleles per sample (Table 1 in S2 File).

**Table 1A.**
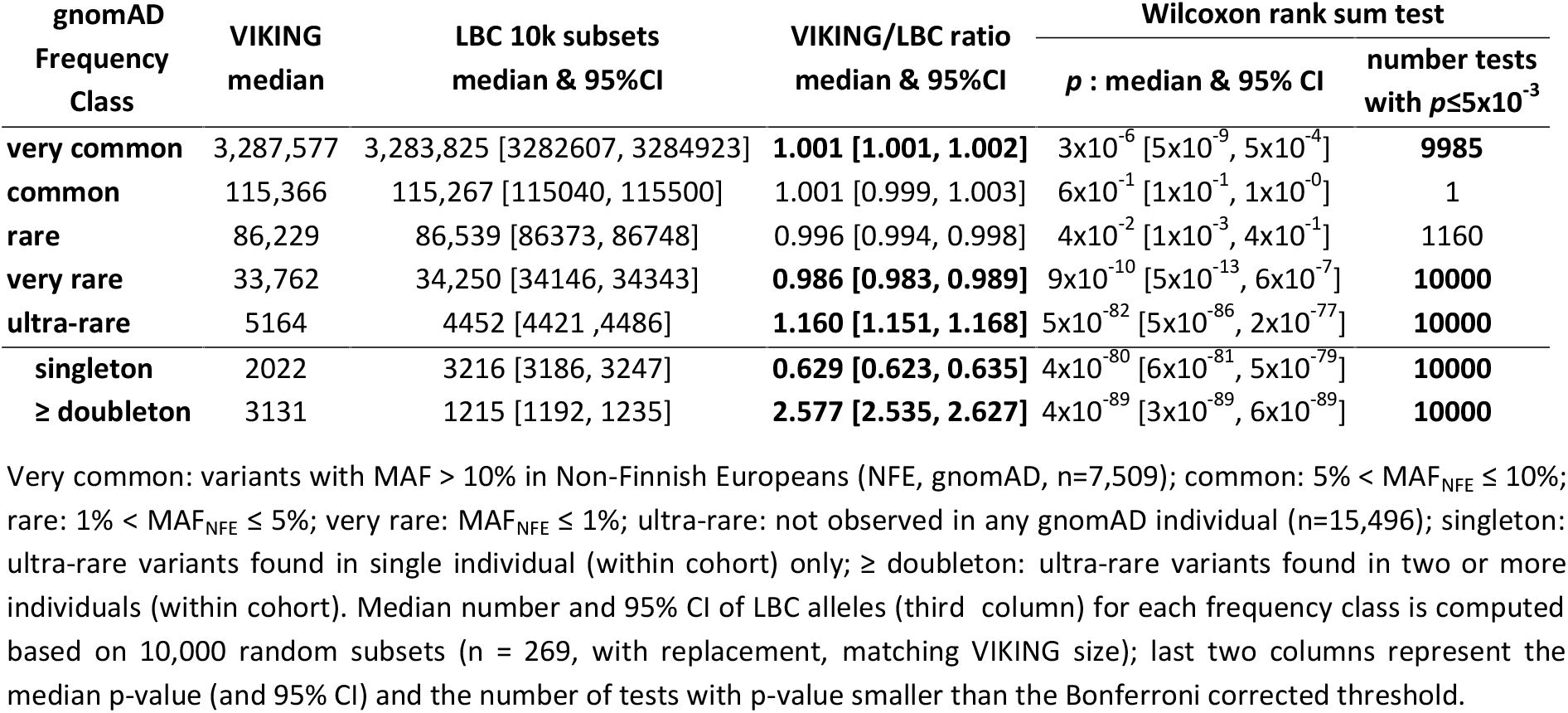
Genome-wide SNP load comparison in VIKING vs LBC (number of alleles per individual)

| gnomAD<br>Frequency<br>Class | VIKING<br>median | LBC 10k subsets<br>median & 95%CI | VIKING/LBC ratio<br>median & 95%CI | Wilcoxon rank sum test |  |
| --- | --- | --- | --- | --- | --- |
| | | | | p : median & 95% CI | number tests<br>with $p \leq 5 \times 10^{-3}$ |
| **very common** | 3,287,577 | 3,283,825 [3282607, 3284923] | **1.001 [1.001, 1.002]** | $3 \times 10^{-6}$ [$5 \times 10^{-9}$, $5 \times 10^{-4}$] | **9985** |
| **common** | 115,366 | 115,267 [115040, 115500] | 1.001 [0.999, 1.003] | $6 \times 10^{-1}$ [$1 \times 10^{-1}$, $1 \times 10^0$] | **1** |
| **rare** | 86,229 | 86,539 [86373, 86748] | 0.996 [0.994, 0.998] | $4 \times 10^{-2}$ [$1 \times 10^{-3}$, $4 \times 10^{-1}$] | **1160** |
| **very rare** | 33,762 | 34,250 [34146, 34343] | **0.986 [0.983, 0.989]** | $9 \times 10^{-10}$ [$5 \times 10^{-13}$, $6 \times 10^{-7}$] | **10000** |
| **ultra-rare** | 5164 | 4452 [4421, 4486] | **1.160 [1.151, 1.168]** | $5 \times 10^{-82}$ [$5 \times 10^{-86}$, $2 \times 10^{-77}$] | **10000** |
| **singleton** | 2022 | 3216 [3186, 3247] | **0.629 [0.623, 0.635]** | $4 \times 10^{-80}$ [$6 \times 10^{-81}$, $5 \times 10^{-79}$] | **10000** |
| **≥ doubleton** | 3131 | 1215 [1192, 1235] | **2.577 [2.535, 2.627]** | $4 \times 10^{-89}$ [$3 \times 10^{-89}$, $6 \times 10^{-89}$] | **10000** |

Very common: variants with MAF > 10% in Non-Finnish Europeans (NFE, gnomAD, n=7,509); common: 5% < MAF<sub>NFE</sub> ≤ 10%; rare: 1% < MAF<sub>NFE</sub> ≤ 5%; very rare: MAF<sub>NFE</sub> ≤ 1%; ultra-rare: not observed in any gnomAD individual (n=15,496); singleton: ultra-rare variants found in single individual (within cohort) only; ≥ doubleton: ultra-rare variants found in two or more individuals (within cohort). Median number and 95% CI of LBC alleles (third column) for each frequency class is computed based on 10,000 random subsets (n = 269, with replacement, matching VIKING size); last two columns represent the median p-value (and 95% CI) and the number of tests with p-value smaller than the Bonferroni corrected threshold.

**Table 1B.** Genome-wide INDEL load comparison in VIKING vs LBC (number of alleles per individual)

| gnomAD<br>Frequency<br>Class | VIKING<br>median | LBC 10k subsets<br>median & 95%CI | VIKING/LBC ratio<br>median & 95%CI | Wilcoxon rank sum test |  |
| --- | --- | --- | --- | --- | --- |
| | | | | p : median & 95% CI | number tests<br>with $p \leq 5 \times 10^{-3}$ |
| **very common** | 331,340 | 329,518 [329368, 329655] | **1.006 [1.005, 1.006]** | $5 \times 10^{-53}$ [$6 \times 10^{-58}$, $5 \times 10^{-48}$] | **10000** |
| **common** | 11,939 | 11,806 [11767, 11839] | **1.011 [1.008, 1.015]** | $3 \times 10^{-10}$ [$9 \times 10^{-14}$, $3 \times 10^{-7}$] | **10000** |
| **rare** | 8731 | 8657 [8630, 8689] | 1.009 [1.005, 1.012] | $2 \times 10^{-4}$ [$1 \times 10^{-6}$, $1 \times 10^{-2}$] | **9362** |
| **very rare** | 4001 | 4080 [4067, 4093] | **0.981 [0.978, 0.984]** | $8 \times 10^{-13}$ [$2 \times 10^{-16}$, $1 \times 10^{-9}$] | **10000** |
| **ultra-rare** | 503 | 411 [407, 415] | **1.224 [1.212, 1.236]** | $1 \times 10^{-82}$ [$5 \times 10^{-86}$, $1 \times 10^{-78}$] | **10000** |
| **singleton** | 183 | 284 [281, 287] | **0.644 [0.638, 0.651]** | $5 \times 10^{-77}$ [$4 \times 10^{-78}$, $7 \times 10^{-76}$] | **10000** |
| **≥ doubleton** | 324 | 124 [122, 127] | **2.613 [2.551, 2.656]** | $2 \times 10^{-89}$ [$2 \times 10^{-89}$, $3 \times 10^{-89}$] | **10000** |

A multidimensional scaling (MDS) analysis revealed that while similar, the two populations are genetically distinct from each other (Fig 2 in S1 File), and this was confirmed by a complementary admixture analysis (Fig 3 in S1 File). However, we adopted a conservative approach and did not exclude Shetland samples showing genotypes commonly found in LBC and *vice versa*. Such samples are representative of the fact that, although the Shetland population is isolated, there has been some gene flow to and from the capital city of Scotland and its surrounding area, where the LBC cohort were recruited. Inclusion of these individuals implies that any observed differences between the variant loads in the two cohorts will tend to be underestimated.

**Fig 2.**
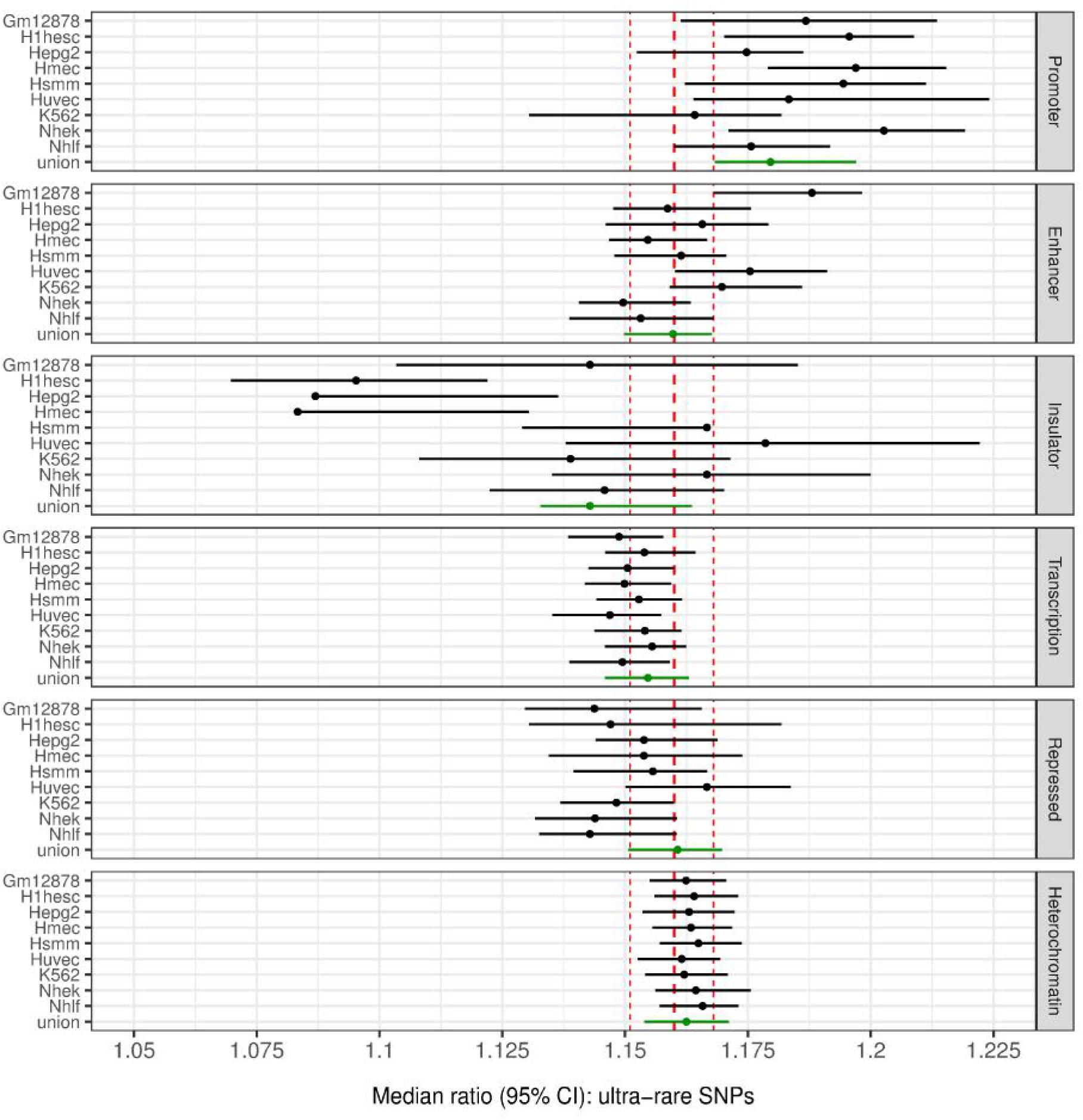
Ultra-rare SNP variant loads in functionally annotated non-coding regions. Circle dots represent the ratio of the median number of variants in a VIKING individual to the median number of variants in an LBC individual; whiskers are 95% CI based in 10,000 randomly selected LBC subsets (n = 269, with replacement). Significance: at least 95% of the 10,000 subsets have *p* ≤ 2×10^-4^ (Bonferroni corrected) and no overlap between the 95% CI for the LBC and the VIKING median values. The red vertical lines represent the median genome-wide load for ultra-rare SNPs and its 95% CI. The higher variance in the Insulator regions estimates could be explained by their relatively small size (17.4Mb). Gm12878: B-lymphoblastoid cells, H1hesc: embryonic stem cells, Hepg2: hepatocellular carcinoma cells, Hmec: mammary epithelial cells, Hsmm: skeletal muscle myoblasts, Huvec: umbilical vein endothelial cells, K562: erythrocytic leukemia cells, Nhek: normal epidermal keratinocytes, Nhlf: normal lung fibroblasts, union: an aggregated comparison between the two cohorts for this chromatin state by considering the union of state’s regions annotated in any of the 9 cell types.

**Fig 3.**
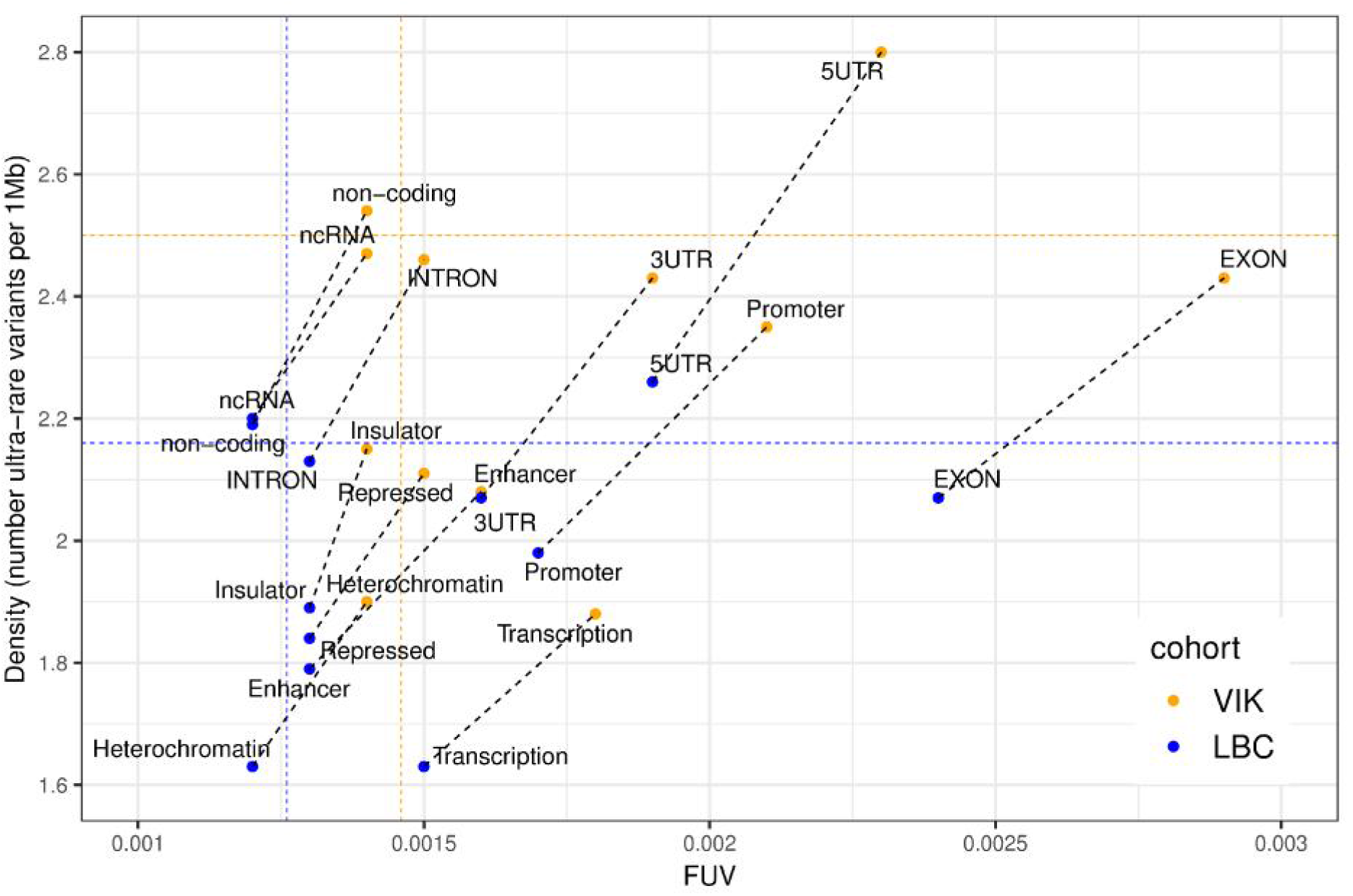
Distribution of ultra-rare SNPs in functional regions. Fraction of ultra-rare variants (FUV) = number of ultra-rare variants / (number of ultra-rare + known variants); Values for regulatory regions are computed as the average over the 9 cell types; non-coding = mappable genome – 5’UTR – exon – intron – 3’UTR – ncRNA; Coloured horizontal and vertical lines represent the genome-wide averages for the two cohorts. Dashed black lines represent the distribution shifts between LBC and VIKING for each of the considered genomic regions. A strictly vertical shift would indicate a proportional increase in the numbers of ultra-rare and known variants from LBC to VIKING, whereas a strictly horizontal shift (no change in the ultra-rare variant density between the two cohorts) would represent a decrease in the number of known variants in VIKING.

### The VIKING cohort is significantly enriched for ultra-rare SNP and INDEL variants genome-wide

To compare the genome-wide variant load in the two cohorts we stratified the variants found in the mappable regions of the 22 autosomal chromosomes based on their presence and MAF observed in the gnomAD genomes dataset (r2.0.1 [39]). We annotated variants as “ultra-rare” if they have not been observed in any individual in the full gnomAD genome dataset (n = 15,496); “very rare” for variants with MAF in Non-Finnish Europeans (NFE, n = 7,509) ≤ 1%, “rare” with 1% < MAF_NFE_ ≤ 5%, “common” with 5% < MAF_NFE_ ≤ 10%, and “very common” with MAF_NFE_ > 10%. To quantify the observed differences accurately for each frequency class, we bootstrapped the LBC data by generating 10,000 random subsets (with replacement) of size 269 individuals each to match the size of the VIKING dataset. For each of these subsets we counted the numbers of variants per individual in the VIKING and LBC cohorts and used the Wilcoxon rank sum test to evaluate the difference in distribution of number of variants between the two cohorts. To annotate the number of variants in a frequency class as significantly different (shown in bold, Table 1), we required at least 95% of the 10,000 subsets to have p-value ≤ 5×10^-3^ (Bonferroni corrected) and no overlap between the 95% CI for the LBC and VIKING median values.

Our results indicate that the VIKING samples are significantly enriched for ultra-rare SNPs (1.16 fold) and INDELs (1.22 fold) not observed in gnomAD (Table 1). Importantly, the observed enrichment is not driven by a greater individual-specific variation in Shetlanders; in fact, a VIKING individual carries less than two-thirds of the number of ultra-rare singleton variants compared to an LBC counterpart (see singleton versus ≥doubleton fractions of ultra-rare variants in Table 1).

To evaluate the potential effect of distant relatedness remaining in the chosen sets of 269 VIKING and 1156 LBC individuals on the ultra-rare variant load, we selected from them the 34 VIKING and 68 LBC individuals with no detectable relationships within each cohort (pi_hat = 0 within cohort). Using the discussed bootstrapping approach on these stricter subsets, we found that ultra-rare SNPs are enriched 1.14 fold (95% CI = [1.13, 1.16], *p* = 6.5×10^-11^, Wilcoxon rank sum test) and ultra-rare INDELs are enriched 1.20 fold (95% CI = [1.18, 1.23], *p* = 6.2×10^-11^) in the VIKING cohort; these values are very similar to the results obtained for the 269 VIKING and 1156 LBC sets (Table 1). Again, the overall enrichment is driven by the shared ultra-rare variants (i.e. ≥doubleton) - 3.03 fold ultra-rare SNP enrichment (*p* = 2.4×10^-12^) and 2.65 fold ultra-rare INDEL enrichment (*p* = 1.7×10^-12^) - whereas the two cohorts exhibit very similar levels of individual-specific ultra-rare variation and their difference is not significant.

These data suggest that genetic drift has increased the frequency of many ultra-rare variants in Shetland compared to those in Lothian. On average, a Shetland individual carries about 2.6 times more ultra-rare variants shared with at least one other Shetlander, compared to the ultra-rare variants shared within the Lothian individuals (Table 1). There is also a small but significant depletion of very rare known variants (MAF_NFE_ ≤ 1%) in VIKING, again due to the action of genetic drift whereby many rare variants are expected to be lost in the population.

### Elevated ultra-rare variant loads in the VIKING cohort at functional regions

Using data provided by Ensembl (GRCh37.p13, Ensembl Genes 92 [40]), we annotated the protein coding and related regions in the mappable sections of the 22 autosomal chromosomes as 5’UTR (a total length of 9.3M bases), exon (30Mb), intron (906Mb), 3’UTR (27.6Mb) and ncRNA regions (7.3Mb); the remaining 1.1Gb of the mappable regions in the reference human genome are labelled as “non-coding” regions (Materials and Methods). To make data from different regions comparable, we examined the number of variant alleles per megabase and used the same framework as for the genome-wide analysis to quantify the observed differences for each of the considered regions. The full results are available in Table 2 and Table 3 in S2 File and illustrated in Fig 4 in S1 File. As with the genome-wide level, in coding regions the two datasets are most divergent in terms of variant loads for ultra-rare and very rare variants; the results for these two regions are presented in Fig 1.

**Fig 4.**
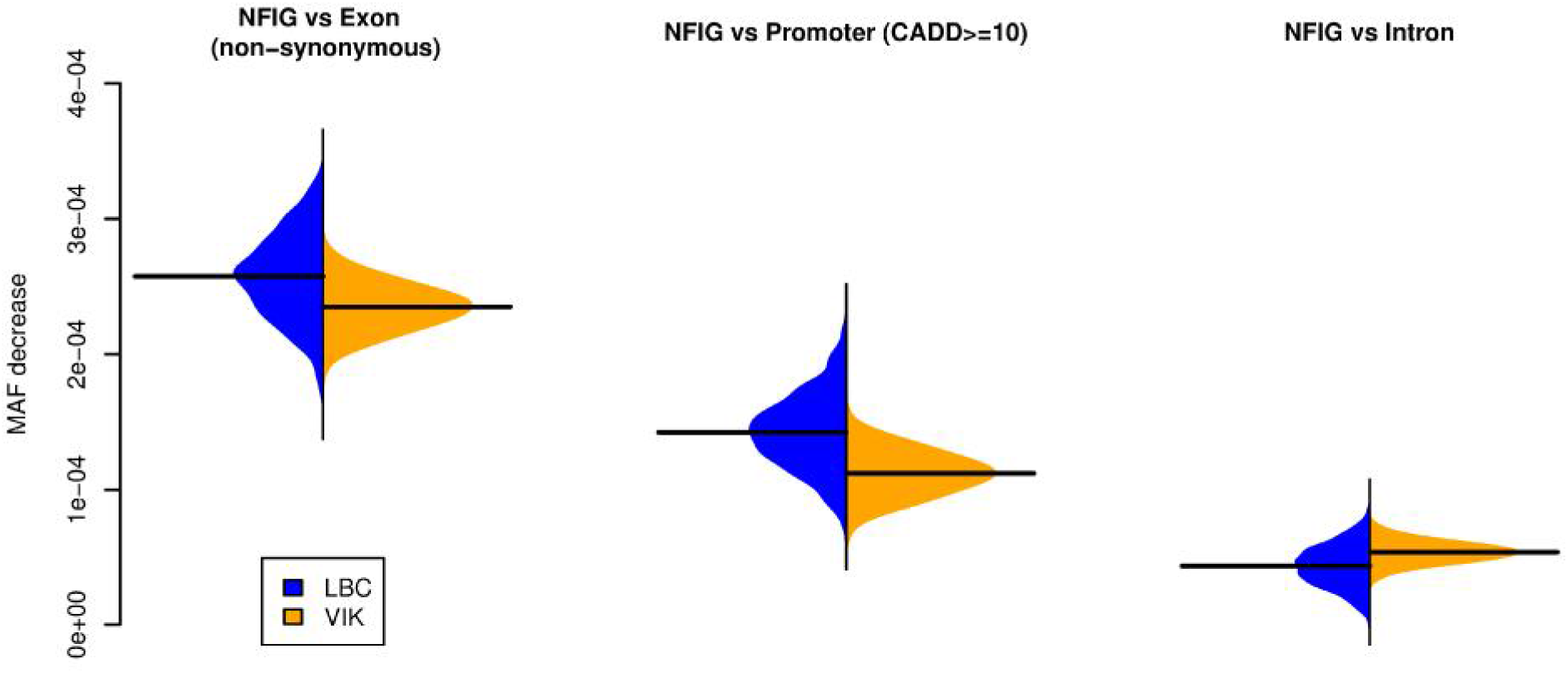
Allelic shift bias (ASB) suggests loss of constraint at VIKING exonic and promoter regions. MAF shifts for very rare SNPs (MAF_NFE_ ≤ 1%) between non-functional intergenic regions (NFIG), considered as baseline, and non-synonymous SNPs in exonic regions, SNPs with CADD score ≥ 10 in promoter regions and intronic SNPs, for each of the cohorts. These MAF differences are calculated using 1000 randomly selected LBC subsets of size 269 individuals (matching the VIKING size; with replacement) and considering only variants shared between the VIKING and the currently evaluated LBC subset, for which we computed the cohorts’ mean MAF in exonic, promoter, intronic and non-functional intergenic regions (see Fig 10 in S1 File). Black horizontal lines represent mean values. The differences in MAF shifts in the two cohorts are statically significant for all three comparisons (*p* < 2.2×10^-16^, one-sided Wilcoxon rank sum test).

**Table 2.** Variants observed in the VIKING cohort but not in gnomAD are often specific to Shetland.

| gnomAD<br>Frequency<br>Class | SNP enrichment |  |  |  | INDEL enrichment |  |  |  |
| --- | --- | --- | --- | --- | --- | --- | --- | --- |
|  | ≥ 2x | ≥ 5x | ≥ 10x | Shetland<br>specific | ≥ 2x | ≥ 5x | ≥ 10x | Shetland<br>specific |
| **very common & common** | ≤0.01% | n/a | n/a | n/a | ≤0.01% | n/a | n/a | n/a |
| **rare** | 0.80% | ≤0.01% | n/a | n/a | 0.72% | ≤0.01% | n/a | n/a |
| **very rare** | 31.64% | 16.01% | 10.49% | n/a | 28.99% | 14.26% | 9.35% | n/a |
| **ultra-rare** | 13.14% | 4.69% | 2.14% | 81.99% | 13.07% | 4.78% | 2.17% | 82.04% |

For variants seen in gnomAD, enrichment is computed against the maximum AF observed in LBC and gnomAD (all populations); for variants not found in gnomAD, enrichment and indigeneity is computed against LBC data.

**Table 3.** Tajima’s D captures demography and suggests relaxation of purifying selection in VIKING.

| Functional Region | VIK median [95% CI] | LBC median [95% CI] | Difference |
| --- | --- | --- | --- |
| Exon | -0.53 [-1.67, 1.24] | -0.85 [-0.86, -0.84] | 0.32 |
| 5'UTR | -0.27 [-1.56, 1.75] | -0.55 [-0.57, -0.53] | 0.28 |
| 3'UTR | -0.15 [-1.57, 1.63] | -0.48 [-0.50, -0.45] | 0.33 |
| ncRNA | 0.06 [-1.45, 2.22] | -0.24 [-0.26, -0.22] | 0.30 |
| Intron | 0.22 [-1.26, 1.22] | -0.19 [-0.20, -0.17] | 0.41 |
| non-coding | 0.38 [-1.04, 1.30] | -0.03 [-0.04, -0.01] | 0.41 |

VIKING Tajima's D values are based on aggregating the results for the 269 unrelated individuals over sliding genomic windows of size 1Mb (Materials and Methods). LBC results are based on aggregating the window medians for 100 random unrelated LBC subsets of size 269 individuals.

Our results show that VIKING samples are significantly enriched for ultra-rare SNPs in all coding related regions – including exonic regions – while potentially more damaging ultra-rare INDELs are restricted to non-coding and intronic regions. The observed exonic enrichment of ultra-rare SNPs is similar to the levels of enrichment seen genome-wide and in non-coding regions, demonstrating that exonic regions in the VIKING cohort have not been protected from the general accumulation of ultra-rare variation in spite of their functional importance. Indeed, the median enrichments seen in exons, 3’UTR and 5’UTR regions are somewhat higher than the genome-wide median enrichment.

We also annotated variants within predicted functional non-coding regions using the coordinates of 15 chromatin states generated for nine cell types by the NIH Roadmap Epigenomics Consortium [41], including promoters (average total length 39.2Mb over the 9 cell types), enhancers (130.5Mb), insulators (17.4Mb), transcribed (530.3Mb), repressed (130.5Mb) and heterochromatin (1.8Gb) regions (Materials and Methods). Using the same approach as for the genome-wide (Table 1) and coding analyses (Fig 1) to quantify variant loads for each of the chromatin states, we again found that the major difference between the two cohorts is for ultra-rare variant loads (Table 4 and Table 5 in S2 File). The observed significant enrichment of ultra-rare SNPs in all predicted regulatory regions was generally indistinguishable from the genome-wide level (Fig 2), suggesting that regulatory regions – similarly to the exonic regions – do not appear to be protected from ultra-rare SNP variants.

As for exonic regions, the median enrichment for promoters is generally somewhat higher than the genome-wide enrichment, particularly for predicted promoters active in H1 embryonic stem cells, HMEC primary mammary epithelial cells and NHEK epidermal keratinocyte cells (Fig 2).

The results for ultra-rare INDELs (Fig 5 in S1 File) are similar, but due to the small number of INDELs present in these regions, the conclusions are less robust. There is no significant difference in the regulatory regions for known SNPs in any of the 9 cell types (Table 6 in S2 File) and the enrichment for known INDELs in VIKING, although significant, is usually below 1% (Table 7 in S2 File).

### Strong founder effects and genetic drift in the VIKING cohort

A likely source of the observed enrichment of ultra-rare variants in the isolated Shetland population is the founder effect [42]. Among the variant sites found in VIKING samples but not present in gnomAD (i.e. the VIKING ultra-rare set) 707,600 SNPs (82%) and 63,549 INDELs (82%) are also absent from LBC (Table 2). These numbers represent 6.56% and 5.87% of all high-quality SNPs and INDELs identified in the VIKING cohort, respectively. Notably, approx. 0.8% of the VIKING SNPs and INDELs are ultra-rare, cohort-specific and seen in at least three VIKING individuals, compared to 0.35% of the LBC variants with the same characteristics, thus highlighting the potential role of founder effects, bottlenecks and restricted effective population size more generally in the VIKING cohort.

There is also evidence of genetic drift for VIKING variants shared only with LBC, as well as for variants shared with geographically more distant populations (Table 2). Among the VIKING ultra-rare variants (i.e. not seen in gnomAD), but present in LBC, there are 18,451 SNPs (2.14%) and 1,678 INDELs (2.17%) with allele frequency in VIKING at least ten times higher than in LBC. Considering the VIKING variants which are very rare in gnomAD Non-Finnish European population (MAF_NFE_ ≤ 1%), there are 359,275 SNPs (10.49%) and 31,713 (9.35%) INDELs with allele frequency in VIKING at least ten times higher than the maximum allele frequency observed in LBC and all gnomAD populations. Collectively, these enriched frequency variants represent 3.50% and 3.08% of all SNPs and INDELs identified in the VIKING cohort, respectively, highlighting the strength of genetic drift.

The above analyses reveal the extent of the contributions played by the founder effect and genetic drift in shaping the genomic variation in the isolated VIKING cohort. About one tenth of all high-quality variants discovered – 10.06% of the SNPs and 8.95% of the INDELs – are either unique to the VIKING cohort or seen at least ten times more frequently in it compared to cosmopolitan WGS populations (LBC and gnomAD).

Another line of evidence supporting the founder effect / genetic drift in the VIKING cohort is based on the analysis of the distribution of allele frequencies across polymorphic sites, also known as the site frequency spectrum (SFS) analysis (Materials and Methods). Our analysis is based on the high-quality variants discovered in the callable regions of the 22 autosomal chromosomes in the two cohorts of unrelated individuals, split to known variants (present in gnomAD at any frequency) and ultra-rare variants (not found in any gnomAD population).

The proportion of known variants (Fig 6 in S1 File) found as singletons was lower for VIKING compared to LBC: 19% (s.d. 6×10^-17^) versus 22% (s.d. 1×10^-16^) and 19% (s.d. 5×10^-3^) versus 21% (s.d. 3×10^-3^) for SNPs and INDELs, respectively, whereas the opposite is true for known variants found in two or more individuals. A similar trend was previously observed comparing the SFS of Finnish against non-Finnish Europeans [43], consistent with past founder effect(s).

The same trend, even amplified, is observed when comparing the SFS of the ultra-rare variants. VIKING individuals exhibit a much lower proportion of ultra-rare variants seen as singletons compared to LBC - 88% (s.d. 7×10^-3^) versus 98% (s.d. 5×10^-16^) and 86% (s.d. 7×10^-3^) versus 97% (s.d. 8×10^-16^) for SNPs and INDELs, respectively. Notably, 12% of the ultra-rare SNPs are shared by two or more among 50 randomly-chosen VIKING subjects compared to only 2% ultra-rare SNPs for LBC; 14% of the ultra-rare INDELs are shared by two or more VIKING subjects compared to 3% for LBC. These results support our finding of increased sharing of ultra-rare variants in VIKING compared to LBC (singleton versus ≥doubleton fractions in Table 1).

The roles played by founder effects and genetic drift in shaping the Shetland isolate were further evidenced by Tajima’s D [44] analysis (Materials and Methods) of the known SNPs (the variants present in the gnomAD dataset) in the six functional regions (Table 3). Tajima’s D values close to zero are considered as evidence for the neutral hypothesis, while negative values reflect high number of rare alleles due to population growth and/or purifying selection and positive Tajima’s D value indicate high number of alleles shared within the population [45].

As expected, for both cohorts we observe strongest purifying selection in exonic regions (the lowest Tajima’s D values), followed by 5’UTR, 3’UTR, ncRNA and intronic regions. The VIKING cohort exhibit higher Tajima’s D scores in all interrogated categories reflecting the specific demographic characteristics of this isolated population. Notably, the consistency of the Tajima’s D upwards shifts in VIKING compared to LBC (∼ 0.3 – 0.4), even in exonic regions, is suggestive of potential relaxation of purifying selection in the VIKING cohort, which we address in the next section.

Lastly, we compared the runs of homozygosity (ROH) identified in the two cohorts. ROHs were identified in VIKING and LBC individuals (Materials and Methods) and split into intermediate (length 0.5-2Mb) and long (≥ 2Mb) ROH (Fig 7 in S1 File). The total length of intermediate ROH in an individual is thought to reflect cryptic relatedness in populations, while the total length of long ROH usually shows large inter-individual variations that may reflect recent inbreeding patterns [3,46,47], or alternatively, a smaller effective population size. The observed correlation between the number of ROH and the total length is largely in accordance with data reported previously [48, 49]. To quantify potential differences between cohorts, similarly to the previous analyses, we generated 10,000 random LBC subsets from the data and for each subset we computed the medians, their ratio and the Wilcoxon p-value (Table 8 in S2 File). ROH with intermediate length were observed in all 269 VIKING and 1156 LBC samples, therefore we selected 10,000 LBC subsets of size 269 individuals (with replacement). We observed slight, but significant decrease in both the number and the total length of intermediate ROH in VIKING (VIKING/LBC median ratio ≈ 0.95, 95%CI ≈ [0.94, 0.96]). Long ROH were detected in 244 (91%) VIKING and 863 (75%) LBC unrelated individuals. Comparing the long ROH only in these individuals (subset size of 244 individuals, with replacement), we observed significant enrichment for both the number (ratio = 3.0 [1.5, 3.0], median *p* = 3×10^-22^) and the total length of ROH in VIKING (ratio = 2.31 [2.16, 2.93], median *p* = 2×10^-31^), consistent with increased parental kinship in the Shetland population.

### Evidence for relaxation of purifying selection in the VIKING cohort

Purifying (negative) selection is a powerful evolutionary mechanism of removing harmful genetic variation. It has been shown previously that isolated populations, due to their smaller effective population size, exhibit weaker purifying selection [19]. The strength of the purifying selection can be assessed by comparison of the distribution of rare derived variants across different functional categories. For example, analysis of the density and frequency of rare variants with derived allele frequency (DAF) < 0.5% in 2623 Icelandic whole genome sequences revealed that promoters had similar fraction of rare variants (FRV) and variant densities as UTRs, whereas enhancers had FRV and densities intermediate between UTRs on the one hand, and intronic, upstream or downstream regions on the other [22]. We performed similar, but more stringent, analyses of the VIKING and LBC data based on the ultra-rare SNPs discovered in the two cohorts and included data for protein coding and related regions (Fig 3). A comparison of the fraction of ultra-rare variants (FUV) and their densities in VIKING and LBC reveals that 5’UTR, exon and promoter regions show the most extreme shifts, driven by accumulation of ultra-rare variants at a higher rate compared to known variants in VIKING.

We sought formal evidence for the relaxation of purifying selection by examining the accumulation of extremely rare (i.e. singleton) variants predicted to have a loss of function (LOF) impact using the SVxy statistic (a comparison of the ratios of damaging to synonymous variants between isolate and other populations), which has previously been shown to identify weakened purifying selection in isolates [19]. As a baseline we used the Non-Finnish European (NFE) population in gnomAD (n = 7,509), extracting all exonic heterozygous SNPs (on the canonical transcript for each gene) found in a single NFE individual only. We filtered these singleton variants into two categories: i) LOF - stop gain, splice donor and splice acceptor variants, as well as missense variants with predicted deleterious CADD score ≥ 20 (the variant is predicted to be amongst the top 1% of deleterious variants in the human genome) [50]; and ii) synonymous (SYN) variants. There were 211,761 LOF and 158,077 SYN singleton alleles in NFE, such that the LOF/SYN ratio was 1.34. Similarly, from the VIKING and LBC ultra-rare variant sets we extracted the exonic singleton LOF and SYN variants, finding 23,787 LOF and 17,122 SYN singletons in the LBC cohort and 3,655 LOF and 2,501 SYN singletons in VIKING. The computed LOF/SYN ratios for the three cohorts correlate with the anticipated declining effective population size across these populations – from continent-wide Europeans (ratio = 1.34), to individuals born in the 1920-30s and living in Lothian, Scotland (ratio = 1.39), to the isolated Shetland population (ratio = 1.46).

For more rigorous evaluation of the potential relaxation of purifying selection in VIKING compared to LBC, we repeated the ultra-rare singleton comparison with an additional requirement of considering only genes for which there is at least one LOF or SYN variant observed in both cohorts [19]. This led to very similar results (4,030 genes, LBC_LOF/SYN_ = 1.40 and VIKING_LOF/SYN_ = 1.47), which indicates a 5.3% enrichment of ultra-rare singleton LOF SNP alleles in the VIKING cohort compared to LBC (*p* = 0.0387, one-sided Wilcoxon rank sum test; Fig 9 in S1 File). In [19], the authors studied 8 isolated populations and found a 1.2% enrichment of LOF alleles in an Orkney cohort (from the adjacent isolated northern Scottish archipelago) with respect to a cosmopolitan UK cohort, although the results are not readily comparable since their analysis was based on all (rather than only ultra-rare) singleton missense variants (regardless of their CADD score and not including nonsense and essential splice variants) as LOF variants and reporting mean instead of median values. Since the major difference in the variant load between VIKING and LBC is due to ultra-rare non-singleton variants (Table 1), we relaxed the singleton requirement above and performed the same analysis considering all ultra-rare variants in the two cohorts (5,365 genes with at least one LOF or SYN variant observed in both cohorts). The result shows a 9.4% enrichment of ultra-rare LOF SNP alleles in the VIKING cohort compared to LBC (*p* = 0.00064, one-sided Wilcoxon rank sum test).

### Allelic shift bias analysis supports widespread loss of selective constraint

LOF-based analyses can be applied only to exonic regions where variants can be split into two distinct categories based on their predicted impact. We developed a more general test, the allelic shift bias (ASB) test, which is designed to assess relaxation of selection in non-coding regions, based on the change in the allele frequency of variants within specific genomic regions across populations, as follows. We selected all SNPs in the VIKING and LBC cohorts found in the gnomAD genome dataset with MAF_NFE_ ≤ 1% in Non-Finnish Europeans. Given their low frequencies, these variants from the ancestral European population are likely to be enriched for SNPs that have been subject to purifying selection. We repeatedly (1000x) randomly selected 269 LBC individuals (matching the VIKING unrelated cohort size, with replacement) and selected MAF_NFE_ ≤ 1% variants shared between this LBC subset and the VIKING cohort. We then computed the mean MAF of such variants for each LBC subset and the VIKING cohort in exonic, promoter, intronic and non-functional intergenic (NFIG) regions (Fig 10 in S1 File). We also calculated the mean MAF of such variants for non-synonymous exonic variants and the predicted deleterious promoter variants (CADD score ≥ 10; predicted top 10% of the most deleterious variants genome-wide).

We estimated the strength of the purifying selection in each cohort as the difference between the mean MAF of the selected variants observed in the NFIG regions, where the effect of purifying selection is assumed to be negligible, and the mean MAF in regions assumed to be subject of active purifying selection. If purifying selection acts with the same strength in two populations there will be equivalent MAF differences in the two cohorts between the NFIG regions and the regions being tested. However, in the scenario where purifying selection is weakened in one of the populations, we expect to observe a bias towards smaller MAF differences in this population. The significance of these shifts can then be measured by a nonparametric statistic comparing the distributions of MAF differences between cohorts.

We applied the ASB test on exonic, promoter and intronic regions (Fig 4). Our results are consistent with the LOF-based observation of weaker purifying selection in VIKING exonic regions. In addition, ASB testing shows a similarly widespread loss of constraint in VIKING promoter regions, suggesting effects on gene expression. We observe higher MAF of very rare variants at LBC intronic regions compared to VIKING, which is most likely due to the more cosmopolitan nature of the LBC cohort and weaker purifying constraint in intronic compared to exonic and promoter regions.

### Functional impacts of rare and ultra-rare VIKING variants

Our analysis of the WGS data of the 269 Shetland individuals revealed 79 exonic variants predicted to impact gene function as significantly enriched (Fisher’s exact test) in VIKING compared to gnomAD, and occurring in 74 unique genes predicted to be largely intolerant to variation (Materials and Methods); 54 of these variants (68%) are ultra-rare (i.e. not found in gnomAD genomes dataset). A lookup for these 54 exonic variants in the order of magnitude larger gnomAD exomes dataset (v2.1.1, n = 125,748) [51] confirms their rarity in general populations: 19 variants (35%) were not found in the gnomAD exomes dataset; 16 variants (30%) were found with overall MAF ≤ 1×10^-5^ (i.e. less than 1 in 100,000), 17 variants (31%) with MAF ≤ 5×10^-5^ (i.e. less than 1 in 20,000) and the remaining 2 variants with MAF ≤ 1×10^-4^ (i.e. less than 1 in 10,000). As of Aug 27, 2019 only one of these 54 variants - rs779590262, a missense variant of uncertain significance (Variation ID 423006) – was present in ClinVar [52], a database aggregating information about genomic variation and its relationship to human health.

Given our small sample size, in order to reduce the search space and the multiple testing correction burden, from the 79 enriched exonic variants predicted to be functional we selected the 40 variants (26 of which ultra-rare) within 38 genes for which a strong evidence of gene-trait association (*p* ≤ 5×10^-8^) is reported in the GWAS Catalog (v1.0.1) [53]; among them are 13 variants (5 of which ultra-rare) in 11 distinct genes that are carried by at least 10 out of the 500 genome-sequenced Shetland individuals (Table 9 in S2 File). We performed genotype-to-phenotype analysis in the 500 VIKING individuals for those 13 variants and the 26 related quantitative traits for which data is available, but found no significant associations (nominal p < 0.0019, Bonferroni corrected for the number of traits). This was not surprising, given that we have 80% power with n = 500 and MAF ≈ 0.01 to detect a variant explaining 3% (or more) of the trait variance at that significance level. Variants with such effects sizes are relatively rare in generally healthy cohorts, highlighting the importance of sample size. We plan to investigate the identified variants and their potential phenotype correlations in ∼1600 additional VIKING samples whose WES is currently underway.

VIKING variants in promoter regions show higher levels of enrichment for ultra-rare variants than other regulatory regions (Fig 2), and analysis of the WGS data of the 269 unrelated VIKING individuals revealed 2,782 (52% ultra-rare) promoter variants significantly enriched compared to gnomAD (Materials and Methods). Since variation in promoter regions is often associated with variation in gene expression, we screened the enriched variants against the list of known eQTLs (with qval ≤ 0.05) in the GTEx dataset (v7) [54] using the data obtained from the GTEx portal [55] and found 6 rare variants (gnomAD MAF<0.05, Shetland MAF≤0.1) predicted to affect the expression of 6 distinct genes (4 of them with strong GWAS Catalog gene-trait correlation, Table 10 in S2 File), as well as 6 very common variants (gnomAD MAF > 0.4) correlated with the expression of 5 distinct genes.

## Discussion

Comparison of high-coverage WGS data for 269 unrelated individuals in the VIKING cohort from the Shetland Islands to similar data from LBC – a more cosmopolitan Scottish sample from the city of Edinburgh and around – reveals evidence of founder effects, genetic drift, and relaxation of purifying selection in Shetland. VIKING individuals exhibit genome-wide enrichment of ultra-rare variants (Table 1). On average 0.15% of all variants found in a VIKING individual have not been previously reported in the gnomAD database of WGS variants discovered in 15,496 individuals from varying ethnic origins. After careful filtering of these ultra-rare variants, we found genome-wide enrichment for ultra-rare SNPs in VIKING compared to LBC of 1.16-fold and for ultra-rare INDELs of 1.22-fold. Importantly, this enrichment is not due to an elevated rate of singleton variants in VIKING individuals, but is a result of higher rates of sharing of ultra-rare variants among Shetlanders.

The existing literature reports similar proportions of ultra-rare variants detected in isolated populations as a fraction of all variants in the population [15,19,20], although a direct comparison is difficult due to different sample sizes, sequencing approaches, genealogical criteria for participant inclusion and reference datasets. Fluctuations in the frequencies of rare variants, usually defined as variants with MAF ≤ ∼1%, have also been observed in isolate cohorts. In some cases, studies found an excess of such variants in isolated populations compared to controls [17,19,20,22], whereas in others, the isolated populations are depleted for such variants [15,21,56]. Although there is an inverse correlation between the observed frequency of a variant and the probability of it being ultra-rare [15,19,20,23], we are aware of no study to date that has explicitly investigated ultra-rare variant loads in isolates. By using the gnomAD genomes database as a reference dataset to separate the variants into ultra-rare and very rare but known (i.e. seen in gnomAD and with MAF in Non-Finnish Europeans ≤ 1%), we were able to show that while the VIKING cohort is depleted for very rare known variants, it is enriched for ultra-rare variants compared to a control cosmopolitan population, in particular for those shared by more than one unrelated individual in the isolated population. The discovered ultra-rare and rare VIKING variants which are predicted to be functional and are significantly enriched in the Shetland isolate compared to gnomAD add to the emerging catalogue of ultra-rare variants from isolated cohorts correlated with various traits of medical importance [20, 23]. Such variants are illustrative of the potential for the so called “jackpot effect” [25].

The VIKING individuals in this study were recruited as phenotypically ‘normal’ healthy individuals and represent only our first view of the Shetland isolate, with further recruitment underway. The detailed demographics and history of the Norse diaspora is still an area of active research (e.g. [57]). We look forward to deep WGS data from relevant Scandinavian populations (with compatible sequencing technologies and sample ascertainment) becoming available in the future. Such data, combined with power increasing strategies (e.g. imputation) and continual GWAS Catalog improvements, will provide much greater opportunities for discovering VIKING variants correlated with various phenotypic traits.

The availability of high-coverage WGS data allows the interrogation of both SNP and INDEL variant loads in regulatory as well as coding regions. Our results suggest that due to the reduced efficiency of purifying selection, the exonic and regulatory regions in the Shetland isolate exhibit ultra-rare SNP loads equal to the genome-wide level. We observe the same trend for higher levels of ultra-rare INDELs in many VIKING regulatory regions, particularly promoters, but VIKING exonic regions appear to be protected from short ultra-rare INDELs (of length up to 75bp), consistent with the higher expected intolerance to variation in exonic compared to regulatory regions, as well as with the previously reported finding that exonic regions are depleted of long (median size of several kbp) copy number variant deletions [58]. Excesses of functional exonic SNPs in isolated populations have been widely reported before but, to the best of our knowledge, this work is the first to provide empirical evidence that while exonic regions in an isolated population may be enriched for ultra-rare SNPs, they appear protected from short ultra-rare INDELs.

It has previously been shown that primate promoters exhibit an increased rate of evolution compared to other genomic regions [59] and this acceleration of nucleotide substitution rate is most pronounced in broadly expressed promoters [60]. It is also widely accepted that variation in regulatory regions plays an important role in complex traits, and trait-associated SNPs are known to be enriched in regulatory regions [61]. Certain recent studies [20,21,23] have suggested that isolated populations may be enriched for regulatory variation. In this work, we explicitly test this hypothesis and show that regulatory regions in the Shetland isolate generally exhibit genome-wide level of ultra-rare variant loads. This suggests that gene expression patterns may diverge relatively rapidly in isolates, producing substantial variation in gene dosage, super-imposed upon the ultra-rare variant loads in coding regions. Currently, our ability to interpret the potential effect of regulatory variants is limited to screening against eQTL databases which inevitably contain incomplete information from previous, modestly powered studies. The generation of RNA sequencing data would enable a fuller understanding of the role ultra-rare regulatory variation plays in isolated populations.

## Materials and Methods

### Participant recruitment and consent

The Viking Health Study - Shetland (VIKING) is a family-based, cross-sectional study that seeks to identify genetic factors influencing cardiovascular and other disease risk in the population isolate of the Shetland Isles in northern Scotland. The 2105 participants were recruited between 2013 and 2015, 95% of them having at least three grandparents from Shetland. Fasting blood samples were collected and many health-related phenotypes and environmental exposures were measured in each individual. All participants gave informed consent for WGS and the study was given a favourable opinion by the South East Scotland Research Ethics Committee (REC Ref 12/SS/0151).

The Lothian Birth Cohort (LBC) study sampled people living in Edinburgh and the Lothians who were recruited and tested in the Scottish Mental Surveys of 1932 and 1947 as described elsewhere [35, 36]; 1369 individuals from the LBC dataset were selected for WGS at the same facility as the VIKING samples. Ethical permissions were obtained from the Lothian Research Ethics Committee (LREC/1998/4/183; LREC/2003/2/29; 1702/98/4/183), the Multi-Centre Research Ethics Committee for Scotland (MREC/01/0/56) and the Scotland A Research Ethics Committee (07/MRE00/58). Written informed consent was obtained from all participants.

### Availability of data and materials

There is neither research ethics committee approval, nor consent from individual participants, to permit open release of the individual level research data underlying this study. The datasets generated and analysed during the current study are therefore not publicly available. Instead, the VIKING WGS data has been deposited in the EGA (accession number EGAS00001003872). VIKING DNA samples are available from Professor Jim Wilson on reasonable request, following approval by the QTL Data Access Committee and in line with the consent given by participants. LBC WGS data has been deposited in the EGA (EGAS00001003818 for the LBC1921 subset, EGAS00001003819 for the LBC1936 subset).

### Variant calling and filtering

The WGS sequencing and initial processing of the samples used in this study was performed at Edinburgh Genomics, University of Edinburgh. The starting point of our analyses were the gVCF files (GRCh38) we received for the 500 VIKING and 1369 LBC individuals, generated as follows. Demultiplexing is performed using bcl2fastq (Illumina, 2.17.1.14), allowing 1 mismatch when assigning reads to barcodes; adapters are trimmed during the demultiplexing process. BCBio-Nextgen (0.9.7) is used to perform alignment, bam file preparation and variant detection. BCBio uses bwa mem (v0.7.13 [62]) to align the raw reads to the reference genome (GRCh38; with alt, decoy and HLA sequences), then samblaster (v0.1.22 [63]) to mark the duplicated fragments, and GATK 3.4 for the indel realignment and base recalibration. The genotype likelihoods are calculated using GATK 3.4 HaplotypeCaller creating a final gVCF file.

We called the variants in each sample individually from its gVCF using GenotypeGVCFs (GATK 3.6); the identified INDELs are limited to 75bp, i.e. about half of the read length. The discovered variants for each sample were decomposed and normalized using VT (v0.5772-60f436c3 [64]). The Variants not in the 22 autosomal or the two sex chromosomes, as well as variants with AC = 0 (after decomposition) were excluded from further analyses and the filter value for all the remaining variants was reset to PASS. The variants in each individual VCF were then split to SNPs and INDELs (GATK 3.6).

An attempt to filter the variants using GATK’s VQSR approach did not produce convincing results – there was no clear separation between the filtered and retained variants in the generated plots. Instead, we adopted a hard-filtering strategy based on the variant call parameters suggested as suitable for hard-filtering by GATK [65]. The cut-off values for these parameters were determined separately for VIKING and LBC cohorts in order to account for potential batch effects since the two cohorts were sequenced at different time points and using different preparation kits – VIKING used the TruSeq PCR-Free High Throughput library, while the earlier sequenced LBC used the TruSeqNano High Throughput library. Using VariantFiltration (GATK 3.6), we marked (FILTER flag in the VCF set to FAIL) SNPs with QD < 7.4/6.9, MQ < 44.0/44.5, FS > 10.0/9.8, SOR > 2.1/2.1, MQRankSum < −2.4/-2.3 or ReadPosRankSum < −1.4/-1.4; and marked INDELs with QD < 5.3/4.9, FS > 9.1/8.8, SOR > 2.9/2.6 or ReadPosRankSum < −1.8/-1.8 in VIKING/LBC cohorts, respectively. These cut-off values were determined as the boundary to the worst-quality 5% of the variants for each of the parameters, using all variants in the SNP and INDEL VCFs for 23/62 randomly chosen VIKING/LBC samples with mean sequencing coverage >= 30x. The chosen cut-off values are more stringent than those suggested by GATK; however, one of our objectives was to minimize the number of false positive calls. In addition, we also marked as FAIL variants with DP < 10. On average, our approach lead to marking 18% and 16% of the VIKING SNPs and INDELs per sample; the corresponding values for LBC were 19% and 18%, respectively. It should be noted that in the later step of merging the variants from all samples in each cohort, we used the GATK’s KEEP_IF_ANY_UNFILTERED option. This allowed for reconsidering variants which failed to pass the hard filtering in some samples, but were called with sufficient quality in other samples to be considered trustworthy and were therefore kept for further analyses. Our analyses suggest that using this option does not introduce a bias towards rarer variants in more related populations (Fig 11 in S1 File).

The individual SNP and INDEL VCFs were lifted over to the human_g1k_v37 reference genome (using picard-2.6.0, http://broadinstitute.github.io/picard) and merged into cohort-wide SNP and INDEL VCFs (CombineVariants, GATK 3.6, using the KEEP_IF_ANY_UNFILTERED option).

Next, we selected only variants from the mappable regions of the 24 chromosomes by identifying and excluding variants from genomic regions known to produce false positive calls at a higher rate due to poor alignability (repeat rich regions and regions with low complexity) using the UCSC tracks for the CRg dataset (36mers) [66], the Duke dataset (35mers) [67] and the DAC dataset [68].

Despite the cohort-specific cut-off values used in the hard-filtering step, we further evaluated our data for the presence of potential technical artefacts due to the different kits used for sequencing of the VIKING (“PCR free”) and LBC (“PCR plus”) cohorts. We were advised (Edinburgh Genomics, personal communication, October 2018) that the use of the “PCR free” kit may result in a higher number of discovered raw INDELs genome-wide due to the elimination of the PCR amplification step in the “PCR plus” kit which may not perform optimally in regions with extreme GC content (resulting in drop of coverage in such regions for “PCR plus”). To address this, we split the mappable regions in the reference human genome to ∼ 1.75 billion consecutive blocks of length 100bp, computed the GC content for each block and assigned it to one of the 100 bins based on its GC content (one bin for each percent difference in the GC content). We then counted and compared the total number of VIKING and LBC variants for all the blocks in each of the 100 bins. As a control, we considered variants from 139 unrelated individuals from the island of Korcula, Croatia, which were sequenced with the “PCR plus” kit (same as LBC), by the same sequencing centre (Edinburgh Genomics) at a time point between the LBC and VIKING cohorts and processed by us in the same manner as for the other two cohorts. The results (Fig 12 in S1 File, Fig 13 in S1 File) suggest that indeed there is enrichment for the “PCR free” kit in regions with extreme GC content, for both SNPs and INDELs. Therefore, we identified and excluded all Shetland and LBC variants which are centred in a 100bp window with GC content less than 15% or greater than 75%. This resulted in excluding 0.35% and 0.93% of the VIKING SNP and INDEL sites, respectively; the corresponding values for the LBC cohort were 0.34% (SNPs) and 0.86% (INDELs).

### Sample selection

In order to avoid bias in the variant load analyses, we first excluded 165 samples from the LBC cohort with mean sequencing coverage < 30x, given that all but two of the 500 Shetland samples have mean coverage >= 30x. Next, we identified and excluded related samples in each cohort. We based this analysis on the discovered biallelic SNPs from the mappable regions in the 22 autosomal chromosomes with MAF >= 2% in the VIKING and LBC cohorts: 5,732,180 and 5,711,775 such markers, respectively. As a relatedness metric, we used PLINK’s [69] pi_hat statistic representing the proportional identity by descent (IBD) between two individuals and computed as pi_hat = P(IBD=2) + 0.5*P(IBD=1). We used PLINK (v1.90b4 [69]) to compute the pi_hat statistic at the markers described above for each pair of samples in each cohort and marked as related any pair of samples with pi_hat >= 0.0625, corresponding to first cousins once removed and closer, and equivalents. From these data, we identified the maximum unrelated sets of samples for each cohort (269 for VIKING and 1160 for LBC) using PRIMUS (v1.9.0 [70]). Our analysis showed that there is no significant bias towards individuals with recent immigration history (i.e., with less than four grandparents from the Shetland Isles) in the unrelated VIKING set (n = 269).

Another potential source of bias could be the presence of individuals with non-European genomic heritage. The VIKING cohort samples were investigated using the genotype array data and only those with no evidence of non-European heritage were submitted for WGS. For the LBC cohort, using data available from the 1000G Project (Phase 3) [71] as controls, we performed MDS analysis (PLINK) and identified and excluded from further analyses four samples with evidence of some African or Asian heritage.

### Variant annotation and ultra-rare variants

The variants were annotated with their predicted functional effect using VEP (v90 [72]) and with their gnomAD filter status and prevalence in all populations available in gnomAD genome dataset (gnomAD, r2.0.1 release, data from 15,496 WGS, downloaded May 26, 2017). All variants in VIKING and LBC datasets passing the hard-filtering described above, but failing the quality filters in gnomAD, were excluded from further analyses. We refer to the variants which passed both our and gnomAD filtering as “known” variants. Furthermore, from variants found in our datasets, but not found in gnomAD (i.e. ultra-rare variants), we kept for further analysis only biallelic SNPs with allele frequency (AF) in the corresponding dataset ≤ 0.1, with depth of coverage (DP) at least 8 and no more than 60 reads and genotype quality (GQ) ≥ 30; and only biallelic INDELs with AF ≤ 0.1, DP ≥ 12 and ≤ 60 and GQ ≥ 40. We refer to those variants as “ultra-rare” (Table 1), noting that some are shared between the VIKING and LBC cohorts. Our tests showed that these ultra-rare variants are generally randomly distributed genome-wide.

### ADMIXTURE analysis

Admixture analysis of the 269 VIKING and 1156 LBC unrelated individuals was performed using the ADMIXTURE tool [73, 74]. The analysis was based on 4,320,501 SNPs (not LD pruned) found in the callable regions in the 22 autosomal chromosomes with combined MAF ≥ 5% in the two cohorts and also present in gnomAD genomes dataset. The admixture_linux-1.3.0 was run with default parameters with 4 threads in unsupervised mode with K= 1, 2 and 3. The cross-validation error for each K computed using the --cv option (5 folds) identified K = 2 as the most suitable modelling choice.

### Site Frequency Spectrum (SFS) analysis

SFS analysis of the 269 VIKING and 1156 LBC unrelated individuals was performed using VCFtools (v0.1.13) [75] using the --freq2 option. Our analysis uses the high-quality variants discovered in the callable regions of the 22 autosomal chromosomes in the two cohorts of unrelated individuals, split to known variants (present in gnomAD at any frequency) and ultra-rare variants (not found in any gnomAD population). All sites with missing genotype(s) were excluded. The means and standard deviations for each frequency (Table 11 in S2 File and Fig 6 in S1 File) were computed based on subsampling the two cohorts to 50 distinct individuals each repeated 100 times (w/o replacement within subsamples, with replacement across subsamples).

### Tajima’s D analysis

Tajima’s D analysis of the 269 VIKING and 1156 LBC unrelated individuals was performed using VCFtools (v0.1.13) using the --TajimaD option and sliding windows of size 1Mb. The analysis was based on the cohorts’ known SNPs (i.e., found with passing quality in the gnomAD dataset) identified in the callable regions of the 22 autosomal chromosomes. The variants were then split into six subsets based on the functional region they reside in: 5’UTR, exon, intron, 3’UTR, ncRNA and non-coding regions. For the VIKING cohort, we computed the median Tajima’s D value and the 95% CI for each region aggregating the results observed for the 269 individuals in the ∼3000 genomic windows of size 1Mb, excluding any window with no SNPs present. For the LBC cohort, we generated 100 random subsets of size 269 unrelated individuals to match the VIKING size (without replacement within subsamples, with replacement across subsamples) and computed the cohort’s median and 95% CI aggregating the 1Mb window medians observed for each of these 100 subsets.

### ROH analysis

The runs of homozygosity (ROH) tracts were called using the `roh` function in bcftools (v1.6) [76] interrogating the high-quality SNPs discovered in the mappable regions of the 22 autosomal chromosomes of the unrelated VIKING and LBC individuals and also present in gnomAD. The `roh` command was invoked with instructions to read the alternate allele frequencies from the VCF file (-- AF-tag AF) and to ignore all variant calls with genotype quality < 30 (-G30).

To establish suitable cut-offs for partitioning the discovered ROH into to intermediate and long based on their length, we used the available data for 10 populations of European ancestry, reported in [46]. Based on these, we computed the mean (511,734bp) and the standard deviation (23,307bp) of the boundary for separating short and intermediate ROHs; the intermediate/long boundary has a mean of 1,567,737bp (s.d. 98,252bp). Conservatively, we picked 0.5Mb as intermediate ROH cut-off and 2Mb as long ROH cut-off, which is in agreement with the long ROH cut-off used in [24].

Next, we examined the density of SNP markers in the detected long and intermediate ROHs (Fig 8 in S1 File). For long ROHs, we observed a bi-modal distribution for the number of SNP markers discovered per 1Kb ROH length indicating potentially poor coverage/reliability for some ROHs, consistent with the findings in [24]. To address this issue, we excluded from further analysis all long ROHs with less than 2 or 3.5 markers per 1Kb ROH length in the VIKING and LBC cohorts, respectively. The difference between the LBC and VIKING cut-off values (ratio = 1.75) correlates well with the ratio of the total number of SNP markers given as input to bcftools for ROH calling (ratio = 1.68, LBC = 16,623,172 SNPs, VIKING = 9,890,893 SNPs). These density cut-offs also appear suitable for intermediate ROHs (Fig 8 in S1 File).

### Annotation of coding regions

Using the Ensembl (Genes 92, GRCh37.p13) data, we split the mappable regions in the reference human genome into six categories – 5’UTR (a total length of 9.3M bases), exon (30Mb), intron (906Mb), 3’UTR (27.6Mb), ncRNA (7.3Mb) and non-coding (1.1Gb) regions. Note that some regions may be overlapping, e.g. a 3’UTR region of one gene might be 5’UTR region for another, etc. The non-coding regions are defined as genome regions which do not fall in any of the above five categories.

### Annotation of regulatory regions

For the regulatory regions we used the chromatin states data generated for nine cell types by Ernst and colleagues [41], downloaded from UCSC Genome browser [77]. For each cell type we extracted the coordinates of the regions assigned to each of the 15 chromatin states (Fig 1 in [41]), followed by union of the regions in states 1, 2 and 3 to obtain a combined Promoter region (average total length of 39.2Mb, s.d. = 7.5Mb over the 9 cell types), Enhancer (130.5Mb, 16.9Mb; states 4, 5, 6 and 7), Insulator (17.4Mb, 4.7Mb; state 8), Transcription (530.3Mb, 58.8Mb; states 9, 10 and 11), Repressed (130.5Mb, 62.3Mb; state 12) and Heterochromatin (1.8Gb, 63.4Mb; state 13); we excluded from consideration states 14 and 15 (“Repetitive/CNV”).

### Significantly enriched and potentially functional exonic variants

First, we selected exonic variants which are more frequent in VIKING compared to LBC and any gnomAD population and are predicted (VEP 90) to have one of the following effects on the gene’s canonical transcript(s): stop gained, splice acceptor/donor variant, start/stop lost, missense, frameshift or inframe insertion/deletion. Next, we annotated these variants with their CADD score (CADD v1.3) and with the pLI and missense z-score values for the harbouring gene [78]. The latter two statistics are provided by the ExAC consortium and are computed based on the deviation between the observed versus expected counts of variants in each gene [39]. The pLI statistic is applicable to nonsense variants - the closer pLI is to 1, the more haploinsufficient the gene appears to be – genes with pLI ≥ 0.9 are considered extremely haploinsufficient. The z-score statistic is related to missense variants, where positive z-scores indicate increased constraint (intolerance to variation). We used the CADD, pLI and z-score information to filter the set of enriched variants (Table 12 in S2 File), which resulted in 1257 potentially functional (CADD ≥ 20 for missense and inframe variants) exonic variants in genes largely intolerant to variation.

From the set of 1257 potentially functional variants which were more frequent in VIKING compared to LBC/gnomAD, we extracted the variants which were significantly enriched compared to gnomAD. For each variant, we performed Fisher’s exact test on the number of variant alleles (AC) and total alleles (AN) at a given position using a Bonferroni corrected *p* = 0.05 / 1257 = 4.×10^-5^. For variants found in gnomAD, we used the AC_POPMAX and AN_POPMAX (the values for the population in which the variant is most prevalent); for variants not seen in gnomAD (AC = 0) we computed the corresponding AN value based on the number of individuals with coverage at least 30x at this position. In summary, we discovered 79 significantly enriched and potentially functional exonic variants in 74 unique genes.

### Significantly enriched promoter region variants in Shetland

From the 470,180 Shetland variants in the aggregated promoter regions (computed as the union of the promoter regions identified in each of the nine cell types [41]), we identified 153,381 variants which were more frequent in VIKING compared to LBC and any gnomAD population. Using the same approach as for exonic variants, we selected only variants that are significantly enriched compared to gnomAD (a Bonferroni corrected *p* = 0.05 /153381 = 3×10^-7^), which resulted in 2782 significantly enriched promoter region variants.

## Acknowledgements

SGP Consortium is funded by the Chief Scientist Office of the Scottish Government Health Directorates [SGP/1] and The Medical Research Council Whole Genome Sequencing for Health and Wealth Initiative. Members of the Scottish Genome Partnership (SGP) include Timothy J. Aitman, Andrew V. Biankin, Susanna L. Cooke, Wendy Inglis Humphrey, Sancha Martin, Lynne Mennie, Alison Meynert, Zosia Miedzybrodzka, Fiona Murphy, Craig Nourse, Javier Santoyo-Lopez, Colin A. Semple, and Nicola Williams. More information about SGP can be found at www.scottishgenomespartnership.org.

The Viking Health Study – Shetland (VIKING) was supported by the MRC Human Genetics Unit core programme grant “QTL in Health and Disease” MC_UU_0007/10. VIKING whole genome sequencing was funded by the Chief Scientist Office of the Scottish Government Health Directorates (grant reference SGP/1) and the Medical Research Council Whole Genome Sequencing for Health and Wealth Initiative (MC/PC/15080). CAS, AM and MST also acknowledge core funding to the MRC Human Genetics Unit. The work of LK was supported by an RCUK Innovation Fellowship from the National Productivity Investment Fund (MR/R026408/1).

The Lothian Birth Cohorts are supported by Age UK (Disconnected Mind grant). The whole genome sequencing of LBC1921 and LBC1936 was funded through an institutional award to the Roslin Institute from the Biotechnology and Biological Sciences Research Council (BBSRC). The LBC and SEH, DCL, and IJD were supported by the Centre for Cognitive Ageing and Cognitive Epidemiology which is funded by MRC and BBSRC (MR/K26992/1).

The CROATIA-Korcula WGS were funded by the MRC through the “QTL in Health and Disease” core programme grant.

The funders had no role in study design, data collection and analysis, decision to publish, or preparation of the manuscript.

DNA extractions and QC were performed at the Edinburgh Clinical Research Facility, University of Edinburgh; whole genome sequencing (WGS) was carried out in the Edinburgh Genomics facility, University of Edinburgh.

Nicola Pirastu selected the most appropriate participants for WGS in VIKING using the ANCHAP software. Thibaud Boutin helped with EGA submission of the VIKING dataset.

We would like to acknowledge the invaluable contributions of the research nurses in Shetland, the administrative team in Edinburgh and the people of Shetland. We also thank the Lothian Birth Cohorts’ participants and research team for their help.

## Supporting information

**S1 File. Supplementary Figures.**

**S2 File. Supplementary Tables.**

## References

1. Wright AF, Carothers AD, Pirastu M. Population choice in mapping genes for complex diseases. Nat Genet. 1999;23(4):397–404.

2. Kristiansson K, Naukkarinen J, Peltonen L. Isolated populations and complex disease gene identification. Genome Biol. 2008;9(8):109.

3. Kirin M, McQuillan R, Franklin CS, Campbell H, McKeigue PM, Wilson JF. Genomic runs of homozygosity record population history and consanguinity. PLoS One. 2010;5(11):e13996.

4. Hatzikotoulas K, Gilly A, Zeggini E. Using population isolates in genetic association studies. Brief Funct Genomics. 2014;13(5):371–7.

5. Zeggini E. Using genetically isolated populations to understand the genomic basis of disease. Genome Med. 2014;6(10):83.

6. Ober C, Tan Z, Sun Y, Possick JD, Pan L, Nicolae R, et al. Effect of Variation in CHI3L1 on Serum YKL-40 Level, Risk of Asthma, and Lung Function. N Engl J Med. 2008;358(16):1682–91.

7. Steinthorsdottir V, Thorleifsson G, Reynisdottir I, Benediktsson R, Jonsdottir T, Walters GB, et al. A variant in CDKAL1 influences insulin response and risk of type 2 diabetes. Nat Genet. 2007;39(6):770–5.

8. Scuteri A, Sanna S, Chen WM, Uda M, Albai G, Strait J, et al. Genome-wide association scan shows genetic variants in the FTO gene are associated with obesity-related traits. PLoS Genet. 2007;3(7):1200–10.

9. Thorleifsson G, Magnusson KP, Sulem P, Walters GB, Gudbjartsson DF, Stefansson H, et al. Common sequence variants in the LOXL1 gene confer susceptibility to exfoliation glaucoma. Science (80-). 2007;317(5843):1397–400.

10. Raelson J V., Little RD, Ruether A, Fournier H, Paquin B, Van Eerdewegh P, et al. Genome-wide association study for Crohn’s disease in the Quebec Founder Population identifies multiple validated disease loci. Proc Natl Acad Sci. 2007;104(37):14747–52.

11. Chen W-M, Erdos MR, Jackson AU, Saxena R, Sanna S, Silver KD, et al. Variations in the G6PC2/ABCB11 genomic region are associated with fasting glucose levels. J Clin Invest. 2008;118(7):2620–8.

12. Styrkarsdottir U, Halldorsson B V, Gretarsdottir S, Gudbjartsson DF, Walters GB, Ingvarsson T, et al. Multiple genetic loci for bone mineral density and fractures. N Engl J Med. 2008;358(22):2355–65.

13. Nakatsuka N, Moorjani P, Rai N, Sarkar B, Tandon A, Patterson N, et al. The promise of discovering population-specific disease-associated genes in South Asia. Nat Genet. 2017;49(9):1403–7.

14. Kaiser VB, Svinti V, Prendergast JG, Chau Y-Y, Campbell A, Patarcic I, et al. Homozygous loss-of-function variants in European cosmopolitan and isolate populations. Hum Mol Genet. 2015 Oct 1;24(19):5464–74.

15. Jeroncic A, Memari Y, Ritchie GR, Hendricks AE, Kolb-Kokocinski A, Matchan A, et al. Whole-exome sequencing in an isolated population from the Dalmatian island of Vis. Eur J Hum Genet. 2016;24(10):1479–87.

16. Leblond CS, Cliquet F, Carton C, Huguet G, Mathieu A, Kergrohen T, et al. Both rare and common genetic variants contribute to autism in the Faroe Islands. npj Genomic Med. 2019;4(1).

17. Gusev A, Shah MJ, Kenny EE, Ramachandran A, Lowe JK, Salit J, et al. Low-pass genome-wide sequencing and variant inference using identity-by-descent in an isolated human population. Genetics. 2012;190(2):679–89.

18. Walter K, Min JL, Huang J, Crooks L, Memari Y, McCarthy S, et al. The UK10K project identifies rare variants in health and disease. Nature. 2015;526(7571):82–9.

19. Xue Y, Mezzavilla M, Haber M, McCarthy S, Chen Y, Narasimhan V, et al. Enrichment of low-frequency functional variants revealed by whole-genome sequencing of multiple isolated European populations. Nat Commun. 2017;8.

20. Southam L, Gilly A, Süveges D, Farmaki AE, Schwartzentruber J, Tachmazidou I, et al. Whole genome sequencing and imputation in isolated populations identify genetic associations with medically-relevant complex traits. Nat Commun. 2017;8.

21. Chheda H, Palta P, Pirinen M, McCarthy S, Walter K, Koskinen S, et al. Whole-genome view of the consequences of a population bottleneck using 2926 genome sequences from Finland and United Kingdom. Eur J Hum Genet. 2017;25(4):477–84.

22. Gudbjartsson DF, Helgason H, Gudjonsson SA, Zink F, Oddson A, Gylfason A, et al. Large-scale whole-genome sequencing of the Icelandic population. Nat Genet. 2015;47(5):435–44.

23. Gilly A, Suveges D, Kuchenbaecker K, Pollard M, Southam L, Hatzikotoulas K, et al. Cohort-wide deep whole genome sequencing and the allelic architecture of complex traits. Nat Commun. 2018;9(1):4674.

24. Mooney JA, Huber CD, Service S, Sul JH, Marsden CD, Zhang Z, et al. Understanding the Hidden Complexity of Latin American Population Isolates. Am J Hum Genet. 2018;103(5):707– 26.

25. Zuk O, Schaffner SF, Samocha K, Do R, Hechter E, Kathiresan S, et al. Searching for missing heritability: Designing rare variant association studies. Proc Natl Acad Sci. 2014;111(4):E455–64.

26. Wainschtein P, Jain DP, Yengo L, Zheng Z, TOPMed Anthropometry Working Group, Trans-Omics for Precision Medicine Consortium, et al. Recovery of trait heritability from whole genome sequence data. bioRxiv. 2019;

27. Davies N. The isles: a history. Macmillan; 1999. 1296 p.

28. Capelli C, Redhead N, Abernethy JK, Gratrix F, Wilson JF, Moen T, et al. A Y chromosome census of the British Isles. Curr Biol. 2003;13(11):979–84.

29. Wilson JF, Weiss DA, Richards M, Thomas MG, Bradman N, Goldstein DB. Genetic evidence for different male and female roles during cultural transitions in the British Isles. Proc Natl Acad Sci. 2001;98(9):5078–83.

30. Goodacre S, Helgason A, Nicholson J, Southam L, Ferguson L, Hickey E, et al. Genetic evidence for a family-based Scandinavian settlement of Shetland and Orkney during the Viking periods. Heredity (Edinb). 2005;95(2):129–35.

31. Vitart V, Carothers AD, Hayward C, Teague P, Hastie ND, Campbell H, et al. Increased Level of Linkage Disequilibrium in Rural Compared with Urban Communities: A Factor to Consider in Association-Study Design. Am J Hum Genet. 2005;76(5):763–72.

32. Gilbert E, O’Reilly S, Merrigan M, McGettigan D, Vitart V, Joshi PK, et al. The genetic landscape of Scotland and the Isles. PNAS. 2019;116(38):19064–19070.

33. VIKING Project [Internet]. [cited 2019 Aug 1]. Available from: https://www.ed.ac.uk/viking/

34. Glodzik D, Navarro P, Vitart V, Hayward C, Mcquillan R, Wild SH, et al. Inference of identity by descent in population isolates and optimal sequencing studies. Eur J Hum Genet. 2013;21(10):1140–5.

35. Taylor AM, Pattie A, Deary IJ. Cohort Profile Update: The Lothian Birth Cohorts of 1921 and 1936. Int J Epidemiol. 2018;47(4):1042–1042r.

36. Deary IJ, Gow AJ, Pattie A, Starr JM. Cohort profile: The lothian birth cohorts of 1921 and 1936. Int J Epidemiol. 2012;41(6):1576–84.

37. LBC Project [Internet]. [cited 2019 Aug 1]. Available from: https://www.lothianbirthcohort.ed.ac.uk/

38. McKenna A, Hanna M, Banks E, Sivachenko A, Cibulskis K, Kernytsky A, et al. The Genome Analysis Toolkit: a MapReduce framework for analyzing next-generation DNA sequencing data. Genome Res. 2010;20(9):1297–303.

39. Lek M, Karczewski KJ, Minikel E V, Samocha KE, Banks E, Fennell T, et al. Analysis of protein-coding genetic variation in 60,706 humans. Nature. 2016;536(7616):285–91.

40. Zerbino DR, Achuthan P, Akanni W, Amode MR, Barrell D, Bhai J, et al. Ensembl 2018. Nucleic Acids Res. 2018;46(D1):D754–61.

41. Ernst J, Kheradpour P, Mikkelsen TS, Shoresh N, Ward LD, Epstein CB, et al. Mapping and analysis of chromatin state dynamics in nine human cell types. Nature. 2011;473(7345):43–9.

42. Mayr E. Systematics and the Origin of Species from the Viewpoint of a Zoologist. Harvard University Press; 1999. 372 p.

43. Wang SR, Agarwala V, Flannick J, Chiang CWK, Altshuler D, Hirschhorn JN. Simulation of finnish population history, guided by empirical genetic data, to assess power of rare-variant tests in Finland. Am J Hum Genet. 2014;94(5):710–20.

44. Tajima F. Statistical method for testing the neutral mutation hypothesis by DNA polymorphism. Genetics. 1989;123(3):585–95.

45. Neininger K, Marschall T, Helms V. SNP and indel frequencies at transcription start sites and at canonical and alternative translation initiation sites in the human genome. PLoS One. 2019;14(4):e0214816.

46. Pemberton TJ, Absher D, Feldman MW, Myers RM, Rosenberg NA, Li JZ. Genomic patterns of homozygosity in worldwide human populations. Am J Hum Genet. 2012;91(2):275–92.

47. Szpiech ZA, Xu J, Pemberton TJ, Peng W, Zöllner S, Rosenberg NA, et al. Long runs of homozygosity are enriched for deleterious variation. Am J Hum Genet. 2013;93(1):90–102.

48. McQuillan R, Leutenegger AL, Abdel-Rahman R, Franklin CS, Pericic M, Barac-Lauc L, et al. Runs of Homozygosity in European Populations. Am J Hum Genet. 2008;83(3):359–72.

49. Ceballos FC, Joshi PK, Clark DW, Ramsay M, Wilson JF. Runs of homozygosity: Windows into population history and trait architecture. Nat Rev Genet. 2018;19(4):220–34.

50. Rentzsch P, Witten D, Cooper GM, Shendure J, Kircher M. CADD: Predicting the deleteriousness of variants throughout the human genome. Nucleic Acids Res. 2019;47(D1):D886–94.

51. Karczewski KJ, Francioli LC, Tiao G, Cummings BB, Alföldi J, Wang Q, et al. Variation across 141,456 human exomes and genomes reveals the spectrum of loss-of-function intolerance across human protein-coding genes. bioRxiv. 2019;

52. Landrum MJ, Lee JM, Benson M, Brown GR, Chao C, Chitipiralla S, et al. ClinVar: Improving access to variant interpretations and supporting evidence. Nucleic Acids Res. 2018;46(Database issue):D1062–D1067.

53. GWAS Catalog [Internet]. [cited 2018 Dec 1]. Available from: https://www.ebi.ac.uk/gwas/

54. Carithers L, Ardlie K, Barcus M, Branton P, Britton A, Buia S, et al. A Novel Approach to High-Quality Postmortem Tissue Procurement: The GTEx Project. Biopreserv Biobank. 2015;13(5):311–9.

55. GTEx (v7) [Internet]. [cited 2018 Dec 1]. Available from: https://gtexportal.org/home/

56. Pedersen CET, Lohmueller KE, Grarup N, Bjerregaard P, Hansen T, Siegismund HR, et al. The effect of an extreme and prolonged population bottleneck on patterns of deleterious variation: Insights from the Greenlandic Inuit. Genetics. 2017;205(2):787–801.

57. Margaryan A, Lawson DJ, Sikora M, Racimo F, Rasmussen S, Moltke I, et al. Population genomics of the Viking world. bioRxiv. 2019;

58. Sudmant PH, Mallick S, Nelson BJ, Hormozdiari F, Krumm N, Huddleston J, et al. Global diversity, population stratification, and selection of human copy-number variation. Science (80-). 2015;349(6253):aab3761.

59. Taylor MS, Kai C, Kawai J, Carninci P, Hayashizaki Y, Semple CAM. Heterotachy in mammalian promoter evolution. PLoS Genet. 2006;2(4):627–39.

60. Young RS, Hayashizaki Y, Andersson R, Sandelin A, Kawaji H, Itoh M, et al. The frequent evolutionary birth and death of functional promoters in mouse and human. Genome Res. 2015;25(10):1546–57.

61. Kindt ASD, Navarro P, Semple CAM, Haley CS. The genomic signature of trait-associated variants. BMC Genomics. 2013;14:108.

62. Li H, Durbin R. Fast and accurate short read alignment with Burrows-Wheeler Transform. Bioinformatics. 2009;25(5):1754–60.

63. Faust GG, Hall IM. SAMBLASTER: Fast duplicate marking and structural variant read extraction. Bioinformatics. 2014;30(17):2503–5.

64. Tan A, Abecasis GR, Kang HM. Unified representation of genetic variants. Bioinformatics. 2015;31(13):2202–4.

65. GATK Hard Filtering [Internet]. [cited 2017 Jun 1]. Available from: https://software.broadinstitute.org/gatk/documentation/article.php?id=3225

66. CRg dataset (36mers) [Internet]. [cited 2017 Jul 1]. Available from: http://hgdownload.cse.ucsc.edu/goldenPath/hg19/encodeDCC/wgEncodeMapability/wgEncodeCrgMapabilityAlign36mer.bigWig

67. Duke dataset (35mers) [Internet]. [cited 2017 Jul 1]. Available from: http://hgdownload.cse.ucsc.edu/goldenPath/hg19/encodeDCC/wgEncodeMapability/wgEncodeDukeMapabilityUniqueness35bp.bigWig

68. DAC dataset [Internet]. [cited 2017 Jul 1]. Available from: http://hgdownload.cse.ucsc.edu/goldenPath/hg19/encodeDCC/wgEncodeMapability/wgEncodeDacMapabilityConsensusExcludable.bed.gz

69. Purcell SM, Chang CC, Chow CC, Tellier LC, Lee JJ, Vattikuti S. Second-generation PLINK: rising to the challenge of larger and richer datasets. Gigascience. 2015;4(7).

70. Staples J, Qiao D, Cho MH, Silverman EK, Nickerson DA, Below JE. PRIMUS: Rapid reconstruction of pedigrees from genome-wide estimates of identity by descent. Am J Hum Genet. 2014;95(5):553–64.

71. Auton A, Abecasis GR, Altshuler DM, Durbin RM, Bentley DR, Chakravarti A, et al. A global reference for human genetic variation. Nature. 2015 Oct 30;526(7571):68–74.

72. McLaren W, Gil L, Hunt SE, Riat HS, Ritchie GRS, Thormann A, et al. The Ensembl Variant Effect Predictor. Genome Biol. 2016;17(1):122.

73. Alexander DH, Novembre J, Lange K. Fast model-based estimation of ancestry in unrelated individuals. Genome Res. 2009;19(9):1655–1664.

74. ADMIXTURE tool [Internet]. [cited 2019 Aug 1]. Available from: http://software.genetics.ucla.edu/admixture/index.html

75. Danecek P, Auton A, Abecasis G, Albers CA, Banks E, DePristo MA, et al. The variant call format and VCFtools. Bioinformatics. 2011;27(15):2156–2158.

76. Li H. A statistical framework for SNP calling, mutation discovery, association mapping and population genetical parameter estimation from sequencing data. Bioinformatics. 2011;27(21):2987–93.

77. 15 chromatin states data tracks [Internet]. [cited 2018 Nov 1]. Available from: http://genome.ucsc.edu/cgi-bin/hgFileUi?g=wgEncodeBroadHmm&db=hg19

78. pLI and z-score file [Internet]. [cited 2017 Oct 1]. Available from: ftp://ftp.broadinstitute.org/pub/ExAC_release/release0.3.1/functional_gene_constraint

