## Supplementary Figures (S1 File) for "Increased ultra-rare variant load in an isolated Scottish population impacts exonic and regulatory regions"

### S1 File: Supplementary Figures

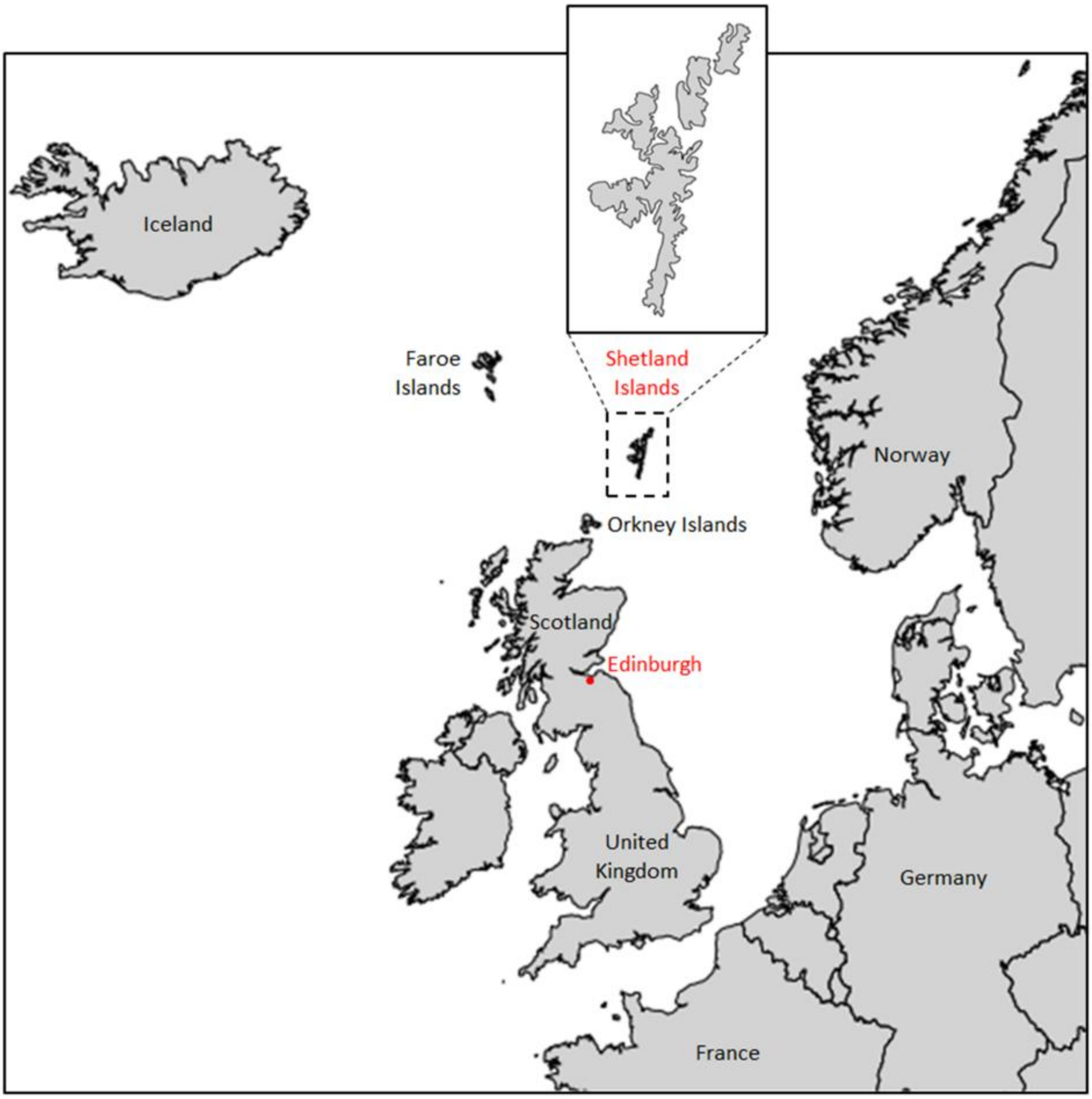


**S1 Fig. Geographic localization of the Shetland Islands.** The Shetland Islands lie scattered between ~160-290 km (~100-180 miles) north of the Scottish mainland and consist of a group of ~100 islands, of which 16 are inhabited, with a population of ~23,000.

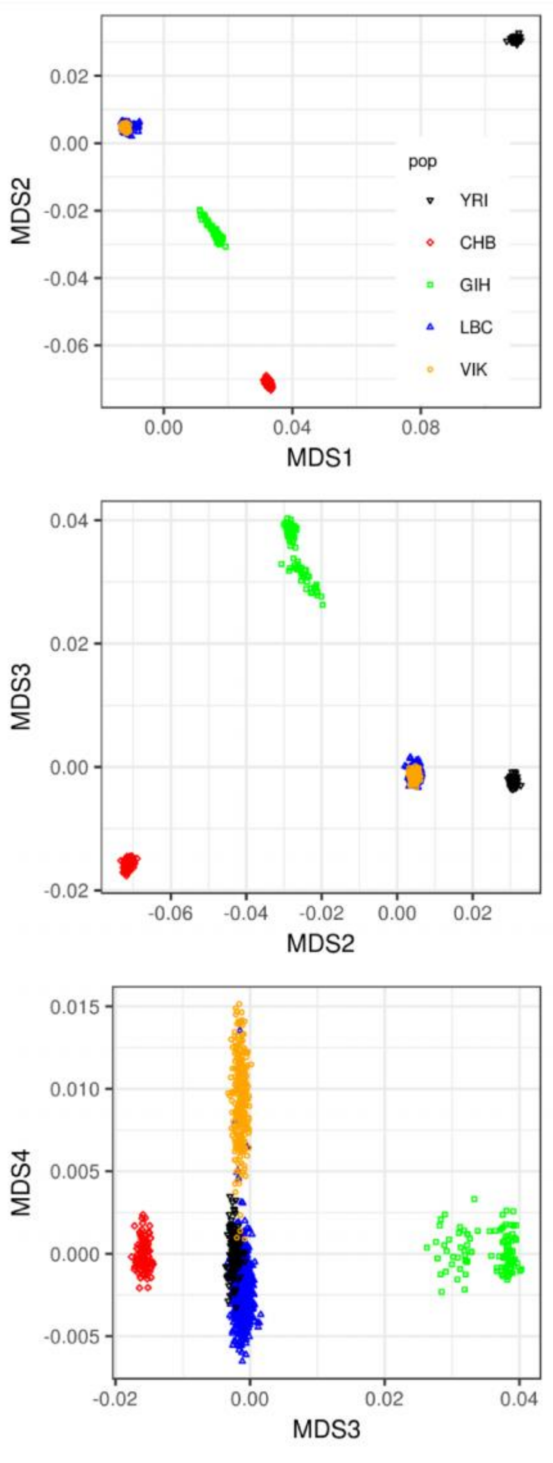


**S2 Fig. VIKING individuals are distinct from LBC controls.** MDS analysis performed with PLINK (1.90b4) of the 269 Shetland (VIK) and 1156 unrelated Lothian (LBC) individuals, using 1000G Gujarati Indians in Houston, Texas (South Asian, GIH, n = 103), Han Chinese in Beijing, China (East Asian, CHB, n = 103) and Yoruba in Ibadan, Nigeria (African, YRI, n = 108) populations as outgroups. The analysis is based on 9,070,695 marker loci from the callable regions in the 22 autosomal chromosomes for which a SNP with MAF ≥ 1% is found in the full 1000G dataset. MDS1 separates African (YRI) from European (LBC+VIK) samples, MDS2: African (YRI) vs East Asian (CHB), MDS3: East Asian (CHB) vs South Asian (GIH), and MDS4 separates VIKING from LBC/rest.

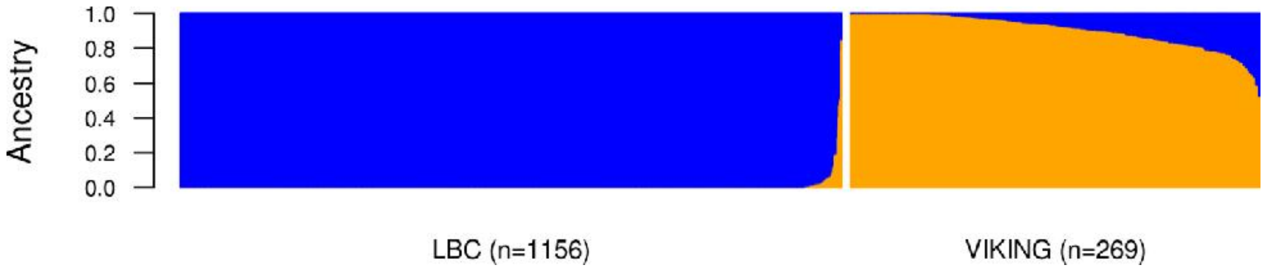


**S3 Fig. Admixture analysis of the VIKING and LBC individuals.** Admixture analysis (admixture_linux-1.3.0 with K = 2) of the 269 VIKING and 1156 LBC individuals based on 4,320,501 not LD-pruned SNPs found in the callable regions in the 22 autosomal chromosomes with combined MAF ≥ 5% in the two cohorts and also present in gnomAD genomes dataset. The tool was run with default parameters with 4 threads in unsupervised mode with K= 1, 2 and 3. The cross-validation error for each K computed using the --cv option (5 folds) identified K = 2 as the most suitable modelling choice.

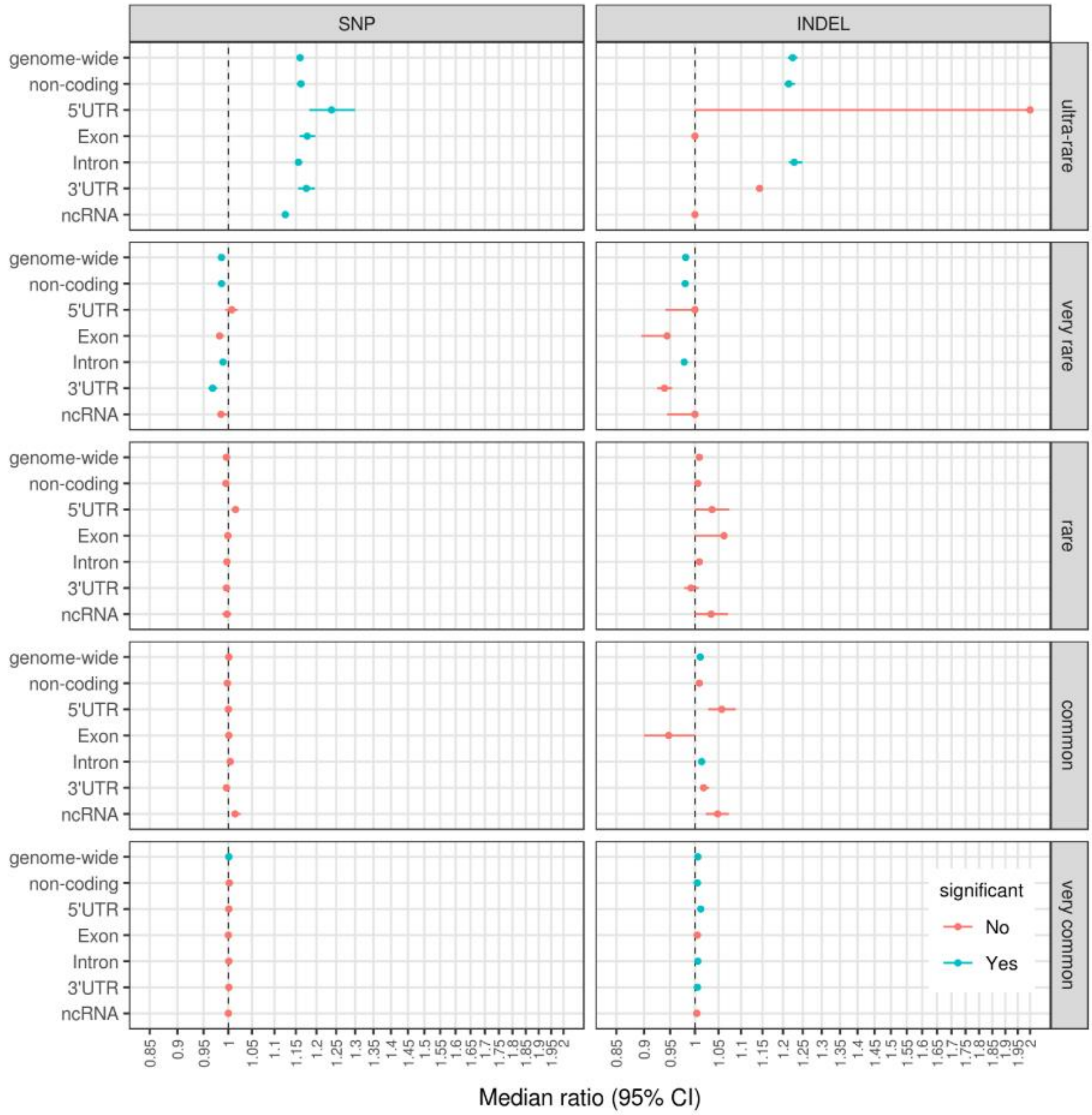


**S4 Fig. Variant load comparison in coding regions.** Circles represent the ratio of the median number of variants in a VIKING individual to the median number of variants in an LBC individual; whiskers are 95% CI based in 10k randomly selected LBC subsets (n = 269). Significance: at least 95% of the 10k subsets have p-value ≤ $8x10^{-4}$ (Bonferroni corrected) and no overlap between the 95% CI for the LBC median and the VIKING median value (see S2 Table).

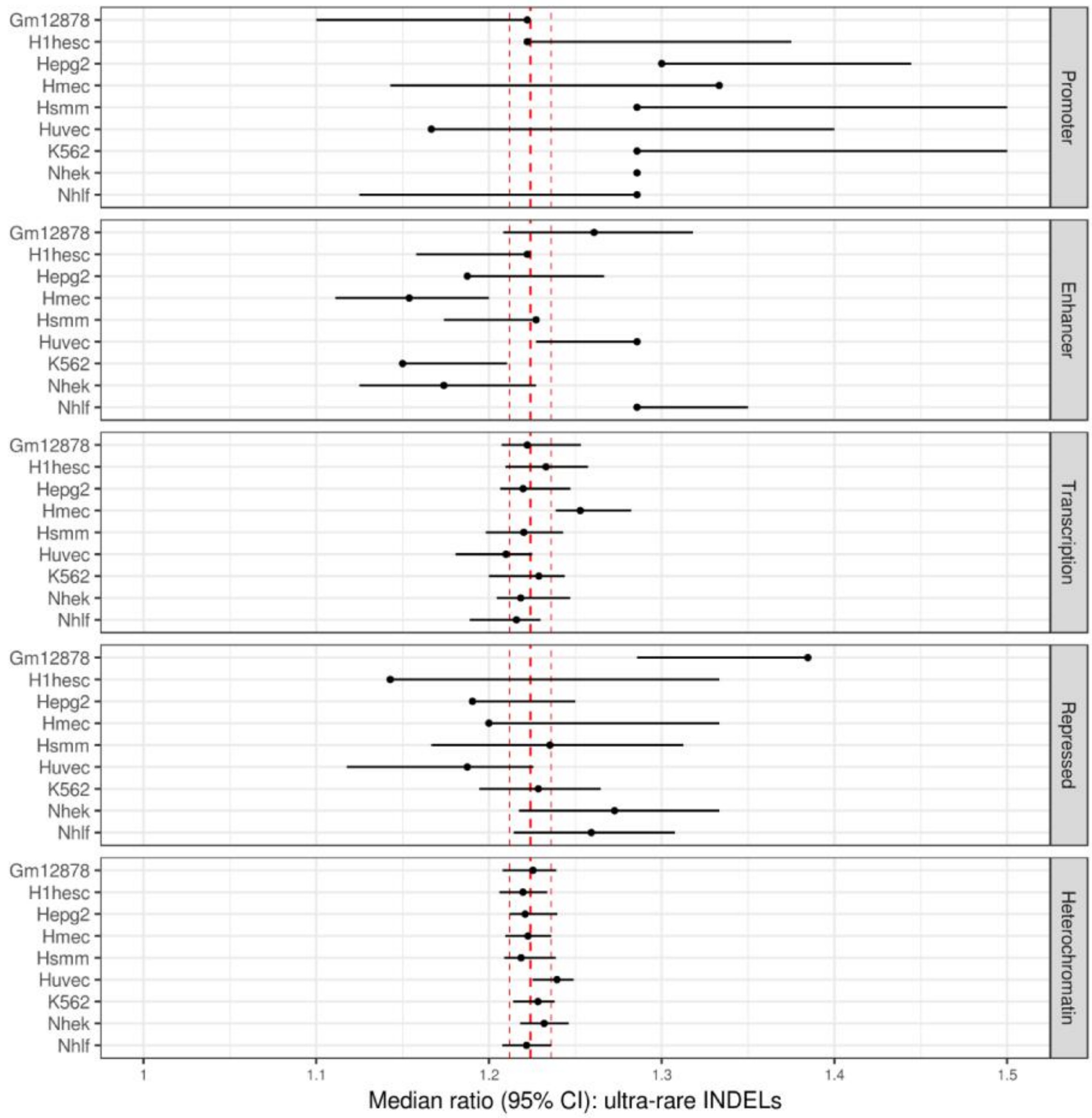


**S5 Fig. Significant differences in variant load in regulatory regions for ultra-rare INDELs in 9 cell types.** Circle dots represent the ratio of the median number of variants in a VIKING individual to the median number of variants in an LBC individual; whiskers are 95% CI based in 10k randomly selected LBC subsets (n = 269). Significance: at least 95% of the 10k subsets have p-value ≤ $2x10^{-4}$ (Bonferroni corrected) and no overlap between the 95% CI for the LBC median and the VIKING median value. Red vertical lines represent the median genome-wide enrichment for ultra-rare INDELs and its 95% CI. No significant difference was observed for any of the cell types in the insulator regions (not plotted).

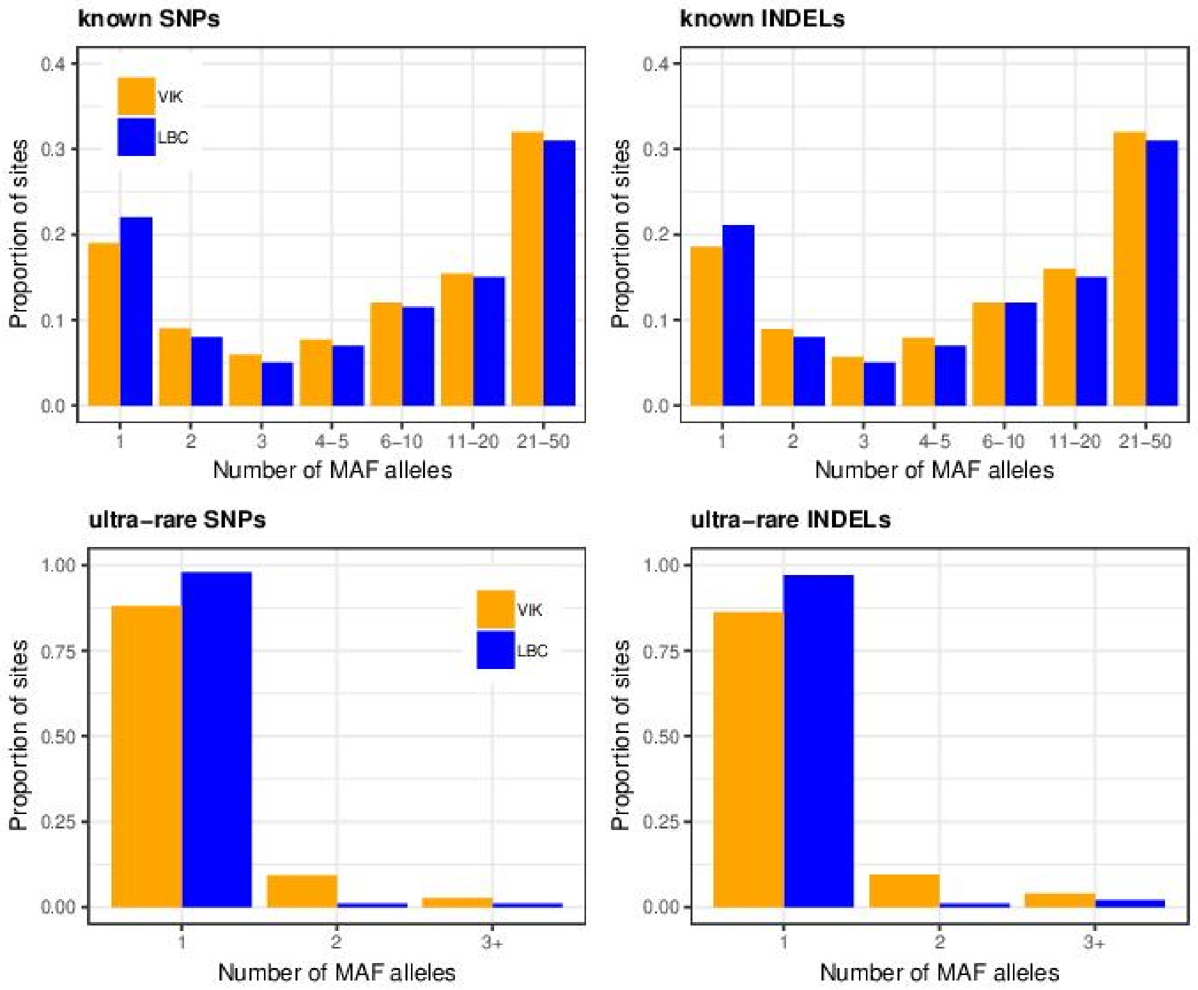


**S6 Fig. Folded SFS analysis of the VIKING and LBC cohorts.** The analysis is based on high-quality SNPs/INDELs discovered in the callable regions of the 22 autosomal chromosomes in the two cohorts of unrelated individuals, split to known variants (present in gnomAD at any frequency) and ultra-rare variants (not found in any gnomAD population). All sites with missing genotype(s) were excluded. The means and standard deviations for each frequency were computed based on subsampling the two cohorts to 50 individuals each repeated 100 times (Table 11 in S2 File).

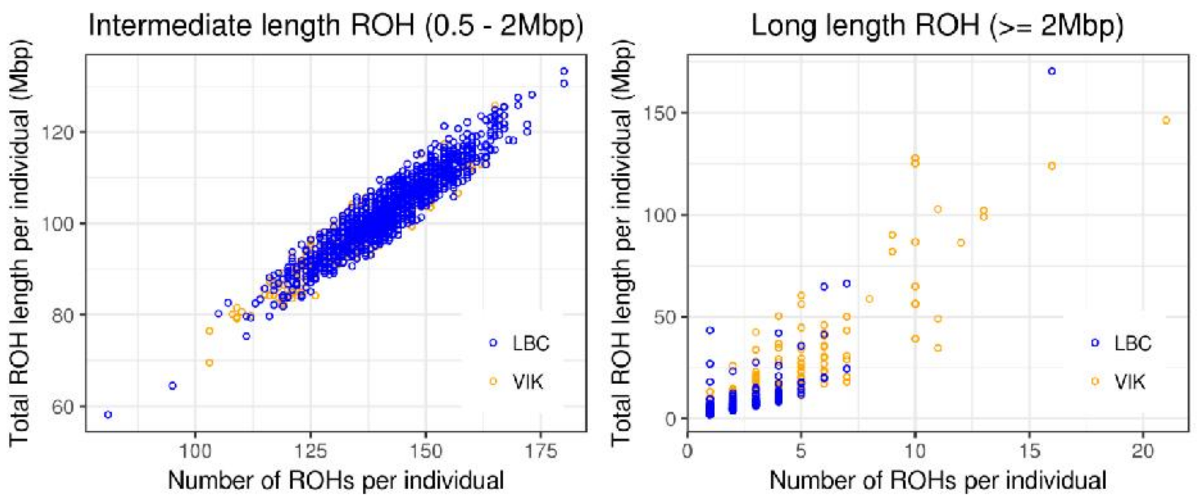


**S7 Fig. Runs of homozygosity (ROH) comparison of the VIKING and LBC cohorts.** Left panel: intermediate length ROH = 0.5 – 2Mb; Right panel: long ROH ≥ 2Mb. Each marker represents a VIKING or LBC individual. Significance: at least 95% of the 10k subsets have p-value ≤ 0.0125 (Bonferroni corrected) and no overlap between the 95% CI for the LBC median and the VIKING median value.

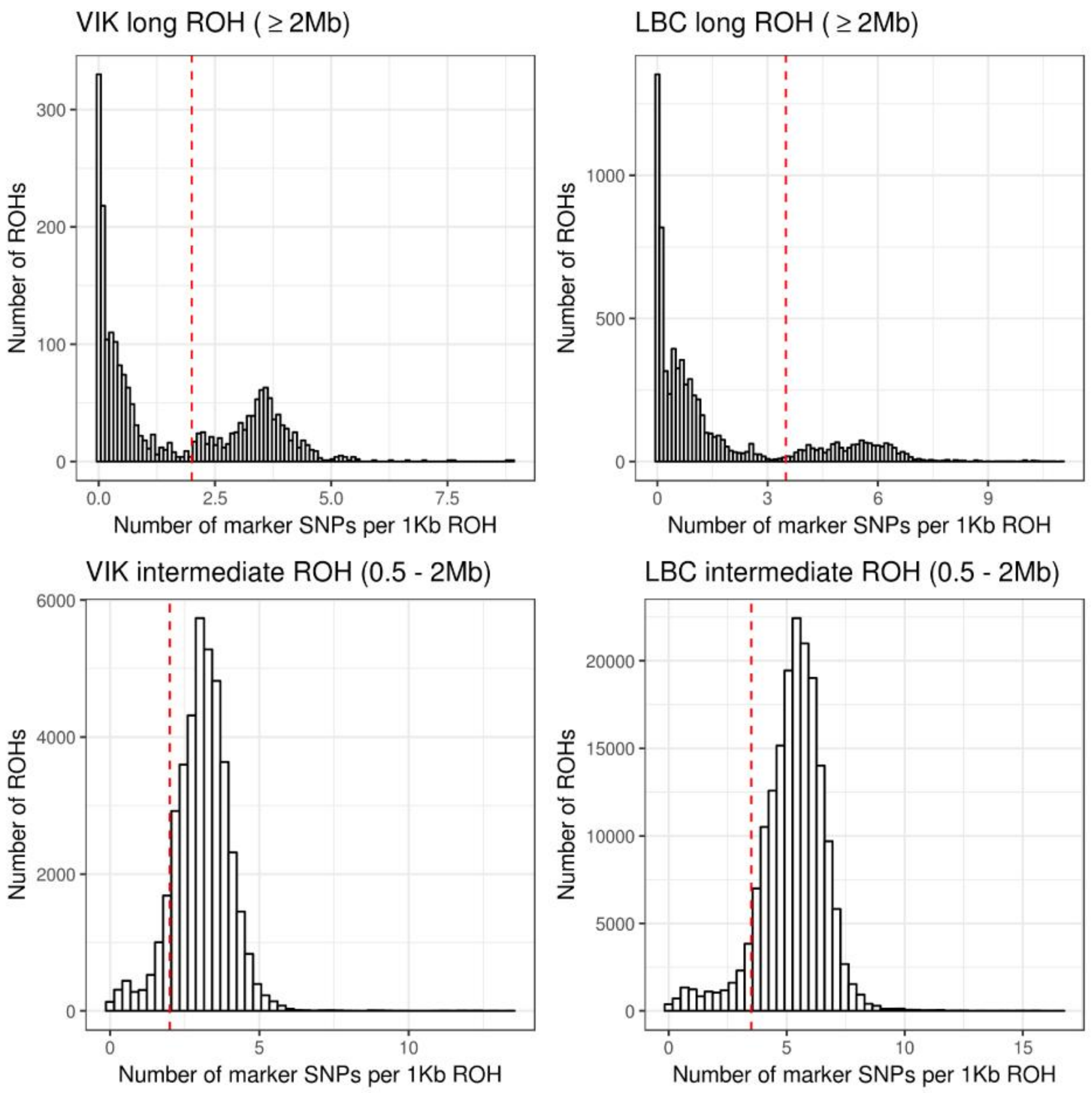


**S8 Fig**. **ROH filtering cut-offs based on SNP density.** For long ROHs (top panel), we observe a bi-modal distribution for the number of SNP markers discovered per 1Kb ROH length indicating potentially poor coverage/reliability for some ROHs. Long ROHs with less than 2 or 3.5 markers per 1Kb ROH length (vertical red lines) in the VIKING and LBC cohorts, respectively, were excluded from further analysis. The chosen density cut-offs also appear suitable for intermediate ROHs (bottom panel).

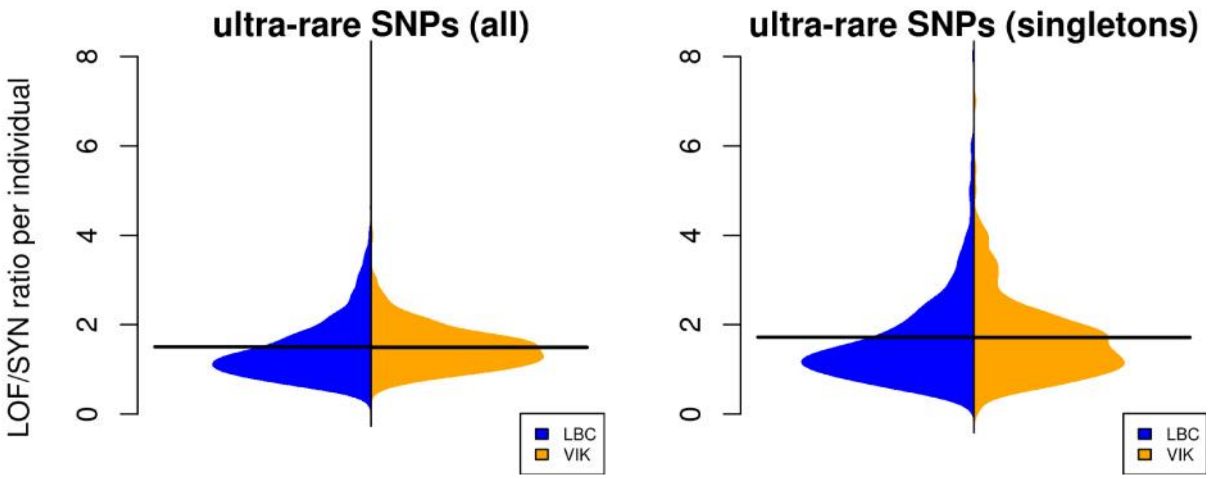


**S9 Fig. LOF/SYN ultra-rare variant ratio per individual in VIKING (n=269) and LBC (n=1156) cohorts.** LOF: loss-of-function variant (stop gain, splice donor/acceptor, missense with CADD score ≥ 20); SYN: synonymous variant. Left panel: based on singleton ultra-rare SNPs only; Right panel: based on all ultra-rare SNPs. Black horizontal lines represent cohort means: all ultra-rare LBC $_{LOF/SYN}$ ratio = 1.504 (median = 1.286), VIKING $_{LOF/SYN}$ ratio = 1.492 (median = 1.444); singleton ultra-rare LBC $_{LOF/SYN}$ ratio = 1.720 (median = 1.400), VIKING $_{LOF/SYN}$ ratio = 1.714 (median = 1.523). The LOF/SYN ratios reported here are computed at individual level and then aggregated (i.e., mean/median), while those reported in the main text (LBC $_{LOF/SYN}$ = 1.40 and VIKING $_{LOF/SYN}$ = 1.47) are computed directly at cohort level (i.e. as the ratio between the number of all LOF variants in the cohort and the number of all SYN variants in cohort).

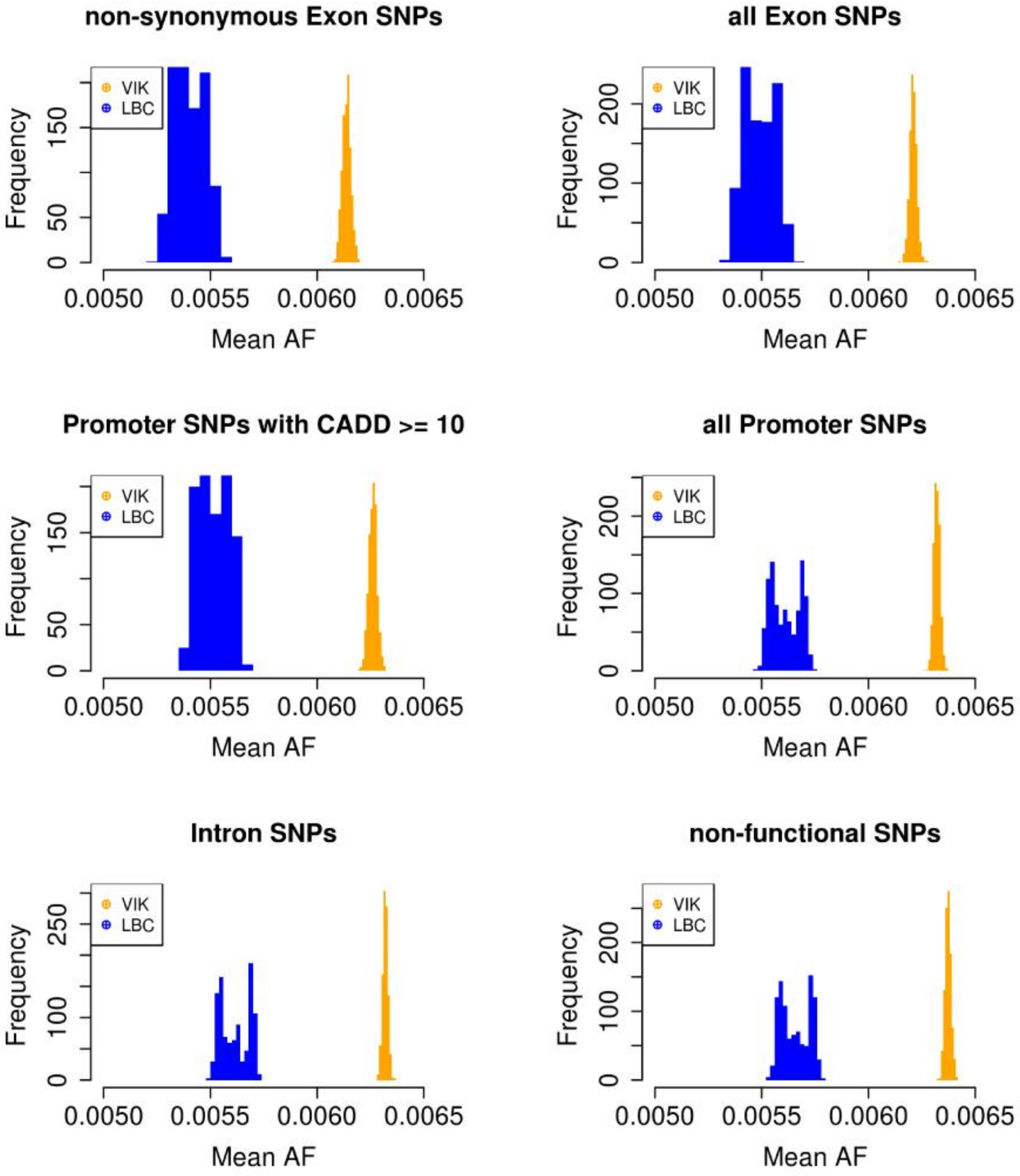


**S10 Fig. VIKING and LBC MAF for shared very rare gnomAD SNPs ($MAF_{NFE}$ ≤ 1%).** Histograms of the mean AF of very rare Non-Finnish European SNPs observed both in the 269 VIKING individuals and 1000 randomly selected LBC subsets (n = 269). Mean number of shared very rare SNPs (1000 LBC subsets): non-synonymous exonic = 13,590; all exonic = 22,802; promoter (CADD ≥ 10) = 14,533; all promoter = 78,781; intronic = 786,271 and non-functional intergenic = 483,429 variants.

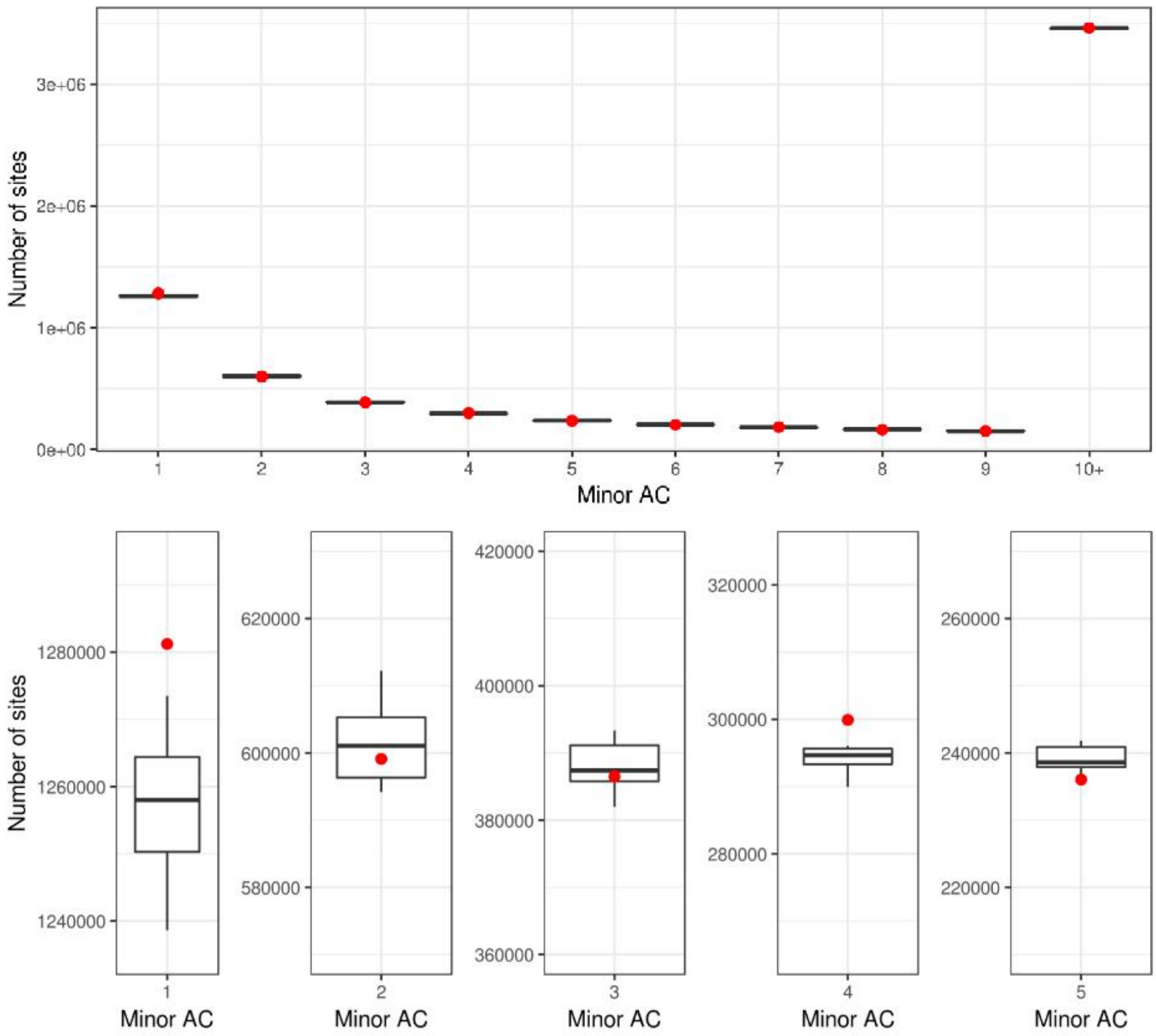


**S11 Fig. KEEP_IF_ANY_UNFILTERED does not introduce a bias towards rarer variants in more related populations.** The red points depict data from the 34 unrelated (pi_hat = 0) VIK individuals. Black boxplots represent the data from 10 control subsets of 34 VIK individuals randomly selected from the remaining 466 VIK individuals (w/o replacement within subsets, with replacement across subsets). The upper and lower "hinges" correspond to the first and third quartiles (the 25th and 75th percentiles); the upper whisker extends from the hinge to the highest value that is within 1.5 * IQR of the hinge, where IQR is the inter-quartile range, or distance between the first and third quartiles; the lower whisker extends from the hinge to the lowest value within 1.5 * IQR of the hinge. Top panel: sites split to those with minor AC from 1 to 9 and 10+, lower panel: zoom in rarer sites with minor AC = 1, 2, 3, 4 and 5.

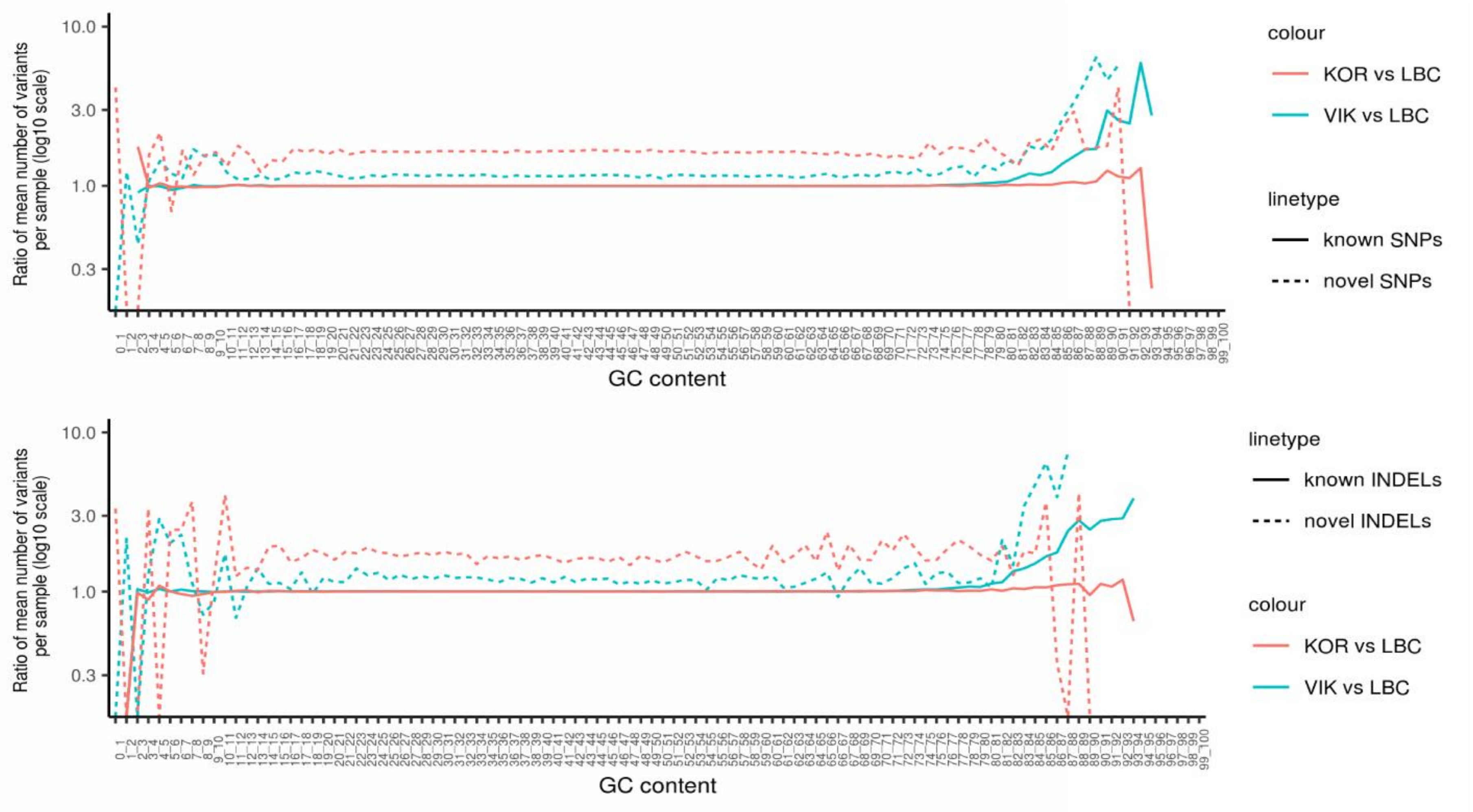


**S12 Fig. Kit effect on the number of discovered variants as a function of the GC content.** Upper panel: SNPs, lower panel: INDELs. VIKING sequenced with TruSeq PCR-Free High Throughput library kit ("PCR-free"); LBC and Korcula with TruSeqNano High Throughput library kit ("PCR plus"). The main difference between the number of variants discovered in samples processed with the "PCR-free" and "PCR plus" kits is in regions with extreme GC content (GC ≤ 15% and GC ≥ 75%), due to the different coverage efficiency by the two kits in such regions (see S13 Fig).

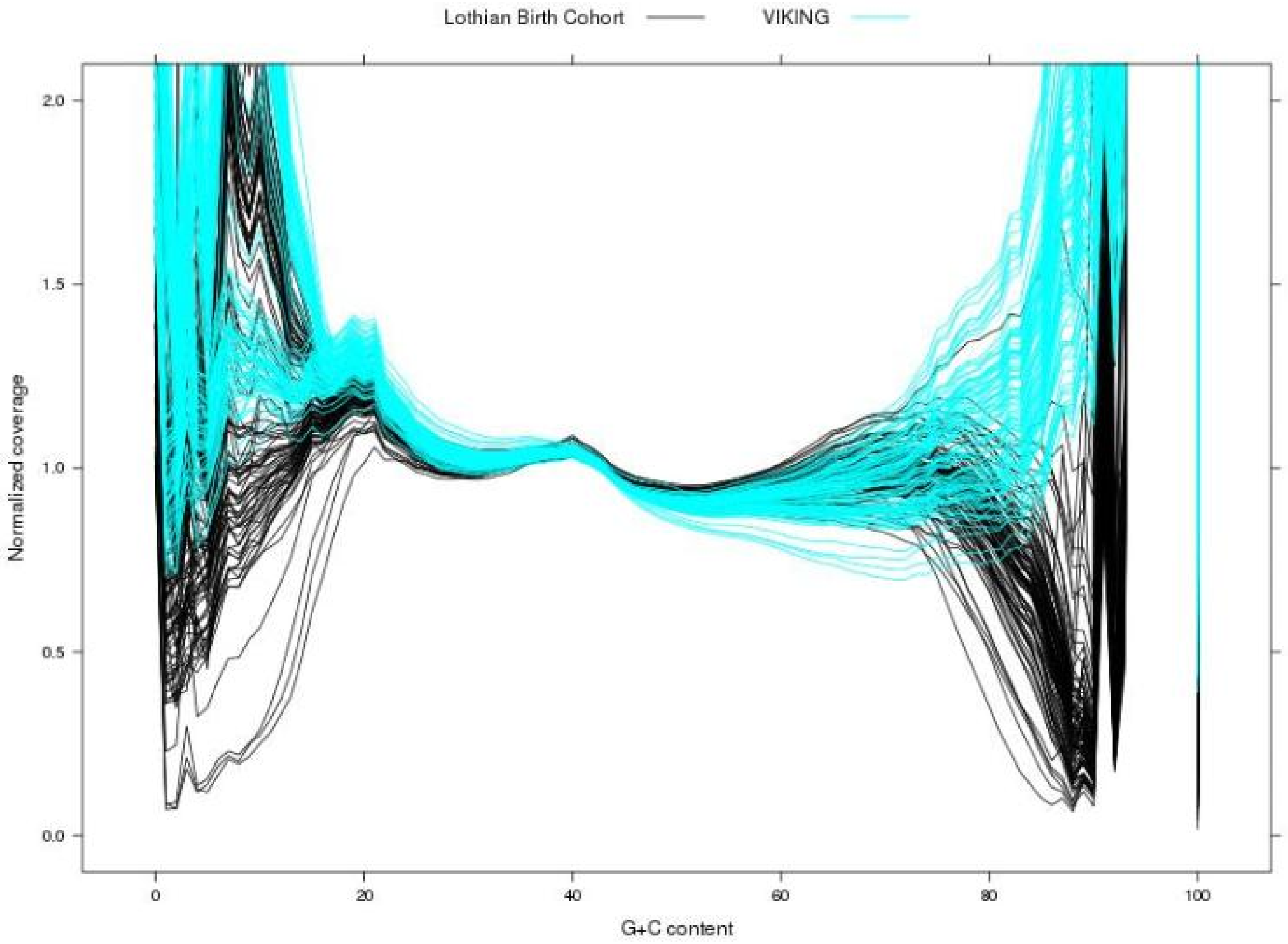


**S13 Fig. Kit effect on the coverage as a function of the GC content.** VIKING sequenced with TruSeq PCR-Free High Throughput library kit ("PCR-free"); LBC sequenced with TruSeqNano High Throughput library kit ("PCR plus"). S13 Fig is based on 100 VIK and 100 LBC samples, randomly selected.
