## Supplementary Tables (S2 File) for "Increased ultra-rare variant load in an isolated Scottish population impacts exonic and regulatory regions"

### S2 File: Supplementary Tables

**S1 Table. Average number of high-quality variant alleles found per unrelated individual in the VIKING and LBC cohorts**. The variants are stratified by their presence in the full gnomAD genomes dataset (n = 15,496) and their prevalence in gnomAD Non-Finnish Europeans (NFE) population (n=7,509).

| **gnomADg MAF** | **VIKING (n=269)** | | **LBC (n=1156)** | |
|---|---|---|---|---|
| | **SNP (s.d.)** | **INDEL (s.d.)** | **SNP (s.d.)** | **INDEL (s.d.)** |
| **Total** | 3,528,153 | 356,552 | 3,524,508 | 354,424 |
| **very common**: $MAF_{NFE} > 10\%$ | 3,287,505 (8,697) | 331,347 (1,041) | 3,283,725 (8,867) | 329,440 (1,322) |
| **common**: $5\% < MAF_{NFE} \leq 10\%$ | 115,419 (2,166) | 11,954 (249) | 115,316 (2,217) | 11,805 (264) |
| **rare**: $1\% < MAF_{NFE} \leq 5\%$ | 86,203 (1,619) | 8,730 (215) | 86,513 (1,732) | 8,662 (214) |
| **very rare**: $MAF_{NFE} \leq 1\%$ | 33,857 (792) | 4,017 (106) | 34,481 (1,873) | 4,104 (206) |
| **ultra-rare**: not in gnomADg | 5,169 (164) | 504 (28) | 4,472 (410) | 413 (40) |

**S2 Table. VIKING vs LBC: SNP load comparison in coding and coding related regions (alleles per individual per 1Mb).** Very common: variants with MAF > 10% in Non-Finnish Europeans (NFE, gnomADg, n=7,509); common: 5% < $MAF_{NFE} \leq 10\%$; rare: 1% < $MAF_{NFE} \leq 5\%$; very rare: $MAF_{NFE} \leq 1\%$; ultra-rare: not observed in any gnomADg individual (n=15,496). Median number and 95% CI of LBC alleles (forth column) for each frequency class is computed based on 10,000 random subsets (n = 269, matching VIKING size); last two columns represent the median p-value (and 95% CI) and the number of tests with p-value smaller than the Bonferroni corrected threshold. To annotate the number of variants in a frequency class as significantly different (shown in bold), we required at least 95% of the 10,000 subsets to have p-value ≤ $8x10^{-4}$ (Bonferroni corrected) and no overlap between the 95% CI for the LBC and VIKING median values. Region annotation (5'UTR, Exon, Intron, 3'UTR, ncRNA) is based on Ensembl data (GRCh37.p13, Ensembl Genes 92) for the mappable sections of the 22 autosomal chromosomes; the remaining 1.1Gb of the mappable regions in the reference human genome is annotated as "non-coding".

| Region | gnomAD Frequency Class | VIKING median | LBC 10k subsets median & 95%CI | VIKING/LBC ratio median & 95%CI | Wilcoxon rank sum test | |
|---|---|---|---|---|---|---|
| | | | | | *p* : median & 95% CI | number of tests with *p* ≤ $8x10^{-4}$ |
| **5'UTR** | very common | 1119.76 | 1119.00 [1117.60, 1120.19] | 1.001 [1.000, 1.002] | $2.3x10^{-1}$ [$1.7x10^{-2}$, $9.0x10^{-1}$] | 1 |
| | common | 43.07 | 43.07 [42.75, 43.39] | 1.000 [0.993, 1.008] | $6.6x10^{-1}$ [$1.6x10^{-1}$, $9.9x10^{-1}$] | 0 |
| | rare | 36.18 | 35.64 [35.43, 35.96] | 1.015 [1.006, 1.021] | $3.8x10^{-2}$ [$8.8x10^{-4}$, $4.0x10^{-1}$] | 224 |
| | very rare | 16.47 | 16.36 [16.15, 16.58] | 1.007 [0.994, 1.020] | $6.5x10^{-1}$ [$1.3x10^{-1}$, $9.8x10^{-1}$] | 0 |
| | **ultra-rare** | **2.80** | **2.26 [2.15, 2.37]** | **1.238 [1.182, 1.300]** | **$1.8x10^{-24}$ [$2.6x10^{-29}$, $5.8x10^{-20}$]** | **10000** |
| **Exon** | very common | 777.50 | 777.74 [777.07, 778.37] | 1.000 [0.999, 1.001] | $6.4x10^{-1}$ [$1.2x10^{-1}$, $9.8x10^{-1}$] | 0 |
| | common | 31.55 | 31.52 [31.35, 31.72] | 1.001 [0.995, 1.006] | $6.6x10^{-1}$ [$1.5x10^{-1}$, $9.8x10^{-1}$] | 0 |
| | rare | 27.95 | 27.98 [27.88, 28.15] | 0.999 [0.993, 1.002] | $6.5x10^{-1}$ [$1.4x10^{-1}$, $9.8x10^{-1}$] | 0 |
| | very rare | 14.47 | 14.74 [14.61, 14.84] | 0.982 [0.975, 0.991] | $1.4x10^{-2}$ [$2.8x10^{-4}$, $2.2x10^{-1}$] | 678 |
| | **ultra-rare** | **2.43** | **2.07 [2.03, 2.10]** | **1.177 [1.159, 1.197]** | **$2.2x10^{-29}$ [$3.4x10^{-34}$, $6.2x10^{-25}$]** | **10000** |
| **Intron** | very common | 1479.11 | 1477.75 [1476.96, 1478.33] | 1.001 [1.001, 1.001] | $2.8x10^{-3}$ [$2.6x10^{-5}$, $7.1x10^{-2}$] | 2790 |
| | common | 52.86 | 52.67 [52.53, 52.85] | 1.004 [1.000, 1.006] | $1.1x10^{-1}$ [$4.8x10^{-3}$, $7.2x10^{-1}$] | 26 |
| | rare | 40.11 | 40.23 [40.10, 40.37] | 0.997 [0.994, 1.000] | $1.1x10^{-1}$ [$5.1x10^{-3}$, $6.9x10^{-1}$] | 19 |
| | **very rare** | **16.06** | **16.24 [16.19, 16.32]** | **0.989 [0.984, 0.991]** | **$6.7x10^{-7}$ [$1.0x10^{-9}$, $1.4x10^{-4}$]** | **9961** |
| | **ultra-rare** | **2.46** | **2.13 [2.11, 2.14]** | **1.156 [1.149, 1.165]** | **$4.0x10^{-81}$ [$4.0x10^{-85}$, $1.7x10^{-76}$]** | **10000** |
| **3'UTR** | very common | 1185.76 | 1184.31 [1183.08, 1185.47] | 1.001 [1.000, 1.002] | $4.7x10^{-2}$ [$1.3x10^{-3}$, $4.6x10^{-1}$] | 156 |
| | common | 45.45 | 45.63 [45.41, 45.88] | 0.996 [0.991, 1.001] | $2.7x10^{-1}$ [$2.1x10^{-2}$, $9.2x10^{-1}$] | 0 |
| | rare | 36.52 | 36.67 [36.42, 36.89] | 0.996 [0.990, 1.003] | $2.4x10^{-1}$ [$1.8x10^{-2}$, $9.2x10^{-1}$] | 2 |
| | **very rare** | **15.60** | **16.10 [15.96, 16.25]** | **0.968 [0.960, 0.977]** | **$5.9x10^{-8}$ [$4.1x10^{-11}$, $2.7x10^{-5}$]** | **9995** |
| | **ultra-rare** | **2.43** | **2.07 [2.03, 2.10]** | **1.175 [1.155, 1.196]** | **$2.0x10^{-33}$ [$2.3x10^{-38}$, $1.4x10^{-28}$]** | **10000** |
| **ncRNA** | very common | 1681.90 | 1682.45 [1680.11, 1684.92] | 1.000 [0.998, 1.001] | $6.8x10^{-1}$ [$1.7x10^{-1}$, $9.9x10^{-1}$] | 0 |
| | common | 59.82 | 58.99 [58.31, 59.41] | 1.014 [1.007, 1.026] | $3.0x10^{-1}$ [$2.3x10^{-2}$, $9.4x10^{-1}$] | 2 |
| | rare | 45.00 | 45.14 [44.73, 45.55] | 0.997 [0.988, 1.006] | $6.7x10^{-1}$ [$1.6x10^{-1}$, $9.8x10^{-1}$] | 0 |
| | very rare | 17.70 | 17.97 [17.70, 18.11] | 0.985 [0.977, 1.000] | $4.2x10^{-1}$ [$4.6x10^{-2}$, $9.7x10^{-1}$] | 1 |
| | **ultra-rare** | **2.47** | **2.20 [2.20, 2.20]** | **1.125 [1.125, 1.125]** | **$3.3x10^{-8}$ [$2.7x10^{-11}$, $1.1x10^{-5}$]** | **9999** |
| **non-coding** | very common | 1723.29 | 1720.59 [1719.61, 1721.27] | 1.002 [1.001, 1.002] | $2.2x10^{-5}$ [$6.8x10^{-8}$, $1.9x10^{-3}$] | 9421 |
| | common | 59.35 | 59.47 [59.26, 59.62] | 0.998 [0.995, 1.002] | $6.0x10^{-1}$ [$1.0x10^{-1}$, $9.8x10^{-1}$] | 0 |
| | rare | 43.68 | 43.89 [43.76, 44.04] | 0.995 [0.992, 0.998] | $7.0x10^{-2}$ [$2.3x10^{-3}$, $5.5x10^{-1}$] | 65 |
| | **very rare** | **16.68** | **16.92 [16.86, 16.98]** | **0.986 [0.982, 0.989]** | **$2.9x10^{-8}$ [$2.9x10^{-11}$, $1.1x10^{-5}$]** | **9996** |
| | **ultra-rare** | **2.54** | **2.19 [2.17, 2.20]** | **1.162 [1.152, 1.169]** | **$8.4x10^{-81}$ [$5.3x10^{-85}$, $5.3x10^{-76}$]** | **10000** |

**S3 Table. VIKING vs LBC: INDEL load comparison in coding and coding related regions (alleles per individual per 1Mb).** Very common: variants with MAF > 10% in Non-Finnish Europeans (NFE, gnomADg, n=7,509); common: 5% < $MAF_{NFE}$ ≤ 10%; rare: 1% < $MAF_{NFE}$ ≤ 5%; very rare: $MAF_{NFE}$ ≤ 1%; ultra-rare: not observed in any gnomADg individual (n=15,496). Median number and 95% CI of LBC alleles (forth column) for each frequency class is computed based on 10,000 random subsets (n = 269, matching VIKING size); last two columns represent the median p-value (and 95% CI) and the number of tests with p-value smaller than the Bonferroni corrected threshold. To annotate the number of variants in a frequency class as significantly different (shown in bold), we required at least 95% of the 10,000 subsets to have p-value ≤ $8x10^{-4}$ (Bonferroni corrected) and no overlap between the 95% CI for the LBC and VIKING median values. Region annotation (5'UTR, Exon, Intron, 3'UTR, ncRNA) is based on Ensembl data (GRCh37.p13, Ensembl Genes 92) for the mappable sections of the 22 autosomal chromosomes; the remaining 1.1Gb of the mappable regions in the reference human genome is annotated as "non-coding".

| Region | gnomAD Frequency Class | VIKING median | LBC 10k subsets median & 95%CI | VIKING/LBC ratio median & 95%CI | Wilcoxon rank sum test | |
|---|---|---|---|---|---|---|
| | | | | | *p* : median & 95% CI | number of tests with $p \le 8x10^{-4}$ |
| **5'UTR** | **very common** | **102.08** | **100.90 [100.57, 101.22]** | **1.012 [1.009, 1.015]** | **$1.2x10^{-8}$ [$7.4x10^{-12}$, $4.9x10^{-6}$]** | **10000** |
| | common | 3.98 | 3.77 [3.66, 3.88] | 1.057 [1.028, 1.088] | $2.5x10^{-3}$ [$2.6x10^{-5}$, $6.7x10^{-2}$] | 2990 |
| | rare | 3.12 | 3.02 [2.91, 3.12] | 1.036 [1.000, 1.074] | $1.7x10^{-2}$ [$3.6x10^{-4}$, $2.5x10^{-1}$] | 581 |
| | very rare | 1.72 | 1.72 [1.72, 1.83] | 1.000 [0.941, 1.000] | $4.8x10^{-1}$ [$5.1x10^{-2}$, $9.7x10^{-1}$] | 0 |
| | ultra-rare | 0.22 | 0.11 [0.11, 0.22] | 2.000 [1.000, 2.000] | $3.7x10^{-5}$ [$1.5x10^{-7}$, $2.7x10^{-3}$] | 9086 |
| **Exon** | very common | 12.51 | 12.44 [12.41, 12.54] | 1.005 [0.997, 1.008] | $3.6x10^{-1}$ [$2.7x10^{-2}$, $9.6x10^{-1}$] | 1 |
| | common | 0.60 | 0.63 [0.60, 0.67] | 0.947 [0.900, 1.000] | $6.7x10^{-1}$ [$1.5x10^{-1}$, $9.8x10^{-1}$] | 0 |
| | rare | 0.57 | 0.53 [0.53, 0.57] | 1.062 [1.000, 1.062] | $9.9x10^{-2}$ [$5.1x10^{-3}$, $6.1x10^{-1}$] | 19 |
| | very rare | 0.57 | 0.60 [0.60, 0.63] | 0.944 [0.895, 0.944] | $5.3x10^{-2}$ [$2.1x10^{-3}$, $4.1x10^{-1}$] | 65 |
| | ultra-rare | 0.10 | 0.10 [0.10, 0.10] | 1.000 [1.000, 1.0000] | $1.1x10^{-1}$ [$5.3x10^{-3}$, $6.7x10^{-1}$] | 20 |
| **Intron** | **very common** | **154.31** | **153.41 [153.32, 153.49]** | **1.006 [1.005, 1.006]** | **$1.3x10^{-41}$ [$4.2x10^{-47}$, $2.4x10^{-36}$]** | **10000** |
| | **common** | **5.64** | **5.56 [5.54, 5.58]** | **1.014 [1.010, 1.018]** | **$2.3x10^{-8}$ [$1.2x10^{-11}$, $1.3x10^{-5}$]** | **9999** |
| | rare | 4.23 | 4.19 [4.17, 4.21] | 1.009 [1.005, 1.013] | $4.4x10^{-4}$ [$2.4x10^{-6}$, $2.2x10^{-2}$] | 6058 |
| | **very rare** | **1.86** | **1.90 [1.89, 1.91]** | **0.978 [0.974, 0.983]** | **$6.0x10^{-10}$ [$1.9x10^{-13}$, $5.6x10^{-7}$]** | **10000** |
| | **ultra-rare** | **0.25** | **0.20 [0.20, 0.21]** | **1.228 [1.215, 1.249]** | **$1.4x10^{-74}$ [$9.8x10^{-79}$, $3.8x10^{-70}$]** | **10000** |
| **3'UTR** | **very common** | **160.43** | **159.63 [159.41, 159.88]** | **1.005 [1.003, 1.006]** | **$3.4x10^{-8}$ [$1.5x10^{-11}$, $1.6x10^{-5}$]** | **9997** |
| | common | 6.27 | 6.17 [6.09, 6.20] | 1.018 [1.012, 1.03] | $1.9x10^{-2}$ [$5.6x10^{-4}$, $2.3x10^{-1}$] | 376 |
| | rare | 4.79 | 4.82 [4.75, 4.90] | 0.992 [0.978, 1.008] | $6.8x10^{-1}$ [$1.7x10^{-1}$, $9.8x10^{-1}$] | 0 |
| | very rare | 2.25 | 2.39 [2.36, 2.43] | 0.939 [0.925, 0.954] | $9.8x10^{-5}$ [$3.6x10^{-7}$, $6.6x10^{-3}$] | 8218 |
| | ultra-rare | 0.29 | 0.25 [0.25, 0.25] | 1.143 [1.143, 1.143] | $1.3x10^{-3}$ [$8.5x10^{-6}$, $4.6x10^{-2}$] | 4065 |
| **ncRNA** | very common | 149.13 | 148.58 [148.17, 149.00] | 1.004 [1.001, 1.006] | $1.6x10^{-1}$ [$8.8x10^{-3}$, $8.3x10^{-1}$] | 15 |
| | common | 6.04 | 5.76 [5.63, 5.90] | 1.048 [1.023, 1.073] | $1.3x10^{-3}$ [$1.2x10^{-5}$, $3.6x10^{-2}$] | 4150 |
| | rare | 4.12 | 3.98 [3.84, 4.12] | 1.034 [1.000, 1.071] | $5.4x10^{-1}$ [$7.8x10^{-2}$, $9.8x10^{-1}$] | 0 |
| | very rare | 2.33 | 2.33 [2.33, 2.47] | 1.000 [0.944, 1.000] | $5.5x10^{-1}$ [$7.9x10^{-2}$, $9.8x10^{-1}$] | 0 |
| | ultra-rare | 0.14 | 0.14 [0.14, 0.14] | 1.000 [1.000, 1.000] | $1.6x10^{-2}$ [$4.2x10^{-4}$, $2.0x10^{-1}$] | 496 |
| **non-coding** | **very common** | **171.56** | **170.64 [170.54, 170.74]** | **1.005 [1.005, 1.006]** | **$3.8x10^{-38}$ [$2.8x10^{-43}$, $4.4x10^{-33}$]** | **10000** |
| | common | 6.11 | 6.06 [6.03, 6.08] | 1.009 [1.005, 1.013] | $3.4x10^{-5}$ [$1.2x10^{-7}$, $2.8x10^{-3}$] | 9113 |
| | rare | 4.37 | 4.35 [4.33, 4.36] | 1.006 [1.001, 1.010] | $2.4x10^{-2}$ [$5.5x10^{-4}$, $3.0x10^{-1}$] | 366 |
| | **very rare** | **2.06** | **2.10 [2.09, 2.11]** | **0.980 [0.977, 0.984]** | **$1.7x10^{-9}$ [$1.7x10^{-12}$, $7.5x10^{-7}$]** | **10000** |
| | **ultra-rare** | **0.25** | **0.20 [0.20, 0.20]** | **1.214 [1.203, 1.230]** | **$6.5x10^{-71}$ [$1.9x10^{-75}$, $4.3x10^{-66}$]** | **10000** |

**S4 Table. VIKING vs LBC: ultra-rare SNP load comparison in different chromatin states (alleles per individual per 1Mb).** To annotate the number of variants in a state/cell type class as significantly different, we required at least 95% of the 10,000 subsets to have p-value ≤ $2\times10^{-4}$ (Bonferroni corrected) and no overlap between the 95% CI for the LBC and VIKING median values; VIKING is enriched for ultra-rare SNPs in all considered states/cell types.

| State | Cell Type | VIKING median | LBC 10k subsets median & 95%CI | VIKING/LBC ratio median & 95%CI | Wilcoxon rank sum test: $p$ : median & 95% CI | Wilcoxon rank sum test: number of tests with $p \leq 2\times10^{-4}$ |
|---|---|---|---|---|---|---|
| Promoter | Gm12878 | 2.379 | 2.005 [1.961, 2.049] | 1.187 [1.161, 1.213] | $7.1\times10^{-39}$ [$1.3\times10^{-44}$, $3.0\times10^{-33}$] | 10000 |
| | H1hesc | 2.391 | 2.000 [1.978, 2.043] | 1.196 [1.170, 1.209] | $4.8\times10^{-50}$ [$2.3\times10^{-55}$, $9.8\times10^{-45}$] | 10000 |
| | Hepg2 | 2.265 | 1.928 [1.909, 1.965] | 1.175 [1.152, 1.186] | $4.6\times10^{-44}$ [$2.2\times10^{-49}$, $8.1\times10^{-39}$] | 10000 |
| | Hmec | 2.378 | 1.987 [1.957, 2.017] | 1.197 [1.179, 1.215] | $2.0\times10^{-39}$ [$6.4\times10^{-45}$, $4.9\times10^{-34}$] | 10000 |
| | Hsmm | 2.372 | 1.986 [1.958, 2.041] | 1.194 [1.162, 1.211] | $2.2\times10^{-43}$ [$9.7\times10^{-49}$, $4.4\times10^{-38}$] | 10000 |
| | Huvec | 2.378 | 2.009 [1.942, 2.043] | 1.183 [1.164, 1.224] | $1.4\times10^{-41}$ [$9.0\times10^{-47}$, $2.0\times10^{-36}$] | 10000 |
| | K562 | 2.208 | 1.897 [1.869, 1.954] | 1.164 [1.130, 1.182] | $3.0\times10^{-32}$ [$2.2\times10^{-37}$, $2.3\times10^{-27}$] | 10000 |
| | Nhek | 2.453 | 2.040 [2.012, 2.095] | 1.203 [1.171, 1.219] | $5.7\times10^{-40}$ [$1.1\times10^{-45}$, $2.1\times10^{-34}$] | 10000 |
| | Nhlf | 2.352 | 2.001 [1.974, 2.028] | 1.176 [1.160, 1.192] | $5.8\times10^{-45}$ [$3.0\times10^{-50}$, $1.1\times10^{-39}$] | 10000 |
| | Union | 2.302 | 1.951 [1.923, 1.970] | 1.180 [1.168, 1.197] | $5.9\times10^{-56}$ [$2.0\times10^{-61}$, $1.6\times10^{-50}$] | 10000 |
| Enhancer | Gm12878 | 2.091 | 1.760 [1.745, 1.790] | 1.188 [1.168, 1.198] | $2.2\times10^{-53}$ [$2.0\times10^{-59}$, $2.8\times10^{-47}$] | 10000 |
| | H1hesc | 2.189 | 1.889 [1.862, 1.907] | 1.159 [1.148, 1.176] | $7.2\times10^{-54}$ [$4.4\times10^{-59}$, $1.3\times10^{-48}$] | 10000 |
| | Hepg2 | 1.964 | 1.685 [1.666, 1.714] | 1.166 [1.146, 1.179] | $1.2\times10^{-53}$ [$6.6\times10^{-59}$, $2.3\times10^{-48}$] | 10000 |
| | Hmec | 2.085 | 1.806 [1.787, 1.818] | 1.155 [1.147, 1.167] | $9.1\times10^{-65}$ [$1.4\times10^{-69}$, $9.9\times10^{-60}$] | 10000 |
| | Hsmm | 2.153 | 1.853 [1.839, 1.875] | 1.161 [1.148, 1.171] | $1.7\times10^{-61}$ [$4.2\times10^{-66}$, $9.8\times10^{-57}$] | 10000 |
| | Huvec | 2.117 | 1.801 [1.777, 1.825] | 1.175 [1.160, 1.191] | $1.0\times10^{-64}$ [$2.5\times10^{-69}$, $6.2\times10^{-60}$] | 10000 |
| | K562 | 1.896 | 1.621 [1.598, 1.636] | 1.170 [1.159, 1.186] | $1.3\times10^{-64}$ [$2.3\times10^{-69}$, $1.1\times10^{-59}$] | 10000 |
| | Nhek | 2.069 | 1.800 [1.779, 1.814] | 1.150 [1.141, 1.163] | $1.4\times10^{-62}$ [$1.0\times10^{-67}$, $2.5\times10^{-57}$] | 10000 |
| | Nhlf | 2.139 | 1.854 [1.831, 1.878] | 1.153 [1.139, 1.168] | $4.7\times10^{-53}$ [$1.3\times10^{-58}$, $1.8\times10^{-47}$] | 10000 |
| | Union | 2.056 | 1.773 [1.761, 1.788] | 1.160 [1.150, 1.168] | $2.8\times10^{-78}$ [$1.1\times10^{-82}$, $1.7\times10^{-73}$] | 10000 |
| Insulator | Gm12878 | 2.136 | 1.869 [1.802, 1.935] | 1.143 [1.103, 1.185] | $9.4\times10^{-12}$ [$1.9\times10^{-15}$, $1.3\times10^{-8}$] | 10000 |
| | H1hesc | 2.106 | 1.922 [1.877, 1.968] | 1.095 [1.070, 1.122] | $2.8\times10^{-16}$ [$9.8\times10^{-21}$, $1.9\times10^{-12}$] | 10000 |
| | Hepg2 | 2.150 | 1.978 [1.892, 1.978] | 1.087 [1.087, 1.136] | $1.8\times10^{-6}$ [$1.3\times10^{-9}$, $2.8\times10^{-4}$] | 9661 |
| | Hmec | 2.072 | 1.913 [1.833, 1.913] | 1.083 [1.083, 1.130] | $7.6\times10^{-10}$ [$1.9\times10^{-13}$, $7.7\times10^{-7}$] | 10000 |
| | Hsmm | 2.252 | 1.931 [1.931, 1.995] | 1.167 [1.129, 1.167] | $3.9\times10^{-12}$ [$5.8\times10^{-16}$, $5.8\times10^{-9}$] | 10000 |
| | Huvec | 2.191 | 1.859 [1.792, 1.925] | 1.179 [1.138, 1.222] | $4.6\times10^{-16}$ [$1.7\times10^{-20}$, $4.3\times10^{-12}$] | 10000 |
| | K562 | 2.138 | 1.877 [1.825, 1.929] | 1.139 [1.108, 1.171] | $4.5\times10^{-20}$ [$5.3\times10^{-25}$, $9.8\times10^{-16}$] | 10000 |
| | Nhek | 2.189 | 1.876 [1.824, 1.928] | 1.167 [1.135, 1.200] | $7.6\times10^{-20}$ [$1.5\times10^{-24}$, $1.4\times10^{-15}$] | 10000 |
| | Nhlf | 2.097 | 1.830 [1.792, 1.869] | 1.146 [1.122, 1.170] | $1.4\times10^{-26}$ [$1.2\times10^{-31}$, $9.0\times10^{-22}$] | 10000 |
| | Union | 2.165 | 1.895 [1.861, 1.911] | 1.143 [1.133, 1.164] | $1.4\times10^{-38}$ [$6.2\times10^{-44}$, $1.4\times10^{-33}$] | 10000 |
| Transcription | Gm12878 | 1.895 | 1.650 [1.637, 1.665] | 1.149 [1.138, 1.158] | $1.2\times10^{-66}$ [$7.9\times10^{-72}$, $3.2\times10^{-61}$] | 10000 |
| | H1hesc | 1.927 | 1.670 [1.655, 1.682] | 1.154 [1.146, 1.164] | $5.9\times10^{-76}$ [$1.6\times10^{-80}$, $7.5\times10^{-71}$] | 10000 |
| | Hepg2 | 1.858 | 1.615 [1.602, 1.626] | 1.151 [1.143, 1.160] | $1.2\times10^{-73}$ [$2.5\times10^{-78}$, $1.4\times10^{-68}$] | 10000 |
| | Hmec | 1.893 | 1.646 [1.633, 1.658] | 1.150 [1.142, 1.159] | $3.4\times10^{-73}$ [$5.4\times10^{-78}$, $4.0\times10^{-68}$] | 10000 |
| | Hsmm | 1.887 | 1.637 [1.624, 1.649] | 1.153 [1.144, 1.162] | $5.6\times10^{-75}$ [$1.4\times10^{-79}$, $5.5\times10^{-70}$] | 10000 |
| | Huvec | 1.873 | 1.633 [1.618, 1.650] | 1.147 [1.135, 1.157] | $3.9\times10^{-70}$ [$5.7\times10^{-75}$, $5.7\times10^{-65}$] | 10000 |
| | K562 | 1.816 | 1.574 [1.564, 1.588] | 1.154 [1.144, 1.161] | $2.0\times10^{-74}$ [$4.0\times10^{-79}$, $3.0\times10^{-69}$] | 10000 |
| | Nhek | 1.864 | 1.613 [1.604, 1.627] | 1.156 [1.146, 1.162] | $1.3\times10^{-73}$ [$3.4\times10^{-78}$, $1.4\times10^{-68}$] | 10000 |
| | Nhlf | 1.878 | 1.634 [1.620, 1.649] | 1.149 [1.139, 1.159] | $1.2\times10^{-73}$ [$2.8\times10^{-78}$, $1.3\times10^{-68}$] | 10000 |
| | Union | 1.911 | 1.655 [1.643, 1.668] | 1.155 [1.146, 1.163] | $9.0\times10^{-81}$ [$6.6\times10^{-85}$, $4.0\times10^{-76}$] | 10000 |
| Repressed | Gm12878 | 2.062 | 1.803 [1.769, 1.825] | 1.144 [1.130, 1.166] | $1.3\times10^{-47}$ [$3.5\times10^{-53}$, $3.8\times10^{-42}$] | 10000 |
| | H1hesc | 2.076 | 1.810 [1.757, 1.837] | 1.147 [1.130, 1.182] | $3.1\times10^{-25}$ [$6.5\times10^{-31}$, $3.8\times10^{-20}$] | 10000 |
| | Hepg2 | 2.202 | 1.908 [1.884, 1.925] | 1.154 [1.144, 1.169] | $7.8\times10^{-50}$ [$1.6\times10^{-55}$, $3.0\times10^{-44}$] | 10000 |
| | Hmec | 2.223 | 1.927 [1.894, 1.960] | 1.154 [1.134, 1.174] | $2.3\times10^{-35}$ [$3.8\times10^{-41}$, $7.5\times10^{-30}$] | 10000 |
| | Hsmm | 2.075 | 1.795 [1.778, 1.821] | 1.156 [1.140, 1.167] | $1.8\times10^{-47}$ [$1.2\times10^{-52}$, $3.4\times10^{-42}$] | 10000 |
| | Huvec | 2.070 | 1.774 [1.748, 1.799] | 1.167 [1.150, 1.184] | $3.6\times10^{-51}$ [$5.4\times10^{-57}$, $2.0\times10^{-45}$] | 10000 |
| | K562 | 2.088 | 1.818 [1.800, 1.837] | 1.148 [1.137, 1.160] | $5.3\times10^{-64}$ [$2.6\times10^{-69}$, $1.4\times10^{-58}$] | 10000 |
| | Nhek | 2.206 | 1.928 [1.901, 1.949] | 1.144 [1.132, 1.161] | $2.7\times10^{-50}$ [$4.8\times10^{-56}$, $8.0\times10^{-45}$] | 10000 |
| | Nhlf | 2.007 | 1.756 [1.729, 1.772] | 1.143 [1.133, 1.160] | $4.3\times10^{-56}$ [$7.8\times10^{-62}$, $3.1\times10^{-50}$] | 10000 |
| | Union | 2.074 | 1.787 [1.773, 1.802] | 1.161 [1.151, 1.170] | $1.8\times10^{-76}$ [$6.4\times10^{-81}$, $1.0\times10^{-71}$] | 10000 |
| Heterochromatin | Gm12878 | 1.896 | 1.631 [1.620, 1.641] | 1.162 [1.155, 1.171] | $3.1\times10^{-82}$ [$3.1\times10^{-86}$, $1.4\times10^{-77}$] | 10000 |
| | H1hesc | 1.890 | 1.623 [1.611, 1.635] | 1.164 [1.156, 1.173] | $1.2\times10^{-81}$ [$1.3\times10^{-85}$, $5.1\times10^{-77}$] | 10000 |
| | Hepg2 | 1.908 | 1.640 [1.627, 1.654] | 1.163 [1.154, 1.172] | $8.1\times10^{-82}$ [$8.6\times10^{-86}$, $3.8\times10^{-77}$] | 10000 |
| | Hmec | 1.898 | 1.631 [1.619, 1.642] | 1.163 [1.156, 1.172] | $5.9\times10^{-82}$ [$5.4\times10^{-86}$, $3.2\times10^{-77}$] | 10000 |
| | Hsmm | 1.895 | 1.627 [1.615, 1.638] | 1.165 [1.157, 1.174] | $1.1\times10^{-81}$ [$1.2\times10^{-85}$, $5.9\times10^{-77}$] | 10000 |
| | Huvec | 1.892 | 1.629 [1.618, 1.641] | 1.162 [1.153, 1.169] | $5.4\times10^{-82}$ [$5.2\times10^{-86}$, $2.5\times10^{-77}$] | 10000 |
| | K562 | 1.920 | 1.652 [1.640, 1.664] | 1.162 [1.154, 1.171] | $4.3\times10^{-81}$ [$3.1\times10^{-85}$, $2.4\times10^{-76}$] | 10000 |
| | Nhek | 1.893 | 1.626 [1.610, 1.637] | 1.164 [1.156, 1.176] | $5.7\times10^{-82}$ [$5.8\times10^{-86}$, $2.5\times10^{-77}$] | 10000 |
| | Nhlf | 1.897 | 1.627 [1.617, 1.639] | 1.166 [1.157, 1.173] | $5.6\times10^{-82}$ [$5.0\times10^{-86}$, $3.4\times10^{-77}$] | 10000 |
| | Union | 1.918 | 1.650 [1.638, 1.662] | 1.163 [1.154, 1.171] | $3.8\times10^{-82}$ [$3.5\times10^{-86}$, $1.6\times10^{-77}$] | 10000 |

**S5 Table. VIKING vs LBC: ultra-rare INDEL load comparison in different chromatin states (alleles per individual per 1Mb).** To annotate the number of variants in a state/cell type class as significantly different, we required at least 95% of the 10,000 subsets to have p-value ≤ $2x10^{-4}$ (Bonferroni corrected) and no overlap between the 95% CI for the LBC and VIKING median values; similar to ultra-rare SNPs, VIKING is enriched for ultra-rare INDELs in almost all considered states/cell types, except the Insulator chromatin state (shown in grey).

| State | Cell Type | VIKING median | LBC 10k subsets median & 95%CI | VIKING/LBC ratio median & 95%CI | Wilcoxon rank sum test | |
|---|---|---|---|---|---|---|
| | | | | | *p* : median & 95% CI | number of tests with *p* ≤ $2x10^{-4}$ |
| Promoter | Gm12878 | 0.242 | 0.198 [0.198, 0.220] | 1.222 [1.100, 1.222] | $3.0x10^{-10}$ [$1.2x10^{-13}$, $2.9x10^{-7}$] | 9999 |
| | H1hesc | 0.239 | 0.196 [0.174, 0.196] | 1.222 [1.222, 1.375] | $7.3x10^{-10}$ [$3.2x10^{-13}$, $5.8x10^{-7}$] | 10000 |
| | Hepg2 | 0.243 | 0.187 [0.168, 0.187] | 1.300 [1.300, 1.444] | $3.6x10^{-20}$ [$2.9x10^{-24}$, $2.7x10^{-16}$] | 10000 |
| | Hmec | 0.241 | 0.181 [0.181, 0.211] | 1.333 [1.143, 1.333] | $1.9x10^{-10}$ [$1.4x10^{-13}$, $1.0x10^{-7}$] | 10000 |
| | Hsmm | 0.248 | 0.193 [0.165, 0.193] | 1.286 [1.286, 1.500] | $3.8x10^{-15}$ [$8.0x10^{-19}$, $8.1x10^{-12}$] | 10000 |
| | Huvec | 0.234 | 0.201 [0.167, 0.201] | 1.167 [1.167, 1.400] | $2.1x10^{-9}$ [$1.7x10^{-12}$, $1.0x10^{-6}$] | 10000 |
| | K562 | 0.255 | 0.198 [0.170, 0.198] | 1.286 [1.286, 1.500] | $1.1x10^{-14}$ [$1.8x10^{-18}$, $3.7x10^{-11}$] | 10000 |
| | Nhek | 0.248 | 0.193 [0.193, 0.193] | 1.286 [1.286, 1.286] | $1.0x10^{-10}$ [$3.9x10^{-14}$, $9.1x10^{-8}$] | 10000 |
| | Nhlf | 0.243 | 0.189 [0.189, 0.216] | 1.286 [1.125, 1.286] | $1.4x10^{-8}$ [$1.4x10^{-11}$, $5.7x10^{-6}$] | 9993 |
| Enhancer | Gm12878 | 0.218 | 0.173 [0.165, 0.181] | 1.261 [1.208, 1.318] | $3.3x10^{-19}$ [$7.4x10^{-24}$, $5.7x10^{-15}$] | 10000 |
| | H1hesc | 0.200 | 0.163 [0.163, 0.173] | 1.222 [1.158, 1.222] | $1.3x10^{-17}$ [$5.9x10^{-22}$, $8.9x10^{-14}$] | 10000 |
| | Hepg2 | 0.183 | 0.154 [0.144, 0.154] | 1.187 [1.187, 1.267] | $3.7x10^{-16}$ [$1.8x10^{-20}$, $2.8x10^{-12}$] | 10000 |
| | Hmec | 0.186 | 0.161 [0.155, 0.168] | 1.154 [1.111, 1.200] | $3.5x10^{-20}$ [$6.9x10^{-25}$, $5.7x10^{-16}$] | 10000 |
| | Hsmm | 0.197 | 0.161 [0.161, 0.168] | 1.227 [1.174, 1.227] | $1.2x10^{-25}$ [$3.7x10^{-30}$, $2.4x10^{-21}$] | 10000 |
| | Huvec | 0.213 | 0.166 [0.166, 0.174] | 1.286 [1.227, 1.286] | $2.4x10^{-30}$ [$2.7x10^{-35}$, $1.4x10^{-25}$] | 10000 |
| | K562 | 0.171 | 0.149 [0.141, 0.149] | 1.150 [1.150, 1.211] | $4.6x10^{-19}$ [$1.6x10^{-23}$, $3.5x10^{-15}$] | 10000 |
| | Nhek | 0.191 | 0.163 [0.156, 0.170] | 1.174 [1.125, 1.227] | $8.6x10^{-23}$ [$1.3x10^{-27}$, $1.6x10^{-18}$] | 10000 |
| | Nhlf | 0.213 | 0.166 [0.158, 0.166] | 1.286 [1.286, 1.350] | $9.2x10^{-30}$ [$2.8x10^{-34}$, $1.7x10^{-25}$] | 10000 |
| Insulator | Gm12878 | 0.200 | 0.133 [0.133, 0.200] | 1.500 [1.000, 1.500] | $4.2x10^{-3}$ [$5.3x10^{-5}$, $9.3x10^{-2}$] | 782 |
| | H1hesc | 0.183 | 0.183 [0.137, 0.183] | 1.000 [1.000, 1.333] | $2.7x10^{-3}$ [$3.3x10^{-5}$, $6.4x10^{-2}$] | 1148 |
| | Hepg2 | 0.172 | 0.172 [0.086, 0.172] | 1.000 [1.000, 2.000] | $7.4x10^{-2}$ [$3.2x10^{-3}$, $5.4x10^{-1}$] | 3 |
| | Hmec | 0.159 | 0.159 [0.159, 0.159] | 1.000 [1.000, 1.000] | $5.8x10^{-2}$ [$1.6x10^{-3}$, $4.8x10^{-1}$] | 28 |
| | Hsmm | 0.193 | 0.193 [0.193, 0.193] | 1.000 [1.000, 1.000] | $2.8x10^{-2}$ [$7.6x10^{-4}$, $3.1x10^{-1}$] | 50 |
| | Huvec | 0.199 | 0.133 [0.133, 0.199] | 1.500 [1.000, 1.500] | $1.2x10^{-1}$ [$5.4x10^{-3}$, $7.3x10^{-1}$] | 1 |
| | K562 | 0.156 | 0.156 [0.156, 0.156] | 1.000 [1.000, 1.000] | $2.9x10^{-1}$ [$2.7x10^{-2}$, $9.2x10^{-1}$] | 0 |
| | Nhek | 0.208 | 0.156 [0.156, 0.156] | 1.333 [1.333, 1.333] | $7.6x10^{-5}$ [$2.7x10^{-7}$, $5.8x10^{-3}$] | 6425 |
| | Nhlf | 0.153 | 0.153 [0.153, 0.153] | 1.000 [1.000, 1.000] | $7.7x10^{-2}$ [$2.9x10^{-3}$, $6.0x10^{-1}$] | 9 |
| Transcription | Gm12878 | 0.211 | 0.173 [0.169, 0.175] | 1.222 [1.207, 1.253] | $5.2x10^{-50}$ [$8.5x10^{-55}$, $3.3x10^{-45}$] | 10000 |
| | H1hesc | 0.211 | 0.171 [0.168, 0.174] | 1.233 [1.210, 1.257] | $1.2x10^{-64}$ [$7.6x10^{-69}$, $2.6x10^{-60}$] | 10000 |
| | Hepg2 | 0.209 | 0.171 [0.168, 0.173] | 1.220 [1.207, 1.247] | $1.7x10^{-58}$ [$4.3x10^{-63}$, $7.9x10^{-54}$] | 10000 |
| | Hmec | 0.210 | 0.168 [0.164, 0.170] | 1.253 [1.239, 1.282] | $3.6x10^{-61}$ [$7.7x10^{-66}$, $2.9x10^{-56}$] | 10000 |
| | Hsmm | 0.205 | 0.168 [0.165, 0.171] | 1.220 [1.198, 1.243] | $3.2x10^{-61}$ [$4.0x10^{-66}$, $2.4x10^{-56}$] | 10000 |
| | Huvec | 0.206 | 0.170 [0.168, 0.175] | 1.210 [1.181, 1.225] | $2.1x10^{-53}$ [$1.0x10^{-58}$, $4.8x10^{-48}$] | 10000 |
| | K562 | 0.206 | 0.168 [0.166, 0.172] | 1.229 [1.200, 1.244] | $4.2x10^{-56}$ [$3.7x10^{-61}$, $4.0x10^{-51}$] | 10000 |
| | Nhek | 0.205 | 0.168 [0.164, 0.170] | 1.218 [1.205, 1.247] | $1.2x10^{-57}$ [$9.4x10^{-63}$, $1.9x10^{-52}$] | 10000 |
| | Nhlf | 0.207 | 0.171 [0.169, 0.174] | 1.216 [1.189, 1.230] | $1.1x10^{-56}$ [$6.4x10^{-62}$, $2.2x10^{-51}$] | 10000 |
| Repressed | Gm12878 | 0.203 | 0.146 [0.146, 0.158] | 1.385 [1.286, 1.385] | $3.0x10^{-25}$ [$1.5x10^{-29}$, $4.9x10^{-21}$] | 10000 |
| | H1hesc | 0.213 | 0.186 [0.160, 0.186] | 1.143 [1.143, 1.333] | $9.0x10^{-10}$ [$3.8x10^{-13}$, $7.7x10^{-7}$] | 10000 |
| | Hepg2 | 0.204 | 0.171 [0.163, 0.171] | 1.190 [1.190, 1.250] | $1.3x10^{-24}$ [$1.5x10^{-29}$, $5.2x10^{-20}$] | 10000 |
| | Hmec | 0.198 | 0.165 [0.148, 0.165] | 1.200 [1.200, 1.333] | $1.2x10^{-13}$ [$1.4x10^{-17}$, $3.3x10^{-10}$] | 10000 |
| | Hsmm | 0.178 | 0.144 [0.136, 0.152] | 1.235 [1.167, 1.312] | $1.1x10^{-20}$ [$3.7x10^{-25}$, $1.9x10^{-16}$] | 10000 |
| | Huvec | 0.194 | 0.163 [0.158, 0.173] | 1.187 [1.118, 1.226] | $6.5x10^{-17}$ [$2.6x10^{-21}$, $5.8x10^{-13}$] | 10000 |
| | K562 | 0.196 | 0.160 [0.155, 0.164] | 1.229 [1.194, 1.265] | $7.6x10^{-29}$ [$1.2x10^{-33}$, $2.9x10^{-24}$] | 10000 |
| | Nhek | 0.194 | 0.153 [0.146, 0.160] | 1.273 [1.217, 1.333] | $9.1x10^{-25}$ [$2.0x10^{-29}$, $2.5x10^{-20}$] | 10000 |
| | Nhlf | 0.181 | 0.144 [0.139, 0.149] | 1.259 [1.214, 1.308] | $2.1x10^{-31}$ [$3.4x10^{-36}$, $8.2x10^{-27}$] | 10000 |
| Heterochromatin | Gm12878 | 0.175 | 0.143 [0.142, 0.145] | 1.225 [1.208, 1.239] | $5.7x10^{-79}$ [$1.9x10^{-82}$, $7.2x10^{-75}$] | 10000 |
| | H1hesc | 0.174 | 0.142 [0.141, 0.144] | 1.220 [1.206, 1.234] | $2.8x10^{-78}$ [$4.9x10^{-82}$, $7.1x10^{-74}$] | 10000 |
| | Hepg2 | 0.176 | 0.144 [0.142, 0.145] | 1.221 [1.212, 1.240] | $3.9x10^{-77}$ [$4.9x10^{-81}$, $9.3x10^{-73}$] | 10000 |
| | Hmec | 0.178 | 0.145 [0.144, 0.147] | 1.223 [1.209, 1.236] | $4.6x10^{-78}$ [$9.0x10^{-82}$, $8.3x10^{-74}$] | 10000 |
| | Hsmm | 0.175 | 0.144 [0.141, 0.145] | 1.219 [1.209, 1.239] | $2.6x10^{-76}$ [$2.2x10^{-80}$, $1.1x10^{-71}$] | 10000 |
| | Huvec | 0.176 | 0.142 [0.141, 0.143] | 1.239 [1.225, 1.249] | $4.6x10^{-80}$ [$1.5x10^{-83}$, $6.0x10^{-76}$] | 10000 |
| | K562 | 0.177 | 0.144 [0.143, 0.146] | 1.228 [1.214, 1.238] | $2.0x10^{-76}$ [$2.1x10^{-80}$, $6.0x10^{-72}$] | 10000 |
| | Nhek | 0.179 | 0.145 [0.143, 0.147] | 1.232 [1.218, 1.246] | $1.0x10^{-77}$ [$1.6x10^{-81}$, $2.0x10^{-73}$] | 10000 |
| | Nhlf | 0.176 | 0.144 [0.143, 0.146] | 1.222 [1.208, 1.236] | $1.2x10^{-76}$ [$1.9x10^{-80}$, $3.3x10^{-72}$] | 10000 |

**S6 Table. VIKING vs LBC: known SNP load comparison in different chromatin states (alleles per individual per 1Mb).** To annotate the number of variants in a state/cell type class as significantly different, we required at least 95% of the 10,000 subsets to have p-value ≤ $2x10^{-4}$ (Bonferroni corrected) and no overlap between the 95% CI for the LBC and VIKING median values; there is no significant difference between the two cohorts for known SNPs in any of the considered states/cell types.

| State | Cell Type | VIKING median | LBC 10k subsets median & 95%CI | VIKING/LBC ratio median & 95%CI | Wilcoxon rank sum test | |
|---|---|---|---|---|---|---|
| | | | | | *p* : median & 95% CI | number of tests with *p* ≤ $2x10^{-4}$ |
| **Promoter** | Gm12878 | 1116 | 1116 [1115, 1117] | 1.000 [0.999, 1.001] | $5.9x10^{-1}$ [$9.1x10^{-2}$, $9.8x10^{-1}$] | 0 |
| | H1hesc | 1148 | 1147 [1146, 1147] | 1.001 [1.001, 1.002] | $8.1x10^{-3}$ [$1.1x10^{-4}$, $1.5x10^{-1}$] | 421 |
| | Hepg2 | 1205 | 1204 [1203, 1205] | 1.001 [1.000, 1.001] | $9.4x10^{-2}$ [$3.1x10^{-3}$, $6.9x10^{-1}$] | 9 |
| | Hmec | 1109 | 1106 [1105, 1108] | 1.002 [1.001, 1.003] | $1.6x10^{-4}$ [$5.8x10^{-7}$, $1.0x10^{-2}$] | 5327 |
| | Hsmm | 1131 | 1128 [1127, 1129] | 1.002 [1.001, 1.003] | $4.3x10^{-5}$ [$1.1x10^{-7}$, $4.5x10^{-3}$] | 7205 |
| | Huvec | 1127 | 1125 [1124, 1126] | 1.002 [1.001, 1.003] | $6.8x10^{-4}$ [$5.0x10^{-6}$, $2.6x10^{-2}$] | 3011 |
| | K562 | 1133 | 1131 [1130, 1132] | 1.002 [1.001, 1.003] | $1.0x10^{-2}$ [$1.3x10^{-4}$, $1.9x10^{-1}$] | 376 |
| | Nhek | 1167 | 1166 [1165, 1167] | 1.001 [1.000, 1.002] | $1.5x10^{-2}$ [$2.2x10^{-4}$, $2.3x10^{-1}$] | 227 |
| | Nhlf | 1143 | 1141 [1140, 1142] | 1.001 [1.000, 1.002] | $7.7x10^{-3}$ [$9.2x10^{-5}$, $1.7x10^{-1}$] | 477 |
| **Enhancer** | Gm12878 | 1253 | 1252 [1251, 1253] | 1.001 [1.000, 1.002] | $1.9x10^{-1}$ [$1.2x10^{-2}$, $8.3x10^{-1}$] | 0 |
| | H1hesc | 1403 | 1403 [1402, 1403] | 1.000 [1.000, 1.001] | $5.6x10^{-1}$ [$8.1x10^{-2}$, $9.8x10^{-1}$] | 0 |
| | Hepg2 | 1275 | 1274 [1273, 1275] | 1.001 [1.000, 1.001] | $1.6x10^{-1}$ [$7.1x10^{-3}$, $8.5x10^{-1}$] | 0 |
| | Hmec | 1365 | 1364 [1363, 1365] | 1.001 [1.000, 1.001] | $1.2x10^{-1}$ [$6.1x10^{-3}$, $7.3x10^{-1}$] | 1 |
| | Hsmm | 1395 | 1393 [1393, 1394] | 1.001 [1.000, 1.001] | $6.5x10^{-2}$ [$2.7x10^{-3}$, $5.1x10^{-1}$] | 10 |
| | Huvec | 1329 | 1327 [1327, 1328] | 1.001 [1.000, 1.002] | $1.2x10^{-2}$ [$2.0x10^{-4}$, $2.0x10^{-1}$] | 261 |
| | K562 | 1202 | 1202 [1201, 1203] | 1.000 [0.999, 1.001] | $6.4x10^{-1}$ [$1.2x10^{-1}$, $9.8x10^{-1}$] | 0 |
| | Nhek | 1353 | 1351 [1350, 1352] | 1.001 [1.001, 1.002] | $1.6x10^{-2}$ [$2.7x10^{-4}$, $2.4x10^{-1}$] | 202 |
| | Nhlf | 1341 | 1340 [1339, 1341] | 1.001 [1.000, 1.001] | $6.6x10^{-2}$ [$2.6x10^{-3}$, $5.3x10^{-1}$] | 13 |
| **Insulator** | Gm12878 | 1455 | 1455 [1454, 1456] | 1.000 [0.999, 1.001] | $6.7x10^{-1}$ [$1.8x10^{-1}$, $9.8x10^{-1}$] | 0 |
| | H1hesc | 1567 | 1566 [1565, 1567] | 1.001 [1.000, 1.001] | $4.9x10^{-1}$ [$6.0x10^{-2}$, $9.7x10^{-1}$] | 0 |
| | Hepg2 | 1504 | 1505 [1504, 1506] | 0.999 [0.998, 1.000] | $1.5x10^{-1}$ [$9.0x10^{-3}$, $7.8x10^{-1}$] | 0 |
| | Hmec | 1445 | 1445 [1444, 1447] | 1.000 [0.999, 1.000] | $5.0x10^{-1}$ [$7.0x10^{-2}$, $9.7x10^{-1}$] | 0 |
| | Hsmm | 1494 | 1494 [1493, 1496] | 1.000 [0.999, 1.001] | $5.0x10^{-1}$ [$6.3x10^{-2}$, $9.8x10^{-1}$] | 0 |
| | Huvec | 1481 | 1481 [1480, 1482] | 1.000 [0.999, 1.001] | $6.7x10^{-1}$ [$1.6x10^{-1}$, $9.9x10^{-1}$] | 0 |
| | K562 | 1497 | 1497 [1495, 1498] | 1.000 [0.999, 1.001] | $6.7x10^{-1}$ [$1.6x10^{-1}$, $9.8x10^{-1}$] | 0 |
| | Nhek | 1452 | 1452 [1451, 1453] | 1.000 [0.999, 1.001] | $6.4x10^{-1}$ [$1.3x10^{-1}$, $9.8x10^{-1}$] | 0 |
| | Nhlf | 1427 | 1428 [1427, 1429] | 1.000 [0.999, 1.000] | $5.7x10^{-1}$ [$8.7x10^{-2}$, $9.8x10^{-1}$] | 0 |
| **Transcription** | Gm12878 | 1023 | 1022 [1022, 1023] | 1.001 [1.000, 1.002] | $1.0x10^{-1}$ [$3.6x10^{-3}$, $7.0x10^{-1}$] | 2 |
| | H1hesc | 1143 | 1142 [1141, 1142] | 1.001 [1.000, 1.001] | $6.9x10^{-3}$ [$6.7x10^{-5}$, $1.5x10^{-1}$] | 599 |
| | Hepg2 | 1038 | 1038 [1037, 1039] | 1.000 [0.999, 1.001] | $2.8x10^{-1}$ [$1.7x10^{-2}$, $9.3x10^{-1}$] | 1 |
| | Hmec | 1083 | 1082 [1082, 1083] | 1.001 [1.000, 1.002] | $2.8x10^{-2}$ [$5.6x10^{-4}$, $3.2x10^{-1}$] | 79 |
| | Hsmm | 1102 | 1101 [1101, 1102] | 1.001 [1.000, 1.001] | $1.6x10^{-2}$ [$2.7x10^{-4}$, $2.4x10^{-1}$] | 172 |
| | Huvec | 1048 | 1047 [1046, 1048] | 1.001 [1.000, 1.002] | $6.3x10^{-2}$ [$1.7x10^{-3}$, $5.5x10^{-1}$] | 21 |
| | K562 | 1019 | 1018 [1017, 1019] | 1.001 [1.000, 1.001] | $1.4x10^{-1}$ [$5.2x10^{-3}$, $8.1x10^{-1}$] | 3 |
| | Nhek | 1063 | 1062 [1061, 1063] | 1.001 [1.000, 1.001] | $5.0x10^{-2}$ [$1.3x10^{-3}$, $4.7x10^{-1}$] | 33 |
| | Nhlf | 1069 | 1068 [1067, 1069] | 1.001 [1.000, 1.002] | $4.6x10^{-3}$ [$4.3x10^{-5}$, $1.1x10^{-1}$] | 798 |
| **Repressed** | Gm12878 | 1392 | 1391 [1390, 1392] | 1.001 [1.000, 1.002] | $1.8x10^{-1}$ [$1.0x10^{-2}$, $8.6x10^{-1}$] | 1 |
| | H1hesc | 1220 | 1219 [1218, 1220] | 1.001 [1.000, 1.002] | $1.8x10^{-1}$ [$1.2x10^{-2}$, $8.3x10^{-1}$] | 1 |
| | Hepg2 | 1503 | 1503 [1502, 1505] | 1.000 [0.999, 1.001] | $6.8x10^{-1}$ [$1.7x10^{-1}$, $9.9x10^{-1}$] | 0 |
| | Hmec | 1438 | 1438 [1437, 1440] | 1.000 [0.999, 1.001] | $5.1x10^{-1}$ [$6.5x10^{-2}$, $9.8x10^{-1}$] | 0 |
| | Hsmm | 1418 | 1418 [1417, 1419] | 1.000 [0.999, 1.001] | $6.6x10^{-1}$ [$1.4x10^{-1}$, $9.8x10^{-1}$] | 0 |
| | Huvec | 1367 | 1368 [1367, 1369] | 1.000 [0.999, 1.000] | $5.0x10^{-1}$ [$6.4x10^{-2}$, $9.7x10^{-1}$] | 0 |
| | K562 | 1473 | 1473 [1472, 1474] | 1.000 [0.999, 1.000] | $6.7x10^{-1}$ [$1.6x10^{-1}$, $9.8x10^{-1}$] | 0 |
| | Nhek | 1524 | 1524 [1523, 1526] | 0.999 [0.999, 1.001] | $3.7x10^{-1}$ [$3.6x10^{-2}$, $9.6x10^{-1}$] | 0 |
| | Nhlf | 1404 | 1404 [1402, 1405] | 1.000 [0.999, 1.001] | $6.6x10^{-1}$ [$1.5x10^{-1}$, $9.8x10^{-1}$] | 0 |
| **Heterochromatin** | Gm12878 | 1392 | 1391 [1390, 1391] | 1.001 [1.000, 1.001] | $8.9x10^{-4}$ [$6.2x10^{-6}$, $3.3x10^{-2}$] | 2574 |
| | H1hesc | 1370 | 1369 [1369, 1370] | 1.001 [1.000, 1.001] | $2.5x10^{-3}$ [$2.3x10^{-5}$, $6.5x10^{-2}$] | 1213 |
| | Hepg2 | 1387 | 1385 [1385, 1386] | 1.001 [1.000, 1.001] | $2.3x10^{-4}$ [$1.0x10^{-6}$, $1.2x10^{-2}$] | 4770 |
| | Hmec | 1375 | 1374 [1373, 1374] | 1.001 [1.000, 1.001] | $1.1x10^{-3}$ [$8.2x10^{-6}$, $3.6x10^{-2}$] | 2295 |
| | Hsmm | 1385 | 1384 [1384, 1385] | 1.001 [1.000, 1.001] | $3.8x10^{-3}$ [$4.6x10^{-5}$, $9.0x10^{-2}$] | 845 |
| | Huvec | 1382 | 1381 [1380, 1381] | 1.001 [1.000, 1.001] | $3.8x10^{-4}$ [$1.8x10^{-6}$, $1.8x10^{-2}$] | 3909 |
| | K562 | 1391 | 1390 [1389, 1390] | 1.001 [1.001, 1.001] | $1.1x10^{-4}$ [$4.3x10^{-7}$, $7.7x10^{-3}$] | 6007 |
| | Nhek | 1370 | 1369 [1369, 1370] | 1.001 [1.000, 1.001] | $4.3x10^{-4}$ [$2.4x10^{-6}$, $1.8x10^{-2}$] | 3725 |
| | Nhlf | 1379 | 1378 [1377, 1378] | 1.001 [1.000, 1.001] | $2.7x10^{-3}$ [$2.9x10^{-5}$, $7.0x10^{-2}$] | 1168 |

**S7 Table. VIKING vs LBC: known INDEL load comparison in different chromatin states (alleles per individual per 1Mb).** To annotate the number of variants in a state/cell type class as significantly different, we required at least 95% of the 10,000 subsets to have p-value ≤ $2x10^{-4}$ (Bonferroni corrected) and no overlap between the 95% CI for the LBC and VIKING median values; non-significant differences shown in grey.

| State | Cell Type | VIKING median | LBC 10k subsets median & 95%CI | VIKING/LBC ratio median & 95%CI | Wilcoxon rank sum test | |
|---|---|---|---|---|---|---|
| | | | | | $p$ : median & 95% CI | number of tests with $p \le 2x10^{-4}$ |
| **Promoter** | Gm12878 | 131.87 | 130.77 [130.57, 130.97] | 1.008 [1.007, 1.010] | $3.8x10^{-17}$ [$1.9x10^{-21}$, $3.0x10^{-13}$] | 10000 |
| | H1hesc | 134.92 | 133.53 [133.33, 133.72] | 1.010 [1.009, 1.012] | $1.3x10^{-25}$ [$1.1x10^{-30}$, $6.0x10^{-21}$] | 10000 |
| | Hepg2 | 130.44 | 129.17 [129.00, 129.37] | 1.010 [1.008, 1.011] | $5.4x10^{-22}$ [$9.0x10^{-27}$, $1.4x10^{-17}$] | 10000 |
| | Hmec | 138.03 | 136.22 [135.98, 136.43] | 1.013 [1.012, 1.015] | $7.1x10^{-30}$ [$5.0x10^{-35}$, $4.6x10^{-25}$] | 10000 |
| | Hsmm | 135.56 | 133.76 [133.57, 133.96] | 1.013 [1.012, 1.015] | $2.3x10^{-30}$ [$2.7x10^{-35}$, $1.3x10^{-25}$] | 10000 |
| | Huvec | 140.53 | 138.65 [138.45, 138.88] | 1.014 [1.012, 1.015] | $1.3x10^{-28}$ [$1.5x10^{-33}$, $7.4x10^{-24}$] | 10000 |
| | K562 | 135.28 | 134.10 [133.93, 134.26] | 1.009 [1.008, 1.010] | $1.3x10^{-16}$ [$1.2x10^{-20}$, $6.5x10^{-13}$] | 10000 |
| | Nhek | 139.76 | 138.24 [138.02, 138.43] | 1.011 [1.010, 1.013] | $7.3x10^{-26}$ [$7.2x10^{-31}$, $2.5x10^{-21}$] | 10000 |
| | Nhlf | 138.09 | 136.52 [136.31, 136.69] | 1.011 [1.010, 1.013] | $6.8x10^{-29}$ [$9.1x10^{-34}$, $3.2x10^{-24}$] | 10000 |
| **Enhancer** | Gm12878 | 120.80 | 120.01 [119.90, 120.14] | 1.007 [1.005, 1.007] | $9.4x10^{-19}$ [$5.4x10^{-23}$, $8.5x10^{-15}$] | 10000 |
| | H1hesc | 125.50 | 124.70 [124.57, 124.85] | 1.006 [1.005, 1.008] | $2.4x10^{-21}$ [$4.5x10^{-26}$, $4.7x10^{-17}$] | 10000 |
| | Hepg2 | 116.88 | 116.11 [116.00, 116.24] | 1.007 [1.005, 1.008] | $9.7x10^{-18}$ [$3.9x10^{-22}$, $1.1x10^{-13}$] | 10000 |
| | Hmec | 122.97 | 122.27 [122.12, 122.38] | 1.006 [1.005, 1.007] | $1.4x10^{-23}$ [$2.4x10^{-28}$, $2.5x10^{-19}$] | 10000 |
| | Hsmm | 126.19 | 125.34 [125.21, 125.41] | 1.007 [1.006, 1.008] | $7.7x10^{-25}$ [$3.1x10^{-29}$, $1.0x10^{-20}$] | 10000 |
| | Huvec | 128.53 | 127.73 [127.62, 127.88] | 1.006 [1.005, 1.007] | $2.0x10^{-20}$ [$3.6x10^{-25}$, $2.9x10^{-16}$] | 10000 |
| | K562 | 112.07 | 111.42 [111.29, 111.53] | 1.006 [1.005, 1.007] | $2.1x10^{-17}$ [$6.5x10^{-22}$, $2.1x10^{-13}$] | 10000 |
| | Nhek | 124.47 | 123.64 [123.51, 123.77] | 1.007 [1.006, 1.008] | $4.3x10^{-24}$ [$5.6x10^{-29}$, $1.1x10^{-19}$] | 10000 |
| | Nhlf | 126.22 | 125.39 [125.28, 125.52] | 1.007 [1.006, 1.008] | $9.1x10^{-24}$ [$1.5x10^{-28}$, $2.6x10^{-19}$] | 10000 |
| **Insulator** | Gm12878 | 138.49 | 137.68 [137.42, 137.95] | 1.006 [1.004, 1.008] | $9.1x10^{-4}$ [$7.3x10^{-6}$, $3.0x10^{-2}$] | 2454 |
| | H1hesc | 136.59 | 135.99 [135.76, 136.22] | 1.004 [1.003, 1.006] | $7.6x10^{-5}$ [$2.6x10^{-7}$, $5.7x10^{-3}$] | 6533 |
| | Hepg2 | 149.21 | 148.61 [148.26, 148.95] | 1.004 [1.002, 1.006] | $4.5x10^{-3}$ [$5.7x10^{-5}$, $9.2x10^{-2}$] | 741 |
| | Hmec | 145.93 | 145.45 [145.05, 145.77] | 1.003 [1.001, 1.006] | $6.0x10^{-3}$ [$7.5x10^{-5}$, $1.2x10^{-1}$] | 579 |
| | Hsmm | 146.92 | 146.79 [146.54, 147.12] | 1.001 [0.999, 1.003] | $3.3x10^{-1}$ [$3.1x10^{-2}$, $9.5x10^{-1}$] | 0 |
| | Huvec | 145.04 | 144.77 [144.44, 145.04] | 1.002 [1.000, 1.004] | $9.7x10^{-2}$ [$5.1x10^{-3}$, $5.8x10^{-1}$] | 2 |
| | K562 | 133.94 | 133.16 [132.90, 133.48] | 1.006 [1.004, 1.008] | $4.4x10^{-6}$ [$6.8x10^{-9}$, $6.4x10^{-4}$] | 9281 |
| | Nhek | 143.94 | 143.37 [143.06, 143.63] | 1.004 [1.002, 1.006] | $1.8x10^{-4}$ [$9.3x10^{-7}$, $1.0x10^{-2}$] | 5209 |
| | Nhlf | 128.74 | 128.17 [127.94, 128.40] | 1.004 [1.003, 1.006] | $1.8x10^{-5}$ [$5.1x10^{-8}$, $1.5x10^{-3}$] | 8390 |
| **Transcription** | Gm12878 | 118.45 | 117.62 [117.48, 117.71] | 1.007 [1.006, 1.008] | $3.5x10^{-29}$ [$2.1x10^{-34}$, $2.3x10^{-24}$] | 10000 |
| | H1hesc | 123.56 | 122.68 [122.59, 122.80] | 1.007 [1.006, 1.008] | $4.5x10^{-41}$ [$2.8x10^{-46}$, $4.8x10^{-36}$] | 10000 |
| | Hepg2 | 118.89 | 118.07 [117.95, 118.15] | 1.007 [1.006, 1.008] | $1.1x10^{-33}$ [$5.0x10^{-39}$, $1.3x10^{-28}$] | 10000 |
| | Hmec | 121.93 | 121.12 [121.03, 121.23] | 1.007 [1.006, 1.007] | $7.3x10^{-32}$ [$4.3x10^{-37}$, $5.9x10^{-27}$] | 10000 |
| | Hsmm | 122.49 | 121.64 [121.55, 121.74] | 1.007 [1.006, 1.008] | $4.8x10^{-37}$ [$1.8x10^{-42}$, $7.0x10^{-32}$] | 10000 |
| | Huvec | 120.42 | 119.57 [119.46, 119.67] | 1.007 [1.006, 1.008] | $5.3x10^{-32}$ [$4.2x10^{-37}$, $4.4x10^{-27}$] | 10000 |
| | K562 | 118.77 | 117.90 [117.80, 118.02] | 1.007 [1.006, 1.008] | $2.2x10^{-33}$ [$7.2x10^{-39}$, $2.4x10^{-28}$] | 10000 |
| | Nhek | 120.05 | 119.18 [119.10, 119.29] | 1.007 [1.006, 1.008] | $1.2x10^{-34}$ [$5.6x10^{-40}$, $1.7x10^{-29}$] | 10000 |
| | Nhlf | 121.62 | 120.76 [120.64, 120.84] | 1.007 [1.006, 1.008] | $2.9x10^{-34}$ [$2.1x10^{-39}$, $2.8x10^{-29}$] | 10000 |
| **Repressed** | Gm12878 | 123.70 | 122.97 [122.83, 123.11] | 1.006 [1.005, 1.007] | $7.0x10^{-13}$ [$9.5x10^{-17}$, $1.7x10^{-9}$] | 10000 |
| | H1hesc | 117.27 | 116.31 [116.13, 116.58] | 1.008 [1.006, 1.010] | $9.9x10^{-10}$ [$4.5x10^{-13}$, $8.7x10^{-7}$] | 10000 |
| | Hepg2 | 128.09 | 127.42 [127.25, 127.58] | 1.005 [1.004, 1.007] | $3.6x10^{-9}$ [$1.9x10^{-12}$, $2.5x10^{-6}$] | 9998 |
| | Hmec | 122.80 | 122.19 [122.00, 122.37] | 1.005 [1.003, 1.006] | $2.3x10^{-6}$ [$3.2x10^{-9}$, $4.4x10^{-4}$] | 9495 |
| | Hsmm | 126.57 | 126.11 [125.97, 126.26] | 1.004 [1.002, 1.005] | $1.6x10^{-7}$ [$1.3x10^{-10}$, $5.0x10^{-5}$] | 9936 |
| | Huvec | 125.25 | 124.82 [124.67, 124.95] | 1.003 [1.002, 1.005] | $2.5x10^{-6}$ [$5.7x10^{-9}$, $3.6x10^{-4}$] | 9547 |
| | K562 | 127.10 | 126.46 [126.34, 126.60] | 1.005 [1.004, 1.006] | $8.4x10^{-15}$ [$4.2x10^{-19}$, $4.7x10^{-11}$] | 10000 |
| | Nhek | 128.13 | 127.45 [127.33, 127.60] | 1.005 [1.004, 1.006] | $2.5x10^{-11}$ [$6.1x10^{-15}$, $4.1x10^{-8}$] | 10000 |
| | Nhlf | 120.58 | 119.88 [119.78, 120.06] | 1.006 [1.004, 1.007] | $1.5x10^{-12}$ [$4.4x10^{-16}$, $2.1x10^{-9}$] | 10000 |
| **Heterochromatin** | Gm12878 | 137.82 | 137.11 [137.05, 137.17] | 1.005 [1.005, 1.006] | $1.2x10^{-42}$ [$9.3x10^{-48}$, $1.3x10^{-37}$] | 10000 |
| | H1hesc | 136.72 | 136.10 [136.03, 136.15] | 1.005 [1.004, 1.005] | $1.2x10^{-39}$ [$8.0x10^{-45}$, $1.5x10^{-24}$] | 10000 |
| | Hepg2 | 138.21 | 137.50 [137.43, 137.57] | 1.005 [1.005, 1.006] | $3.8x10^{-42}$ [$2.7x10^{-47}$, $3.5x10^{-37}$] | 10000 |
| | Hmec | 137.03 | 136.35 [136.28, 136.42] | 1.005 [1.005, 1.006] | $7.6x10^{-43}$ [$5.3x10^{-48}$, $1.1x10^{-37}$] | 10000 |
| | Hsmm | 137.74 | 137.08 [136.99, 137.15] | 1.005 [1.004, 1.005] | $1.0x10^{-39}$ [$9.4x10^{-45}$, $8.7x10^{-35}$] | 10000 |
| | Huvec | 137.09 | 136.39 [136.32, 136.46] | 1.005 [1.005, 1.006] | $3.7x10^{-44}$ [$3.3x10^{-49}$, $3.6x10^{-39}$] | 10000 |
| | K562 | 139.31 | 138.58 [138.53, 138.66] | 1.005 [1.005, 1.006] | $2.1x10^{-41}$ [$2.8x10^{-46}$, $1.4x10^{-36}$] | 10000 |
| | Nhek | 137.52 | 136.85 [136.77, 136.91] | 1.005 [1.004, 1.006] | $3.6x10^{-41}$ [$2.9x10^{-46}$, $4.5x10^{-36}$] | 10000 |
| | Nhlf | 138.56 | 137.41 [137.32, 137.46] | 1.005 [1.004, 1.005] | $1.6x10^{-40}$ [$1.2x10^{-45}$, $1.8x10^{-35}$] | 10000 |

**S8 Table. Comparison of the ROH regions discovered in VIKING and LBC.** To annotate a difference as significantly different, we required at least 95% of the 10,000 subsets to have p-value ≤ 0.0125 (Bonferroni corrected) and no overlap between the 95% CI for the LBC and VIKING median values. The ROHs used for the analysis are filtered to exclude ROH regions with poor SNP density (see Fig 8 in S1 File).

| | | **VIKING** | **LBC (10k sub-samples)** | | | **VIKING/LBC ratio** | | | **Wilcoxon rank sum test** | | | |
|---|---|---|---|---|---|---|---|---|---|---|---|---|
| | | median | median | 95% LO | 95% HI | median | 95% LO | 95% HI | median | 95% LO | 95% HI | number of tests with p ≤ 0.0125 |
| **Intermediate** | **number** | 135 | 142 | 140 | 143 | 0.951 | 0.944 | 0.964 | $9.4 \times 10^{-12}$ | $3.3 \times 10^{-15}$ | $1.0 \times 10^{-8}$ | 10000 |
| | **length** | 97.1 Mb | 102.4 Mb | 101.2 Mb | 103.9Mb | 0.948 | 0.935 | 0.959 | $8.5 \times 10^{-12}$ | $2.5 \times 10^{-15}$ | $1.0 \times 10^{-8}$ | 10000 |
| **Long** | **number** | 3 | 1 | 1 | 2 | 3.000 | 1.500 | 3.000 | $2.7 \times 10^{-22}$ | $3.8 \times 10^{-26}$ | $1.1 \times 10^{-18}$ | 10000 |
| | **length** | 9.6 Mb | 4.2 Mb | 3.3 Mb | 4.5 Mb | 2.307 | 2.155 | 2.934 | $1.7 \times 10^{-31}$ | $3.1 \times 10^{-35}$ | $7.3 \times 10^{-28}$ | 10000 |

**S9 Table. The 13 exonic variants found to be significantly enriched in VIKING compared to gnomADg (Fisher's Exact Test) in genes predicted to be largely intolerant to variation and for which a strong evidence of gene-trait association ($p \leq 5x10^{-8}$) is reported in the GWAS Catalog (v1.0.1).** From the gnomAD dataset we report the MAF for the population with the maximum MAF for the variant; gnomADg is WGS data (n = 15,496) and gnomADe is WES data (v2.1.1, n = 125,748). The p-value for the VIKING vs gnomADg enrichment for a variant is calculated using Fisher's Exact Test.

| chr | pos | id | ref | alt | VIKING MAF | LBC MAF | gnomADg MAF (max) | gnomADe MAF (max) | Gene | Effect | VIKING/gnomADg enrichment p-value | gene-trait correlation (p-value ≤ $5x10^{-8}$ in GWAS Catalog v1.0.1) |
|---|---|---|---|---|---|---|---|---|---|---|---|---|
| 19 | 49,232,226 | rs2287922 | G | A | 0.625 | 0.565 | 0.4815 | 0.4918 | RASIP1 | missense_variant | $4.0x10^{-19}$ | Retinal vascular calibre, Urinary metabolites (H-NMR features), Inflammatory skin disease, C-reactive protein levels or LDL-cholesterol levels (pleiotropy), Mean platelet volume |
| 17 | 48,649,340 | rs768796872 | G | A | 0.013 | 0 | 0 | 0.0001651 | CACNA1G | missense_variant | $2.3x10^{-16}$ | Alzheimer's disease (cognitive decline) |
| 5 | 98,215,264 | rs370760984 | C | T | 0.011 | 0.0008651 | 0.0003332 | 0.0002320 | CHD1 | missense_variant | $7.3x10^{-16}$ | Eosinophil percentage of white cells, Eosinophil counts, Eosinophil percentage of granulocytes, Neutrophil percentage of granulocytes |
| 3 | 10,400,565 | rs774434270 | C | T | 0.005576 | 0.0004325 | 0 | 0.00003266 | ATP2B2 | missense_variant | $4.0x10^{-13}$ | Autism spectrum disorder or schizophrenia |
| 19 | 42,753,837 | rs368169058 | G | A | 0.013 | 0 | 0 | 0.0002846 | ERF | missense_variant | $7.6x10^{-13}$ | Monocyte count |
| 14 | 79,423,644 | rs139593796 | G | A | 0.011 | 0 | 0.0001332 | 0.0001055 | NRXN3 | missense_variant | $3.7x10^{-12}$ | Waist circumference, Body mass index, Obesity, Waist-hip ratio, Hip circumference, Cerebrospinal fluid biomarker levels, Initial pursuit acceleration |
| 20 | 47,247,325 | rs138500849 | A | T | 0.022 | 0.002595 | 0.002045 | 0.001959 | PREX1 | missense_variant | $4.2x10^{-11}$ | Colorectal cancer, Multiple myeloma, Diastolic blood pressure, Intelligence (multi-trait analysis) |
| 17 | 48,703,924 | rs762245146 | C | G | 0.011 | 0 | 0.00006696 | 0.0001834 | CACNA1G | missense_variant | $1.4x10^{-10}$ | Alzheimer's disease (cognitive decline) |
| 22 | 40,042,737 | rs201769752 | T | C | 0.011 | 0.001298 | 0.0002004 | 0.0008119 | CACNA1I | missense_variant | $2.1x10^{-10}$ | IgG glycosylation, Schizophrenia, Autism spectrum disorder or schizophrenia, Cognitive ability (multi-trait analysis), Intelligence (multi-trait analysis) |
| 10 | 78,647,084 | . | G | C | 0.007435 | 0.0004325 | 0 | 0 | KCNMA1 | missense_variant | $1.6x10^{-9}$ | Obesity, Hypospadias, Myopia |
| 10 | 78,708,961 | rs148156399 | A | G | 0.007435 | 0 | 0 | 0.0000088 | KCNMA1 | missense_variant | $8.8x10^{-9}$ | Obesity, Hypospadias, Myopia |
| 3 | 47,162,897 | rs114719990 | T | C | 0.013 | 0.0008651 | 0.001399 | 0.001446 | SETD2 | missense_variant | $3.4x10^{-7}$ | HDL cholesterol levels, Macrophage inflammatory protein 1b levels, Monocyte count, Lymphocyte counts, Lymphocyte percentage of white cells, Monocyte percentage of white cells, Granulocyte percentage of myeloid white cells, White blood cell count, Platelet distribution width |
| 18 | 52,928,743 | rs147445499 | G | A | 0.011 | 0.0004325 | 0.0008661 | 0.001184 | TCF4 | missense_variant | $4.7x10^{-7}$ | Schizophrenia, Fuchs's corneal dystrophy, Sclerosing cholangitis and ulcerative colitis (combined), Autism spectrum disorder or schizophrenia, Neuroticism |

**S10 Table. The 6 rare variants (gnomADg MAF < 0.05, Shetland MAF ≤ 0.1) predicted to be eQTLs (GTEx v7, qval ≤ 0.05) and to affect the expression of 6 distinct genes.** From the gnomAD dataset we report the MAF for the population with the maximum MAF for the variant; gnomADg is WGS data (n = 15,496). The p-value for the VIKING vs gnomADg enrichment for a variant is calculated using Fisher's Exact Test.

| chr | pos | id | ref | alt | VIKING MAF | LBC MAF | gnomADg MAF (max) | Mapped Gene | Tissue | VIKING/gnomADg enrichment p-value | gene-trait correlation (p-value ≤ $5x10^{-8}$ in GWAS Catalog v1.0.1) |
|---|---|---|---|---|---|---|---|---|---|---|---|
| 1 | 154,909,169 | rs17356361 | C | T | 0.097 | 0.043 | 0.034 | PMVK | Esophagus_Mucosa | $2.5x10^{-9}$ | Atrial fibrillation, Lung function (FEV1/FVC), Parkinson's disease |
| 11 | 65,379,532 | rs138504384 | GTC | G | 0.067 | 0.031 | 0.021 | ZNHIT2 | Cells_Transformed_fibroblasts | $6.9x10^{-9}$ | |
| 9 | 136,341,547 | rs117965396 | C | T | 0.061 | 0.026 | 0.019 | CACFD1 | Esophagus_Muscularis<br>Muscle_Skeletal | $1.4x10^{-8}$ | |
| 1 | 155,066,403 | rs138207102 | C | T | 0.052 | 0.022 | 0.015 | CHRNB2 | Brain_Putamen_basal_ganglia | $3.3x10^{-8}$ | Smoking behaviour (cigarettes smoked / day) |
| 10 | 81,839,199 | rs150170834 | G | A | 0.046 | 0.019 | 0.013 | FAM213A | Liver | $3.4x10^{-8}$ | Lung function (FEV1/FVC), Heel bone mineral density, Blood protein levels, Systolic blood pressure, Intraocular pressure, Sunburns |
| 1 | 46,935,356 | rs150548439 | A | C | 0.063 | 0.033 | 0.022 | LRRC41 | Artery_Aorta | $2.0x10^{-7}$ | Menopause (age at onset) |

**S11 Table. Mean and standard deviation of proportion of sites with particular number of MAF alleles in the VIKING and LBC cohorts.** The analysis is based on high-quality SNPs/INDELs discovered in the callable regions of the 22 autosomal chromosomes in the two cohorts of unrelated individuals, split to known variants (present in gnomAD at any frequency) and ultra-rare variants (not found in any gnomAD population). All sites with missing genotype(s) were excluded. The means and standard deviations for each frequency were computed based on subsampling the two cohorts to 50 individuals each repeated 100 times (also see Fig 6 in S1 File).

| | | number MAF alleles in known SNPs | | | | | | | number MAF alleles in ultra-rare SNPs | | |
|---|---|---|---|---|---|---|---|---|---|---|---|
| | | **1** | **2** | **3** | **4-5** | **6-10** | **11-20** | **21-50** | **1** | **2** | **3+** |
| **VIK** | mean | 0.190 | 0.090 | 0.060 | 0.077 | 0.120 | 0.154 | 0.320 | 0.881 | 0.093 | 0.027 |
| | s.d. | $5.6x10^{-17}$ | $8.4x10^{-17}$ | $2.0x10^{-3}$ | $4.5x10^{-3}$ | $2.0x10^{-16}$ | $4.9x10^{-3}$ | $2.2x10^{-16}$ | $6.8x10^{-3}$ | $5.6x10^{-3}$ | $4.6x10^{-3}$ |
| **LBC** | mean | 0.220 | 0.080 | 0.050 | 0.070 | 0.115 | 0.150 | 0.310 | 0.980 | 0.010 | 0.010 |
| | s.d. | $1.1x10^{-16}$ | $5.6x10^{-17}$ | $9.8x10^{-17}$ | $8.4x10^{-17}$ | $5.0x10^{-3}$ | $2.5x10^{-16}$ | $5.6x10^{-16}$ | $4.5x10^{-16}$ | $7.0x10^{-18}$ | $7.0x10^{-18}$ |

| | | number MAF alleles in known INDELs | | | | | | | number MAF alleles in ultra-rare INDELs | | |
|---|---|---|---|---|---|---|---|---|---|---|---|
| | | **1** | **2** | **3** | **4-5** | **6-10** | **11-20** | **21-50** | **1** | **2** | **3+** |
| **VIK** | mean | 0.186 | 0.089 | 0.057 | 0.079 | 0.120 | 0.160 | 0.320 | 0.863 | 0.096 | 0.040 |
| | s.d. | $4.9x10^{-3}$ | $3.0x10^{-3}$ | $4.6x10^{-3}$ | $2.6x10^{-3}$ | $2.0x10^{-16}$ | $1.1x10^{-16}$ | $2.2x10^{-16}$ | $7.1x10^{-3}$ | $6.0x10^{-3}$ | $1.4x10^{-3}$ |
| **LBC** | mean | 0.211 | 0.080 | 0.050 | 0.070 | 0.120 | 0.150 | 0.310 | 0.970 | 0.010 | 0.020 |
| | s.d. | $2.9x10^{-3}$ | $5.6x10^{-17}$ | $9.8x10^{-17}$ | $8.4x10^{-17}$ | $2.0x10^{-16}$ | $2.5x10^{-16}$ | $5.6x10^{-16}$ | $7.8x10^{-16}$ | $1.7x10^{-3}$ | $1.4x10^{-17}$ |

**S12 Table. Functional VIKING variants enriched in genes largely intolerant to variation.** Applied filtering criteria are denoted with 'yes'.

| | **total number enriched variants** | **filtering criteria** | | | **number filtered enriched variants** |
|---|---|---|---|---|---|
| | | **CADD >= 20** | **pLI >= 0.8** | **missense z-score >= 3** | |
| **SNPs** | | | | | |
| stop_gained | 557 | | yes | | 23 |
| splice_acceptor_variant | 9 | | yes | | 1 |
| splice_donor_variant | 27 | | yes | | 3 |
| start_lost | 46 | | yes | | 5 |
| stop_lost | 23 | | yes | | 2 |
| missense_variant | 26054 | yes | | yes | 1165 |
| **INDELs** | | | | | |
| frameshift_variant | 749 | | yes | | 50 |
| inframe_insertion | 133 | yes | yes | yes | 0 |
| inframe_deletion | 396 | yes | yes | yes | 12 |
